# Nearest Neighbour Interactions between Amino Acid Residues in Short Peptides and Coil Libraries

**DOI:** 10.64898/2026.01.19.700493

**Authors:** Reinhard Schweitzer-Stenner

## Abstract

Intrinsically disordered proteins (IDP) or proteins with intrinsically disordered regions (IDR) perform a plethora of functions mostly in a cellular environment. As unfolded proteins, IDPs can adopt molten globule or coil ensembles of conformations. Regarding the latter the question arises whether they are describable as a self-avoiding random coil. Locally, this requires that amino acid residues sample the entire sterically allowed region of the Ramachandran plot with very similar probabilities and independent on the conformational dynamics of their neighbours. However, various lines of experimental and bioinformatic evidence suggest a more restricted, side chain and nearest neighbor dependent conformational space for individual residues. Over the last 25 years short peptides and coil libraries were employed to determine conformational propensities of amino acid residues in unfolded states. The question arises whether conformational ensembles obtained from these two sources are comparable. In this paper, a variety of metrics were used to compare Ramachandran plots of a limited number of GXYG peptides (X,Y: guest residues) with XY dimers in the coil library of Ting et al.(PLOS 6, e1000763, 2010). The results reveal major differences between corresponding plots, which might in part due to the fact that solely the influence of one of the two neighbours of a given residue is probed by the above coil library while averages were taken over the respective opposite neighbours. The presented results suggest that coil libraries alone might not be a sufficient tool for determining the characteristics of statistical coils of IDPS and IDRs alike.

## Introduction

It is now well established that intrinsically disordered proteins (IDPs) and folded proteins with intrinsically disordered regions (IDRs) perform a plethora of functions mostly in a cellular environment. Their very existence shatters the conventional dogma that stipulate the need for a stable folded protein for its function.(1–6) Estimates of the fraction of biologically relevant IDPs vary in the literature.(2,6) A recent study reported 30% of eukaryotic proteins to be at least partially disordered.(7) In many cases IDPs or IDRs of otherwise folded proteins are involved in molecular recognition processes that involve disorder → order transitions.(8–13) In less frequent cases transitions into fuzzy or even another disordered state occur. (10,14,15) Protein disorder was found to dominate in proteins associated with signal transduction, cell-cycle regulation, gene expression and chaperone action.(5,6,16,17) Besides being of pivotal importance for biological processes IDPs can also malfunction by self-assembly into fibrillar structures that have been implicated in several neurological diseases.(2,18–20). Aggregation of IDPs is also involved in the droplet formation due to liquid-liquid phase separations.(21)

Generally, IDPs exhibit a dynamic behavior that allows them to exist as an ensemble of energetically similar conformations under physiological conditions. Several lines of evidence suggest that in structural terms IDPs can be treated as unfolded proteins so that their conformational ensembles can be classified either as collapsed molten or pre-molten globules (if e.g. water is a poor solvent) or as self-avoiding random coils (if water is a good solvent).(3) A more nuanced picture emerged from NMR studies using residual dipolar coupling and secondary chemical shifts, which revealed the occurrence of local residual structure in ensembles which otherwise appear dominantly coiled. The propensities of proteins for these global states generally depend on their amino acid composition and in particular on the respective charge distributions.(22–25)

This paper focusses on the local structural characteristics of totally unfolded, coiled proteins. Traditionally, they are described by invoking the random coil concept. Conceptually, both the ideal Flory random coil model and the self-avoiding random walk model are based on the assumption that a polymer can be described as a freely jointed chain of subunits.(26,27) In the case of polypeptides/proteins this would require a nearly unhindered rotation around the NC^α^ and C^α^C’ bonds that connect the peptide groups. Apparently, this idealization is at variance with the observation that steric hindrance and electrostatic interactions between residues restrict the corresponding dihedral backbone angles ⍰ and ψ.(28–31) The sterically allowed regions of the Ramachandran plot of individual amino acid residues are believed to be very similar with the sole exception of glycine (more conformationally flexible due to the lack of chirality) and proline (more restricted due to the pyrrolidine ring). Hence, individual sequences of polypeptides and proteins would not matter much for coiled proteins with regard to experimentally accessible parameters like the radius of gyration and end to end distances. Indeed, the radii of gyration *R*_*g*_ of proteins and IDPs in denaturing solvents were found to be describable by the classical scaling law (N: number of residues) with scaling exponents clustering around 0.6,(32,33) the expectation value for a self-avoiding random walk in a good solvent.(26,34)

While totally unfolded (denatured) proteins behave globally like self-avoiding random coils in most cases, experimental, computational and bioinformatic evidence suggest a more restricted and side chain dependent conformational sampling of the Ramachandran space than assumed even by restricted random coil models. The latter assume that the Ramachandran plots of all residues besides proline and glycine are describable by a very broad and shallow minimum of the Gibbs energy landscape in the upper left quadrant of the Ramachandran plot so that all conformations in the region between ϕ=−160° and −50° as well as between ψ=180° and 90° are practically isoenergetic. Traditionally, it was called the β-strand region.(28,29) A less extended region between ϕ=−160° and −50° as well as ψ=20° and −70° exhibits similar minimal Gibbs energies as the broad region in the upper left quadrant. It encompasses all types of right-handed helical and type I/II β-turn conformation generally sampled by their i+2 residue. Much smaller regions of higher Gibbs energy are associated with left-handed helical and other turn structures. In Figure 1, the local restricted random coil distribution is approximated by a mixture of nearly isoenergetic Gaussian basins the parameters of which are listed in Table S1 (taken from ref.(35)). Contrary to this view, spectroscopic experiments on short host-guest residue type peptides such as blocked dipeptides, NH_3_^+^-GXG-COOH and Ac-GGXGG-NH_2_, peptides (X: guest residue) revealed that e.g. alanine intrinsically exhibits a high preference for a conformation called polyproline II (pPII, ϕ > −100°), while valine slightly favors β-strand (ϕ ≤ −100°) over pPII (cf. ref. (36) and references cited therein). Protonated aspartic acid samples turn-supporting conformations to a significant extent. Similar though not identical conformational preferences have been obtained in so-called coil libraries. They were constructed from the dihedral angles adopted in segments of folded proteins, which do not depict one of the well-known regular structures (ref. (37) and references cited therein). They are considered as representative of coiled proteins because it is assumed that any non-local interactions and steric constraints are averaged out if one builds Ramachandran distributions with a large number of proteins that possess a significant fraction of loops. Computationally, an increasing number of molecular dynamics (MD) simulations using modified Amber, CHARMM or OPLS forcefield and TIP water models is capable of reproducing the high pPII propensity of alanine to some extent,(38–44) but a reproduction of the side chain specificity of Ramachandran distributions has only recently been achieved, though only on a qualitative level.(45)

**Figure 1.**
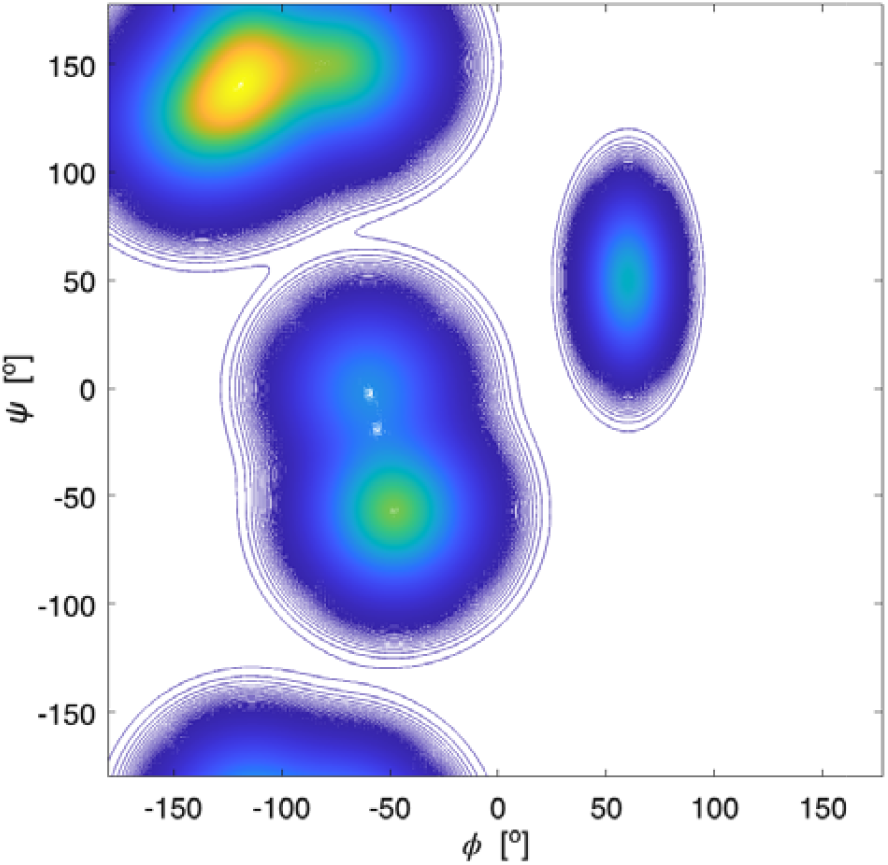
Ramachandran plot of a (local) restricted random coil distribution of conformational ensembles. The respective parameters have been taken from ref.(35). They are listed in Table S1.

The discovery of residue specific propensities changed our picture of coiled peptides and proteins in that it suggests that, locally, unfolded peptides and proteins exhibit a statistical coil rather than a (restricted) random coil behaviour. In the statistical coil case, the broad basin in the upper left quadrant splits into two much more confined minima with side chain dependent free energies, which in many cases are not identical. Conformations in the right-handed helical/β-turn I/II region are generally of significantly higher Gibbs energy. Hence, the free energy landscape is describable by a statistical canonical set of Boltzmann factors. The (restricted) random coil case thus appears as a special case of a statistical coil. Generally, statistical coil ensembles carry significant less conformational entropy then a classical restricted random coil distribution.(35,46)

Even though the above conformational restrictions of the Ramachandran space seem to have a limited influence on global properties of coils such as the radius of gyration or end to end distance,(33,46,47) their assessment is of utmmost importance for producing statistical coil reference systems based on which local transient residual structures in unfolded proteins and IDPs can be identified by nuclear magnetic resonance spectroscopy (ref.(48) and references cited therein). Information about the local properties of coiled peptides and proteins also allow for an accurate estimation of configurational entropies.(49,50) Unfortunately, the task of identifying the coiled state of proteins is complicated by nearest and potentially even second nearest neighbour interactions (NNIs) between residues (ref.(37) and references cited therein). The occurrence of such interactions is well established in general, but the underlying mechanisms are not well understood. It is still not entirely clear to what extent NNIs involve correlated conformational motions between residue, in which case they would depend on the conformation adopted by the neighbours. Several lines of evidence suggest that such structural correlations exist, in which case the isolated pair hypothesis (IPH), another pillar of the random coil model, would no longer be valid (ref.(37) and references cited therein). As a consequence, configurational entropies and free solvation energies would no longer be additive.

Carrying out a complete quantitative analysis of NNIs is difficult because they are side chain dependent. Hence, even if one focuses on NNIs and thus ignores second neighbour effects, one would have to analyse e.g. 20^3^ GXYZG unblocked penta-peptides or blocked tetra-peptides in solution in order to cover all possible combinations. The number reduces to 400 GXYG tetra-peptides (or blocked tripeptides), if one assumes that the influence of down- and upstream neighbours are additive. Several lines of evidence suggest that this is too optimistic an assumption.(51–54) Spectroscopic investigations of tetra- and pentapeptides in water clearly revealed significant nearest neighbour interactions, which most likely involve correlation effects.(53–57) Even though the number of investigated peptides has been significant, they cannot be considered as fully representative for the entire manifold of side chain combinations. Therefore, coil libraries have emerged as a preferred source for NNI analysis. A coil library of proteins is a database of backbone and side chain conformations, which generally excludes helical and sheet sequences of proteins. For some libraries, turns have also been excluded. However, the use of thus restricted libraries makes it difficult to obtain statistically significant Ramachandran plots of individual residues for all possible up- and downstream neighbours (cf. the Ramachandran plots that can be constructed at (58)). As a remedy, Ramachandran plots for individual amino acid residues have been constructed by averaging over all neighbours.(59–62) Such a strategy would be valid if individual NNIs with different neighbours were very similar, at variance with the results of the above spectroscopic studies on tetra- and pentapeptides. (52–54,57)

In this paper, I explore the influence of residue averaging by a comparative analysis of Ramachandran distributions of amino acid residues in tetra-peptides and in the coil-library of Dunbrack and coworkers.(63) Due to my best knowledge, the latter is the only one that provides statistically significant information about the specific influence of individual neighbours on Ramachandran distributions of all twenty amino acid residues. The authors restricted averaging to either the right (downstream) or the left (upstream) neighbours, respectively. In other words, for all-XY pairs, the Ramachandran plots of X were reported for all Y neighbours, while the influence of the left neighbours were averaged, whereas plots for Y were produced for all X neighbours with average over the right neighbours (XY-all pairs). Ting et al. used a mathematical algorithm to construct complete Ramachandran plots with a resolution of 5° for the ϕ and ψ coordinate from the obtained data points taken from a restricted coil library (no helices, strands and turns). A limited comparison of these coil library Ramachandran plots with tetra-peptide distributions had been undertaken by Schweitzer-Stenner and Toal based on the use of calculated Hellinger distances (*vide infra*).(56) The current paper presents the results of a by far more comprehensive analysis, which compares respective population of mesostates as well as experimental scalar NMR-based J-coupling constants of tetrapeptides with coupling constants calculated for respective coil library distributions of Ting et al.. Revised Hellinger distance calculations are augmented by comparisons of configurational Gibbs entropies for individual residues. To my best knowledge, such a set of different metrics have not yet been applied to any coil library. The results of this analysis reveal that the diversity of individual NNIs is not sufficiently represented by coil library distributions and that averaging over neighbours underestimates the influence of neighbours.

### Theoretical background

The basics of the so-called Gaussian model implemented for the analysis of J-coupling constants as well Raman, IR, and vibrational dichroism amide I’ profiles of tri-, tetra- and pentapeptides have been described in several earlier papers of my research group.(64,65) Briefly, observables *O* (i.e. scalar J-coupling constants and amide I’ profiles in the above spectra) were calculated as ensemble average of a statistical coil distribution:

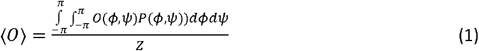

where O represents the considered spectroscopic observable (vide supra) and *Z* denotes the partition sum of the ensemble. In the past, the distribution function *P*(*ϕ,ψ*) was constructed as a superposition of two-dimensional Gaussian functions:

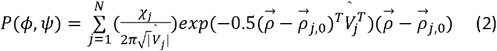

where the index *j* represents the *j*^th^ conformation summed over *N* different sub-distributions with statistical weights (mole fractions) *χ*_*j*._ The position vector 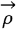 represent a coordinate pair (*ϕ,ψ*), 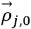 denotes the position the center of the sub-distribution. 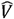 is a 2 x 2 matrix, the diagonal of which contains of the standard deviations of the Gaussian sub-distributions with respect to ϕ and ψ. The off-diagonal elements account for correlations between the two coordinates, which in the case of unequal standard deviations for ϕ and ψ lead to a rotation of the axes of the respective ellipse. In the current study, the Gaussian model was solely used for non-linear least square fits of eq. (1) to a set of experimental scalar J-coupling constants of GXYG peptides (i.e. the ϕ-dependent ^3^J(H^N^H^Cα^), ^3^J(H^N^C’), ^3^J(H^Cα^C’), ^3^J(H^N^C^β^) and the ψ-dependent ^1^J(NC_α_), which were carried out with a Matlab program (X and Y represent the two guest residues). For the fitting procedure, the statistical weights as well as the positions of the Gaussian sub-distributions were used as free parameters in an iterative procedure described by Milorey et al.(54) The standard deviations of the distributions were slightly varied between 10° and 20° for a fine-tuning of the fits. Scalar coupling constants were calculated by using the Karplus parameters in ref.(66) for the ^3^J constants and in ref.(67) for the ^1^J constant. Ramachandran plots were calculated with a resolution of 2° for ϕ and ψ. In order to account for the amide I’ band profiles, conformations of the entire peptide rather than residue conformations have to be used. The final output of the above fitting procedure was used as input for a truncated model applied to reproduce the amide I’ profiles. This reduction was necessary to reduce the computational cost of the conformational sampling. A total of 21 pairs of ϕ, ψ angles represents an ellipse around the center pPII and β-strand troughs, respectively. Hence, we substitute the two-dimensional Gaussian distributions by ellipses (not cuboids, as stated in ref.(54)) with axis widths of 2*Δ*⍰_*j*_ and 2*Δψ*_*j*_ in ⍰ and *ψ* direction, where Δ⍰_*j*_ and Δ*ψ*_*j*_ were set equal to the standard deviations of the respective Gaussian distributions. The height of the ellipses is given by the statistical weights of the conformations divided by the total number of representing data points (i.e. 21). Representative plots of these distributions are shown in Figure S1. In addition, the less dominant conformations (right-handed helical, inverse γ-turn, asx-turn, etc.) were represented by a single conformation positioned at the respective center of the Gaussian sub-distribution with the statistical weights obtained from the fits of the Gaussian model to J-coupling constants. Several checks revealed that the truncated model reproduced the coupling constants produced by the fitting to the J-coupling data sufficiently well. If the input distribution reproduced the measured amide I’ profiles, the analysis was considered complete. If the measured and calculated amide I’ profiles differed significantly, the above parameters were slightly modified to reduce the deviations. The newly obtained set of parameters was used a starting point for refitting the J-coupling constants. The procedure was repeated until convergence was obtained. This fitting procedure, which was first implemented by Milorey et al. for their conformational analysis of GRRG and GRRRG peptides,(54) was found to be more reliable and efficient than the procedure that our group implemented in the earlier study of GXYG peptides.(57) Therein, we had used the identical sub-spaces of the Ramachandran plot for manual simulations of the entire data set of J-coupling constants and amide I profiles. The resolution of the conformational sampling was 6°, much lower than the 2° used for the current Gaussian analysis of tetra- and penta-peptides.(54) A change of the positions of Gaussian sub-distributions could move part of them outside of the area probed by the respective simulations. Therefore, in order to put all the available tetra-peptide data on an equal footing, I revisited the data of Toal et al.(57) with the above procedure of Milorey et al.(54) It should be noted that the same exciton coupling model was used in both studies to link residue conformations to amide I’ band profiles. Since vibrational excitonic coupling between amide modes in different residues depends on the dihedral angles between the coupled oscillators the intensity distributions of amide I’ are excellent indicators of backbone conformations. Details of the utilized excitonic model are described in earlier papers.(65)

Different metrics were used to compare the Ramachandran plots of amino acid residues flanked by different neighbours in the investigated tetra-peptides and in the coil library of Ting et al.(63) The Hellinger distance H is a measure of overlap between different distributions. It is defined as follows:

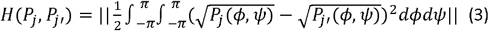

where *P*_*j*_(*ϕ,ψ*) and *P*_*j’*_(*ϕ,ψ*) denote the Ramachandran probability density distributions of the *j*^th^ residue in tri- and tetrapeptides, respectively. The Hellinger distance can vary between 0 (total agreement) and 1 (completely distinct).^†^ To interpret these numbers, earlier introduced criteria were used.(56) If the Hellinger value lies in the region below 0.1, the corresponding distributions are considered as very similar. If 0.1 < H ≤ 0.25 and 0.25 < H ≤ 0.4, the distributions are moderately similar and moderately dissimilar, respectively. H values larger than 0.4 indicate very dissimilar distributions. The Hellinger distance responds very sensitively to changes of the positions of individual basins (i.e. Gaussian distributions). Sole changes of statistical weights have a lesser influence on H.

An alternative metric is the Gibbs (or Shannon) entropy which for an individual residue is written as:

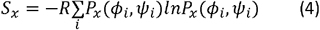

where x is just an indicator of the considered residue and R is the gas constant. The numerical value of *S*_*x*_ depends on the mesh size of the Ramachandran plot (2° for the investigated peptides, 5° for the coil library Ramachandran plots).(49,50) Therefore, we just used the differences between the entropies of X with different neighbours as a suitable metric for characterizing NNIs.(35)

Calculating the Gibbs entropies of residues does not provide a complete picture, if NNIs involve correlations between conformational motions of neighbours. In such a case only the conformational distributions of entire peptides and even of unfolded and disordered proteins have to be taken into account, because thermodynamic parameters are no longer additive. In order to model such a scenario e.g. for our tetrapeptides, one has to use the Ramachandran plots of GXG peptides as reference and model the NNI induced differences in terms of interaction free energies.(46,52,55,68) The statistical weight of a distinct conformation of a tetra-peptide GXYG where X has adopted conformation L and Y the conformation K is thus written as:

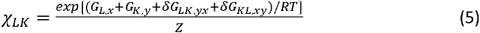

where G_L,x_ and G_K,y_ are the Gibbs free energies of the states L and K of the residues X and Y in GXG/GYG tripeptides. Hence, glycine as neighbour is used as a reference point, for which one would expect minimal nearest neighbour interactions. The interaction free energy between the two conformations in X and Y are denoted as *δG*_*LK,yx*_ and *δG*_*KL,xy*_. The latter are obtained by reproducing the differences between the statistical weights of the conformations in GXG/GYG and GxyG.(52,55) Z is the partition sum of the system. It can conveniently be calculated by employing Zimm-Bragg like transfer matrices. Details can be found in ref.(55) The configurational Gibbs entropy of a XY ‘dimer’ can now be written as:

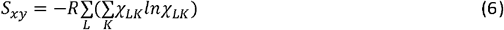

where *χ*_*LK*_ is the normalized statistical weight (mole fraction) of dimers adopting the conformations *L* and *K*. Ramachandran plots of peptide and coil library residues can be compared by calculating the respective population of mesostates. Figure S2 exhibits a graphic representation of the mesostates, which were used earlier for the comparison of the Gaussian model with the results of molecular dynamics calculations.(44) Each mesostate is associated with a specific secondary structure. It should be noted that the classical β-strand region is subdivided into areas associated with parallel (pβ), antiparallel (aβ) and transitional β-strand (βt). While pβ and aβ are canonical, βt is a more recent addition introduced by Tran et al.(69) More coarse-grained mesostate models partition this area only into a pPII and a β-strand region. The higher resolution of our model allows for a more nuanced picture of Ramachandran distributions. Since the same mesostates are used for all systems investigated, populations depend on the statistical weight of the respective Gaussian sub-distribution, its width and its position. The population of mesostates are just calculated as mole fraction by integrating the population density over its area in the Ramachandran plot.

## Results

This section is organized as follows. First, the experimental data reported for GXYG peptides by Toal et al.(57) are revisited using the protocol described in the preceding section. Thus, obtained conformational propensities (statistical weights) and basin positions (of Gaussian sub-distributions) are combined with the results of the more recent tetra-peptide studies of Milorey et al.(53,54) Second, respective Ramachandran plots of GXYG and coil library distributions are compared on a qualitative, visual level. In a third step such comparisons are carried out on a more quantitative level by using calculated J-coupling constants, mesostate populations, Hellinger distances and configurational entropies as metrics.

### Conformational propensities and basin positions

In a first step the experimental data which Toal et al. reported for a series of cationic GXYG peptides in aqueous solution were revisited by employing the protocol of Milorey et al. (*vide supra*). The choice of cationic peptides was motivated by the need to obtain sufficiently high intensities of amide proton signals. The influence of terminal charges on the conformational distributions of central residues in AAA, GAG and GVG have been found to be negligible, while it is moderate for GDG.(43,53) These finding suggests that even in tripeptides hydration of terminal groups cause a significant shielding and thus a reduction of electrostatic interactions. The results of the GXYG analysis are presented in Tables S1 and S2 as well as in Figures S3-S7. Table S1 compares the experimental J coupling constants with the ones resulting from the fitting procedure described in the preceding section. The corresponding calculated coupling constants reported by Toal et al. are listed for comparison. Table S2 lists the obtained statistical weights and peak coordinates of the Gaussian sub-distributions for the two non-terminal X,Y residues of the investigated GXYG. Corresponding values reported by Toal et al. are listed in parenthesis for comparison. Figure S3 shows a representative set of fits to the amide I’ profiles in the IR, isotropic Raman, anisotropic Raman and VCD spectrum of the investigated GXYG peptides. The fits to corresponding band profiles were obtained with the same spectral parameters (i.e. positions and halfwidths of three Gaussian bands related to transitions into the first excited excitonic state of coupled oscillators) used by Toal et al. It should be noted that owing to the influence of nearest neighbour and non-nearest neighbour coupling, amide I’ profiles reflect the conformational states of the peptide rather than the ones of individual residues, which are specifically probed by the utilized J-coupling constants. Figures S4-S7 depict the statistical weight of pPII, β-strand and of all helical and turn-supporting conformations for a specific residue in different GXYG peptides (e.g. of alanine in GD**A**G, GS**A**G, G**A**LG, G**A**VG, G**A**FG and GF**A**G, target residue in bold). In addition to the revised statistical weights of the GXYG peptides investigated by Toal et al.(57), recent results obtained for GAFG, GFAG, GDFG, GFDG, GDDG and GKKG have been added to the graphs.(52–54) Representative Ramachandran plots of GXYG peptides are shown below, where they are compared with corresponding coil library plots.

Similarities and discrepancies between the new and old parameter values of the Gaussian model are briefly delineated in the Supporting Information. Here, it can just be stated that the main observations of Toal et al. remain valid, namely the rather pronounced decrease of the pPII propensity of alanine in the presence of unlike neighbours with the exception of phenylalanine (average decrease by 29%). Concomitantly, the β-strand propensity is significantly increased. Aspartic acid ranks highest in terms of the nearest neighbour sensitivity of statistical weights, but in this case pPII increases by an average of 50% in the presence of unlike neighbours. Surprisingly, the pPII fraction of phenylalanine increases on average by 30%. On the contrary, the respective pPII sensitivities of serine, leucine and lysine are comparatively low (average decreases between 12% and 15%), again in line with Toal et al. Aspartic acid and serine itself as neighbour causes significant changes and rather position dependent changes. Like neighbours (i.e. in G**K**KG, GK**K**G, G**D**DG, GD**D**G, G**R**RG, GR**R**G and GL**L**G produce much more pronounced changes). (52–54)

In order to facilitate a thermodynamic understanding of the plots in Figures S4-S7, Figure S8 depicts the Gibbs free energy difference between pPII and β-strand basins for different populations of the upper left quadrant of the Ramachandran plot. A comparison of these plots with the mole fractions listed in Table S2 reveal that a significant number of NNI induced population changes are associated with Gibbs energy changes of several kJ/mol. For instance, serine and valine destabilize the pPII conformation of alanine, by Gibbs free energy changes of ca. 4 and 3 kJ/mol respectively.

### Comparison of peptide and coil library Ramachandran plots

Figures 2 and 3 show the Ramachandran plots of a selected number of investigated residue pairs in GXYG and all-XY/XY-all segments of the coil library (all indicates the averaging over all 20 neighbours). The plots of the remaining tetra peptides and coil library segments investigated in this paper are shown in Figures S9-S16. In addition to the tetra-peptide plots, GXG plots for the respective target residues are shown, e.g. GAG for G**A**VG and all-**A**G for all-**A**Y, where bold characters indicate the target residue.

**Figure 2:**
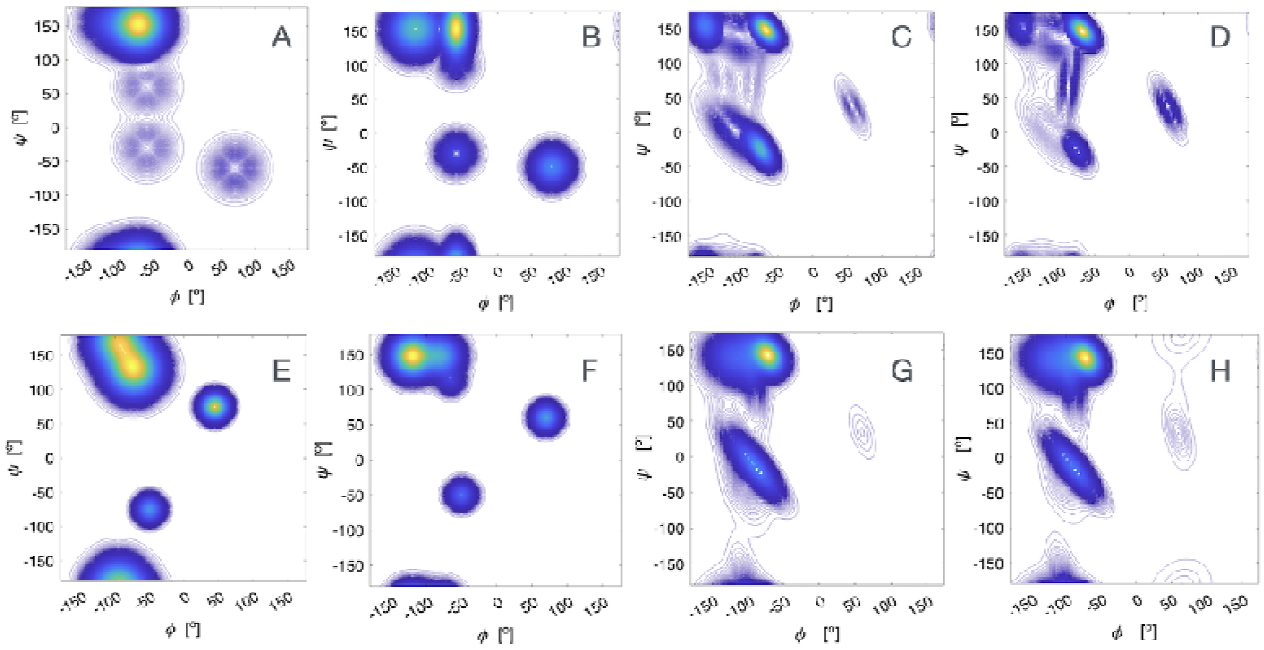
Ramachandran plot of alanine residues in cationic GAG (A), cationic G**A**VG (B), in the coil library segments all-**A**G and all-**A**V (C and D), in cationic GLG (E), cationic GL**L**G (F) and in the coil library segments GL-all and LL-all (G and H). The tripeptide Ramachandran plots were created with the Gaussian parameters reported in ref.(70) The plots for G**A**VG and GLLG were produced for this study by an analysis of experimental data reported by Toal et al.(57), as described in the main text. The plots for A and L in the above coil library segments were created with files obtained from the website of the Dunbrack group with the kind permission of Dr. Dunbrack.(63)

**Figure 3:**
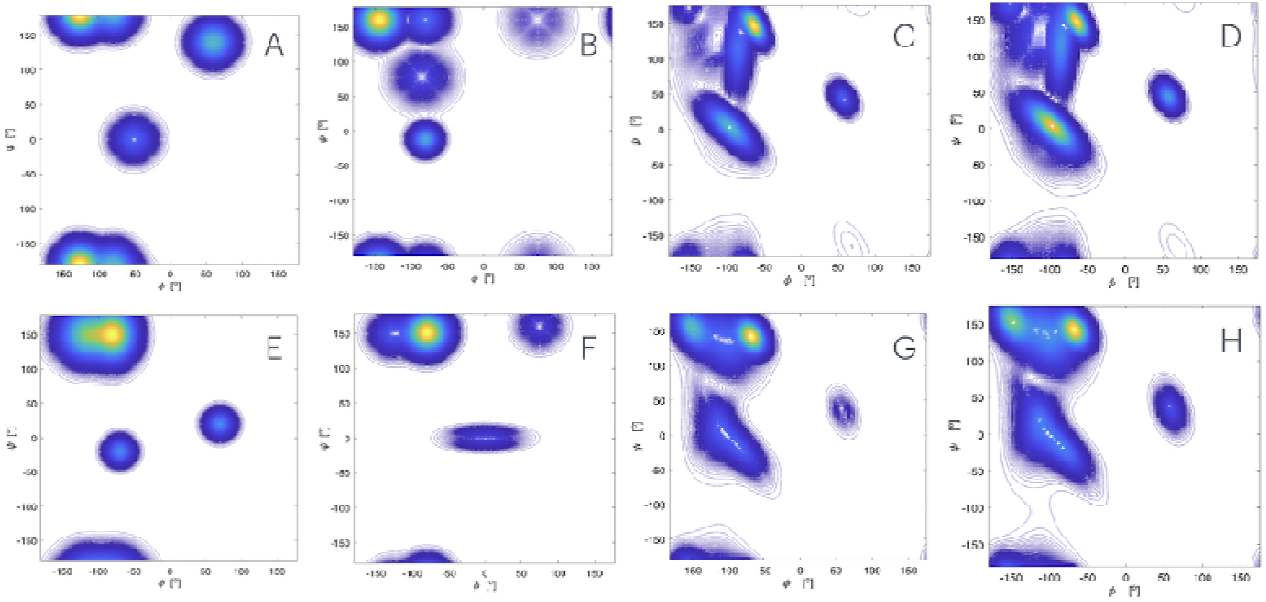
Ramachandran plot of the aspartic acid residue in cationic GDG (A), cationic GDDG (B), in the coil library segment G**D**-all and D**D**-all (C and D), in cationic GFG (E), cationic GDFG (F) and in the coil library segment G**F**-all and D**F**-all (F). The tripeptide Ramachandran plots were created with the Gaussian parameters reported in refs.(70,71), with slight modifications of the GDG mole fractions.(53) The plots for GD**D**G and GD**F**G were created by using the respective Gaussian parameters reported in earlier papers.(53) The plots for D and F in the above coil library segments were created with files obtained from the website of the Dunbrack group with the kind permission of Dr. Dunbrack.(63)

A first superficial look at all the displayed Ramachandran plot reveals the following qualitative picture. In line with the above discussed statistical weight values of investigated tetrapeptides, nearest neighbour interactions vary the pP2 <-> β equilibrium, though to a different extent (vide supra). Moreover, for quite a substantial number of peptides, they increase the distance between the two basins along the ϕ-coordinate (GS**A**G, GD**L**G, GS**L**G, G**L**LG, GV**L**G, G**S**AG, G**S**KG, and G**S**VG). These changes do not correlate with population changes of the pPII and β-strand. Turn supporting conformations, which are significant in the Ramachandran plots of D and S containing peptides, are significantly varied by their neighbours. Generally, interactions between like residues in homo-dimers (G**D**DG, GD**D**G, G**R**RG and GR**R**G) are more pronounced than the ones in hetero-dimers. The coil dimer Ramachandran plots do not convey a similar story. While they clearly differ for different amino acid residues (just compare the all-XG plots), NNI induced changes seem to be rather limited. Most interestingly and also surprisingly, the data suggest a dominant population of the pPII basin for all residues with the exception of valine, for which the β-strand is more significantly populated, while the pPII basin has moved into a region which is generally associated with the i^th^ residue of type II β-turns. The Ramachandran plots of most coil library dimers exhibit a significant distance between pPII and β-basins with the latter positioned outside of the canonical positions of residues in antiparallel and parallel β-sheets. This suggests a non-Arrhenius Kramers-type structure of the transition state.

The above description of peptide and coil Ramachandran is pretty much based on visual inspection. In what follows I use metrics introduced in the preceding section to provide a more quantitative comparison of Ramachandran plots.

### Comparison of J-coupling constants

While the above discussed Ramachandran plots suggest significant differences between conformational distributions obtained from a Gaussian analysis of short peptides and those derived from coil library data of Ting et al., these discrepancies themselves do not per se show that peptide and coil library distributions are different, since the Ramachandran plots obtained for the former are model dependent. Fortunately, the Ramachandran plots of the coil library contain enough data points to allow for the calculation of ensemble averaged J-coupling constants. Figures 4 displays the experimental and calculated J-coupling constants for a representative series of tetrapeptides (experimental and Gaussian analysis) and related XY dimers of the Ting et al. library.

**Figure 4:**
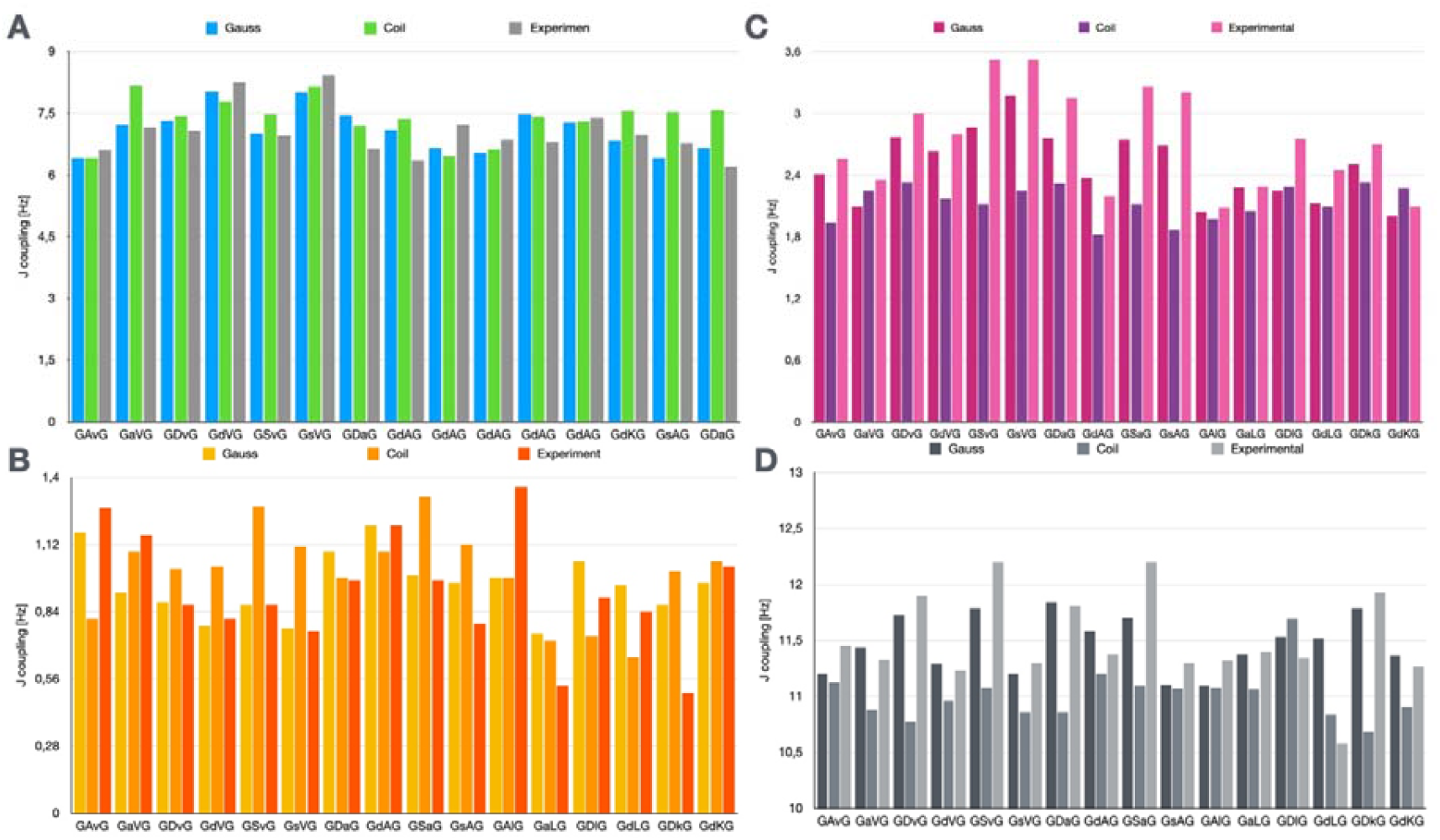
Comparison of experimental, Gaussian modelling and coil library J-coupling constants of the indicated residues (by capital letters) with different neighbours in GXYG peptides (lower case for better identification). Lower case letters indicate the neighbours. Experimental values were taken from refs.(52,57) A: ^3^J(H^N^ H^Cα^), B: ^3^J(H^N^ C’), C: ^3^J(H ^Cα^ C’) and D: ^1^J(NC_α_)

Generally, the data in Figure 4 reveal that Gaussian coupling parameter values are closer to experimental ones than coil library-based values. For ^3^J(H^N^,H^Cα^), the differences between coil library and experimental values are most notable for **A**V, S**A** and **D**A pairs. Discrepancies are even more pronounced and numerous for the other three coupling constants of **A**V, **S**A, **D**A, **D**V, **S**V, **D**L, **D**K and S**V**. Coil values for the ψ-dependent ^1^J((NC_α_) are systematically lower than the experimental GXYG values which, as discussed further below, is indicative of a larger population of right-handed helix like conformations. A visual inspection of the Ramachandran plots in Figures 1,2, and S6-S16 suggests that differences between corresponding coil library distributions are by far less significant than the ones between the Gaussian plots. This impression is put on a more quantitative basis by calculating the average experimental J coupling constants for the GXYG peptides with a common residue (e.g. G**F**AG, GA**F**G, G**F**DG and GD**F**G) and the corresponding coil library dimers (e.g. all-**F**A, A**F**-all, all-**F**D and D**F**-all). The respective values are listed in Table 1. First, a comparison with corresponding GXG values reveal residue specific differences for both data sets. Second, while differences between average coupling constant values are noteworthy, they are not as pronounced as the differences between the standard deviations, which for most of the considered sequences are larger for the experimental than for the coil library-based coupling constants. This result strongly suggests that compared with GXYG the side chain specificity of nearest neighbour interactions is significantly reduced in the Ting et al. coil library. It should be noted here that the term ‘standard deviation’ does not indicate that the averaged data set constitute a statistical ensemble.

**Table 1.**
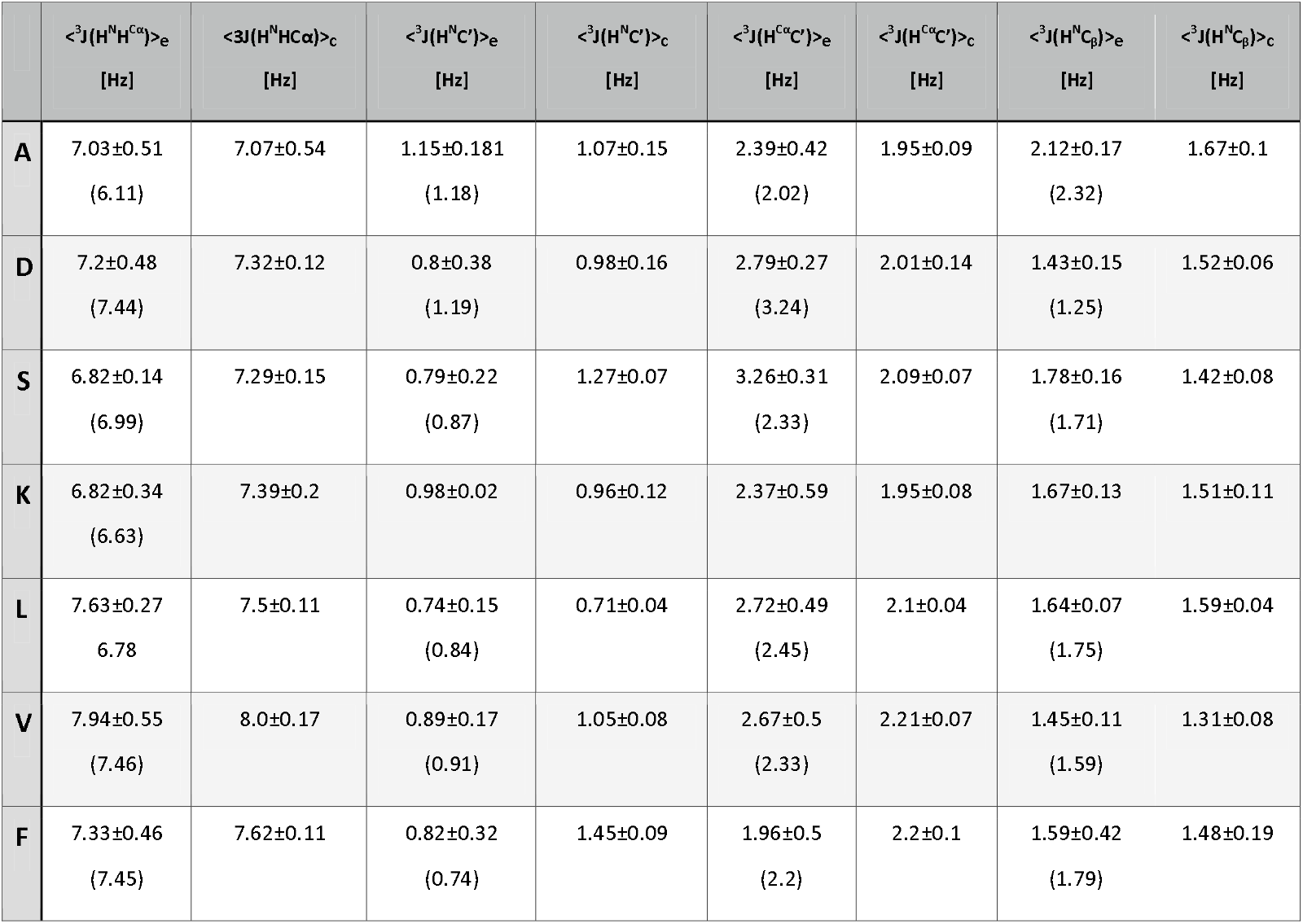
Listing of average experimental (e) (52–54,57) and calculated J-coupling constants of the Ting et al. coil library (c).(63) The average was taken over the entire set of residue dimers in GXYG and coil library segments containing the indicated residue type. Coupling parameters of these residues in GXG peptides are listed in parenthesis for comparison.(70–72)

### Comparison of mesostate populations

In the Gaussian model conformational propensities are expressed in terms of statistical weights (mole fractions) of the utilized Gaussian sub-distributions. Within the context of the model this makes sense thermodynamically, since these Gaussian distributions correspond to a classical harmonic approximation of the respective potential energy minimum. However, this parametrization does not allow a quantitative comparison with Ramachandran plots from molecular dynamics simulations and coil libraries. As shown earlier for the use of the former, calculating the populations of mesostates (Figure S2) provides a handy solution.(44)

The mesostate populations of the investigated residues in peptides and the Ting et al. coil library are shown in Figure S17. The populations of right-handed helices and β-turn I/IIi+2 conformations as well as the ones of antiparallel and parallel β-strand mesostates have been added up. For alanine, pPII dominates all distributions. For GAVG, the pPII population is significantly reduced, but it is still slightly larger than the combined β-strand confirmations, while the respective mole fractions are practically identical (Figure S4). The pPII mesostate of aspartic acid is comparatively weakly populated in GDG. Both, tetra-peptide and coil library data indicate that nearest neighbour interactions significantly increase the pPII population at the expense of asx-turns and βap. pPII is dominantly populated in the distributions of lysine and leucine; some exceptions notwithstanding (GSKG, GVKG, GLLG, GALG and GVLG). Most of the serine distributions exhibit comparable sampling of pPII and βap while most of the valine distributions are dominated by βt and βap (exceptions are DV-all, all-VL and KV-all). The sampling of the combined mesostate of right-handed helices and β-turn I/IIi+2 in the coil library distributions is significantly more pronounced than it is in the investigated peptides. Discrepancies between mesostate populations and the statistical weights displayed in Figures S4-S7 result from the fact that the Gaussian sub-distributions overlap and may contribute to more than one mesostate. This is certainly true for many pPII and β-strand Gaussians with relatively high ψ-values, which overlap with the ε-mesostate. Generally, pPII sub-distributions may contribute to the βt mesostate. Since some ψ-values of residues decrease in the presence of non-glycine neighbours, the pPII sampling can increase even if the statistical weight of the sub-distribution decreases. Alternatively, a narrowing of e.g. the pPII sub-distribution reduces the overlap with other mesostates thus increasing the population of its own mesostate.

Figure 5 compare the changes of mesostate populations in the investigated tetra-peptides and corresponding coil library sequences caused by upstream and downstream neighbours. For the calculation of these changes, GXG peptides were used as a reference for the tetra-peptides, while all-XG and G**Y**-all were used for all-XY and XY-all.

**Figure 5.**
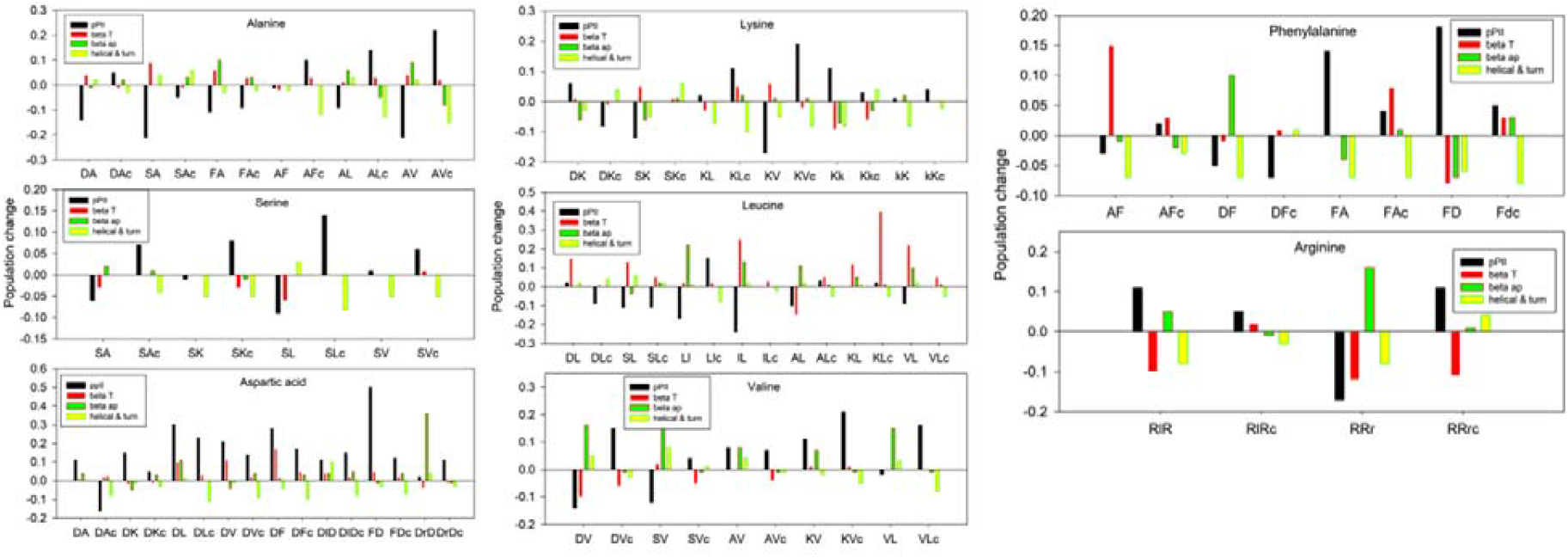
Plot of differences between the population of selected mesostates in GXYG and GXG (GYG) peptides and between populations in all-XY and all-XG as well as between XY-all and GY-all coil library segments. Note that the populations of mesostates associated with the parallel and antiparallel β-sheet structures as well as with helical and turn conformations have been added up, respectively.

The bar diagrams in Figure 5 reveal a complex picture, from which I distil only the most significant differences and similarities between peptide and coil distributions. For alanine in tetra-peptides all pPII populations decrease, in particular in the presence of serine and valine. This observation is in line with the behaviour of the corresponding Gaussian statistical weights (Figure S4), but it is in contrast to the pPII changes in the coil library plots of AF-all, AL-all, and all-AV. Similar discrepancies exist at least partially for the series of serine, lysine and valine. However, for aspartic acid and phenylalanine, pPII changes occur in the same direction. The changes obtained for aspartic acid are peculiar, in that nearly all pPII changes are positive, with all-DA as sole exception. For phenylalanine, the positive pPII changes are pronounced for FA and FD. In the coil library, the changes move in same direction, but to a much lesser extent.

As one can infer from Figure S17, the combined population of the right-handed helical and type II/I β-turn mesostates is much more pronounced in nearly all coil library Ramachandran plots even though it is generally reduced by nearest neighbour interactions with of non-glycine neighbours.

It should be noted that the displayed changes do not add up to zero for the following reasons. First, the Ramachandran plots of GxG peptides exhibit substantial sampling of the so-called ε-region, which lies right above the lower boundary of the lower left quadrant. The population of this region is generally reduced in tetra-peptides. Second, the Ramachandran plots of coil library residues are more inhomogeneous than the Gaussian plots. While they display relatively sharp maxima at the basin positions, the transition regions are more populated in the coil library distributions which are not covered by the mesostates in Figure S2.

Taken together, a comparison of conformational mesostate occupations in GXYG and in coil libraries reveal major differences. Overall, the nearest neighbour induced changes in the later seem to be less pronounced. Interestingly, however, they are also sometimes qualitatively different. This notion particularly applies to pPII the population of which is enhanced by aliphatic neighbours in coil libraries (V and L), whereas the same type of neighbour decreases the pPII population in GXYG. Right-handed helical and type I/II β-turn i+2 conformations is generally more populated in the Ting et al. library.

### Comparison of Hellinger distances and configurational entropies

It should be noted that observed changes of statistical weights convey only a part of the picture. As one can infer from Table S2, noticeable changes of the ϕ-position and to a more limited extend of the ψ-position were obtained for all investigated series, including the leucine and serine series, for which the propensity changes are moderate. As a consequence, related J-coupling constants are different (Table S1). To illustrate this point, let us look on the propensities and ^3^J(H^N^ H^Cα^) value of some leucine residues: The statistical weights for pPII of GA**L**G, G**L**LG, GV**L**G and GD**L**G are 0.42, 0.43, 0.4 and 0.49. Apparently, they are very similar. The first three values are identical within their statistical uncertainty, which generally does not exceed ± 0.5.(64) The corresponding ^3^J(H^N^ H^Cα^) values are 7.4, 7.62, 8.15 and 6.78 Hz. These variations lie clearly outside of the statistical error limits (< ±0.2 Hz). As shown below, changes of basin positions lead to rather large values of the Hellinger distance.

Eqs. (3) was used to calculate the Hellinger distance between the Ramachandran plots of GXG (GYG) and GXYG/GXYG for all investigated amino acid residue targets. In addition, Hellinger distances between plots of all-XY/XY-all and all-XG/GY-all were calculated. As clearly indicated by the bar plots in Figure 6 the Hellinger values obtained for the Ramachandran plots of the tetra-peptides exceed by far those of the Ting et al. library. With the exception of the serine series, all Hellinger values of tetra-peptide residues indicate a moderate or large dissimilarity between corresponding Ramachandran plots of tetra- and tripeptides. On the contrary, the overwhelming majority of the values obtained for the coil library sequences lie in the moderate similarity region. ^‡^ As indicated above, Hellinger distances are very sensitive to changes of basin positions and respond only moderately to changes of basin populations. Thus, the data plotted in Figure 6 suggest minimal changes of basins in the coil library, in line with what one would infer from a visual inspection of Figures 1, 2 and S9-S16. Altogether, the plots in Figure 6 clearly suggest that nearest neighbour induced changes are much less pronounced in the Ting et al. library than in the investigated GXYG peptides.

**Figure 6:**
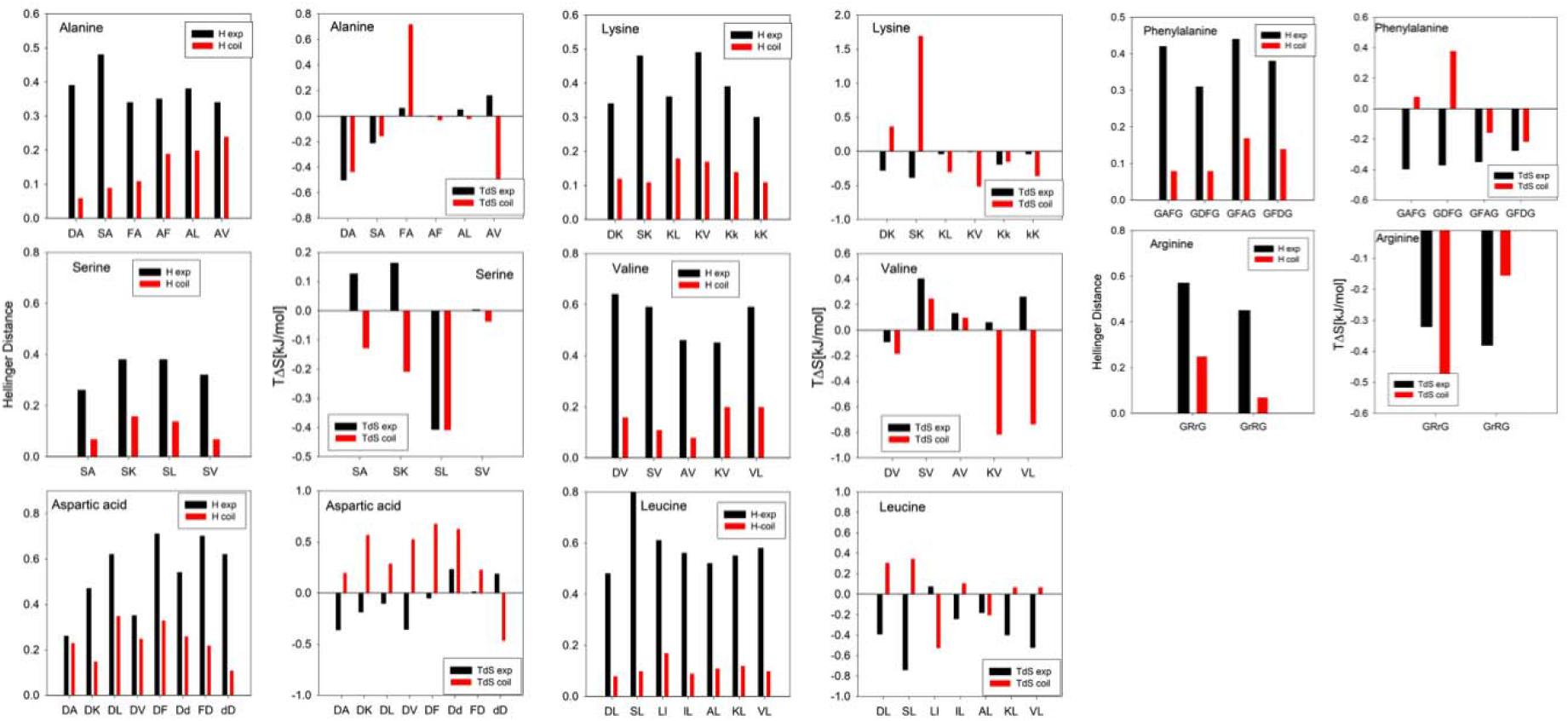
Hellinger distances and configurational entropy contribution to Gibbs free energy changes at room temperature due to nearest neighbour interactions for the indicated amino acid residues in GXYG peptides (black) and corresponding all-XY and XY-all segments of the Ting et al. library (red).

Since nearest neighbour interactions affect Ramachandran distributions, the configurational entropy changes as well. The corresponding changes of the Gibbs energy contribution are also plotted in Figure 6. Note that the depicted values were calculated for the indicated residue, not for the entire dimeric XY segments. Negative values for a distinct residue X(Y) imply that its Ramachandran plots is less entropic in the presence of the neighbour Y. The displayed TΔS values suggest a by far more complex picture than the Hellinger distances. While entropy changes in the tetra-peptides are dominant for some residues, they are rather small for others (F**A, A**V, **S**A, **D**F, F**D, K**L, **K**V, **K**K, **K**K and **L**L. Even more surprising is the observation that entropy changes of residues in GXYG and coil library dimers carry opposite signs for all serine neighbours and nearly all aspartic acid neighbours (S**L** and **D**D being an exception). Opposite behavior is also observed for e.g. K**V, V**L, S**K**, and D**F**. These findings suggest that relatively short side chains with hydrogen bonding capacity such as aspartic acid and serine behave differently as neighbours and as targets in short peptides and in coil libraries. For most GXYG peptides, nearest neighbour interactions decrease the entropic contribution to the Gibbs energy of their neighbours by several hundreds of Joule/mol.

Thus, these data suggest that a coil state of a proteins might carry less configurational entropy than assumed based on the (local) random coil model. However, the mixture of entropy increases and decreases obtained for the Ting et al. coil library does not allow for such a clear assessment. A more detailed discussion of this issue is given in the subsequent paragraph.

## Discussion

Nearly thirty years ago an analysis of ^3^J(H^N^ H^Cα^) coupling constants of unfolded proteins lead to the conclusion that Flory’s isolated pair hypothesis is not a valid supposition.(73) Subsequently, the influence of nearest and non-nearest neighbours on the conformational propensity was mostly inferred from the analysis of coil libraries (46,49,50,60,62,63,68,74,75) and from computational work on model peptides(69,75–77). The general view that emerged from these analyses suggested that aliphatic and aromatic neighbours decrease a residue’s pPII propensity. For alanine, Avbelji and Baldwin attributed this to conformational dependent changes of the solvation free energy.(76) Computational work of the Sosnick group revealed that the observed nearest neighbour interactions involve correlations between the dynamics of interacting residues, which implies that these interactions depend on the adopted conformation of the respective neighbours.(75) This is an important information since a violation of the isolated pair hypothesis requires such correlation effects. A conformational detuning that depends solely on the side chain properties of neighbours would not eliminate the additivity of conformational and solvation free energies and would therefore not lead to a violation of the isolated pair hypothesis.

Experimentally, the issue of nearest neighbour interactions was early recognized by the NMR community. Chemical shifts and to a lesser extent ^3^J(H^N^ H^Cα^) coupling constants in short glycine-based host-guest peptides were considered as ideal references for a random coil conformation. However, the context dependence of these parameters in proteins led to the conclusion that their dependence on the choice of the neighbour has to be taken into account.(78,79) All attempts to resolve this issue without the need of investigating a total of at least 400 amino acid pairs were based on the assumption that nearest neighbour effects depend solely on the neighbour, not on the targeted residue itself. This notion implies for instance that valine as neighbour has the same influence on alanine and aspartic acid. Along this way, Schwarzinger et al. investigated the influence of the X-residue on the adjacent glycine residues in Ac-GGXGG-NH2.(80) The use of glycine as representative target was later challenged by Kjaergaard and Poulsen, who introduced glutamine as a more useful alternative.(81)

The first experimental work aimed at exploring in more detail the residue specific nearest neighbour influence on chemical shifts and ^3^J(H ^N^ H ^Cα^) in short peptides was carried out by Jung et al.(82) These authors undertook the incredible task of measuring the ^3^J(H ^N^ H ^Cα^) coupling constants and H^N^ and H ^Cα^ chemical shifts of 361 Ac-XY-NH_2_ blocked dipeptides. They subjected their data solely to a statistical analysis in that they calculated the average coupling constant and the corresponding standard deviation for a given target amino **a**cid residue. Only coupling constants that were outside of this standard deviation were considered as reflecting nearest neighbour coupling. However, using the standard deviation as a criterion for statistical significance of difference is problematic in this context unless it reflects the statistical errors of the experimental values, which are generally small.(57) Nevertheless, it should be noted that neighbour induced changes of the J-coupling constant within a given target series is less pronounced for the dipeptide data set than it is for GXYGs. Significant differences between corresponding coupling constants are noteworthy for valine and leucine, mostly at the C-terminal position (A**L**, A**V, V**L, K**V**, with LL as an exception; cf. Figure S18). One might be tempted to rationalize such differences in terms of end effects, i.e. different influences of the terminal NH^3+^ and COOH groups of GXYG and the blocking groups the dimers used by Jung et al., but such an explanation seems to be at variance with the experimentally based notion that the conformational propensity of blocked dipeptides with a single aliphatic residues and two peptide groups resemble the one of the respective guest residue in GxG.(43) However, one has to note a peculiarity of the peptides used by Jung et al. Generally, dipeptides contain a single amino acid flanked by two peptide groups with methyl groups attached to the carbonyl carbon and the amide nitrogens. They can be regarded as suitable mimics of glycine. A corresponding system with two residues would actually be a blocked tripeptide with three peptide groups and terminal methyl groups. On the contrary, the C-terminal residue of the peptides used by Jung et al. are linked to a primary amide which differ from secondary amides. Their electronic structures can be expected to be different. In water, the amide I wavenumber (predominantly a C=O stretching mode) of a primary amide is significantly blue-shifted by approximately 20 cm^-1^ with respect to its position in secondary amide spectra (acetamide versus N-methylacetamide).(83) This suggests a stabilisation of the C=O double bond over C=N in the peptide’s resonance structure.

The study of Jung et al. was followed by the above-described work of Toal et al.(57) on GXYG peptides dealt within this paper. Compared with the former, it is rather limited in scope (15 peptide, later augmented by another 8 GXYG and 2 GXYZG peptides by Milorey et al.(52–54)). However, the data basis for each peptide is much broader (four ϕ- and one ψ-dependent scalar coupling constants and four different amide I’ profiles in different vibrational spectra). This large data set allowed for the construction of Ramachandran plots in terms of a superposition of two-dimensional Gaussian sub-distributions. While certainly still a simplification, the modelling of GxG peptides with the Gaussian model yielded a far better agreement with experimental data than molecular dynamics simulations with rather different force fields and water models.(44,84) Hence, it is justified to use the obtained GXYG Ramachandran distributions as a suitable approximation for a comparison with data sets obtained from restricted coil libraries. The use of the latter is based on the assumption that a large ensemble of residues located in ordered but not regularly structured segments of proteins produce conformational distributions representative of the coiled state of unfolded and disordered peptides and proteins. Restrictions to which even residues in structured loops of proteins are subjected are being thought to average out.

One of the main problems one encounters with the use of coil libraries is that even the enormous number of protein structures used for assembling the latest version of coil libraries insufficient to even explore the total of 8000 XYZ-segments which would account for exploring nearest neighbour interactions between all twenty amino acid residues. In order to address this issue, researchers average over all down- and upstream neighbours to obtain a sufficient number of entries to their Ramachandran plots. Bax and colleagues used this strategy to obtain a set of secondary chemical shifts and scalar coupling constants considered as a good approximation for the coiled state of peptides and proteins.(59,60) Ting et al. dealt with the limitations of their data set by an interpolation procedure called the hierarchical Dirichlet process.(63,85) Thus, they could produce Ramachandran plots with a 5° resolution in ϕ and ψ direction. Even with this approach they could not explore the full set of XYZ combinations, but they could restrict residue averaging to either the upstream or downstream neighbour of a selected target residue, which lead to Ramachandran plots for X**Y**-all and all-**X**Y.

The comparison of GXYG and coil library distributions revealed similarities, but also significant differences. In line with results on alanine containing peptides, the coil library confirms the comparatively high propensity of this residue type for pPII. Qualitatively, the coil library distribution also exhibits the relatively low pPII and the high turn (right-handed helical and type II/II i+2) propensities of aspartic acid. The relatively high β-strand propensity of valine is also reproduced. However, these similarities are more than outweighed by differences between the influence of nearest neighbour interactions. They are discussed in detail in the following.

In order to shed some light on the differences between tetrapeptide and coil library distributions, the average population of the combined mesostates of right-handed helical and the very similar type I/II β-conformation in GXYG and the Ting et al. library were compared to each other. Since the respective numbers are very low for the former the averaging was carried out over all neighbours in the investigated GXYG set. The result of this analysis is visualized by the upper plot in Figure 7. The regression analysis yielded a slope of 0.433±0.12 and a Pearson correlation coefficient value of 0.81. This value is indicative of a strong correlation, but one should keep in mind that the data basis of the plot is limited. The obtained correlation nevertheless indicates that the population of the right-handed helical/type I/II β region, which are merged in many Ramachandran plots, is by about a factor of two higher in the coil library, in spite of the relative to glycine negative influence of neighbours. Such a population enhancement is not surprising. One of the limitations of the use of short peptides as model systems for a statistical coil state of a protein is that the population of the right-handed helical region is underestimated compared with the situation in polypeptides and proteins. All theories of helix ⟺ coil transitions invoke a cooperative process where the helix formation goes through a nucleation process. The mole fraction of right-handed helices in GXG peptides or blocked dipeptides should be closely related to the nucleation parameters σ and v of the Zimm-Bragg and Lifson-Roig parameters, respectively.(70) With an increasing length of an oligo-peptide, the formation of short transient helices becomes more likely. The total fraction would depend on the respective values of the helix propensity parameters s and w. For oligo-alanines, the increase of the helical fraction with the number of residues has been predicted by MD simulations,(38,86) while experimental results are somewhat conflicting.(87,88) Generally, it is permissible to state that the increased population of right-handed helices and very similar turn-supporting structures in the coil library of Ting et al. is something to be expected. The above scaling factor of 2.3 should be taken as an average subject to the above indicated significant standard deviation of ±0.6.

If one ignores other less populated β- and γ-turn structures as well as asx-turns and left-handed helices the result of the above analysis suggests most of the differences between nearest neighbour interactions in GXYG and coil library segments involve redistributions between pPII and β-strand. If this supposition is correct, the total fraction of extended mesostates in the two systems should exhibit some correlation. The lower panel plot in Figure 8 compares the two fractions for all the pairs investigated experimentally. The Pearson coefficient of 0.64 still indicates a medium level correlation. However, it should be noted that the data basis for this correlation plot is much stronger than the one used for the comparison of right-handed helical populations. The slope of the correlation plot for the extended structures is 0.58±0.15, which suggests a reduced variability of the extended structure population in the coil library.

**Figure 8.**
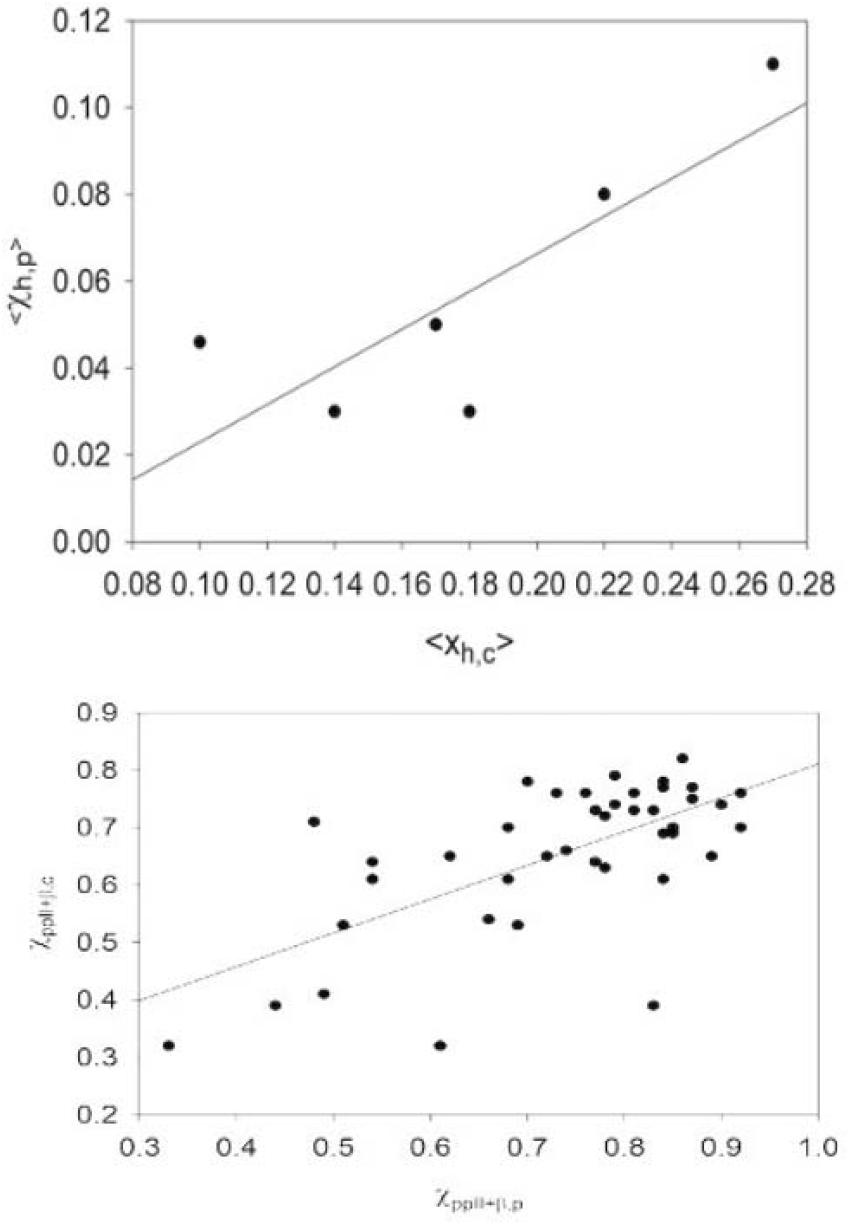
Correlation plots comparing mesostate populations in GxyG and the Ting et al. coil library.(63) Upper panel: Comparison of the combined population of right-handed α-helix and type I/II β-turn i+2 mesostates in GXYG (p) and the coil library (c) averaged over all target neighbours of A, V, S, D, L and F. Lower panel: Comparison of the combined population of pPII and all β-strand mesostates in GXYG (p) and the coil library (c). The solid lines result from a regression analysis described in the text.

The plots in Figure 6 reveal that nearest neighbour interactions in the coil library enhance the pPII population more than they do in the investigated GXYGs. The **A**V dimer in GXYG and all-AV exemplifies this difference, in that valine substantially reduces the pPII population of alanine in G**A**VG, while it enhances it in all-**A**V. In order to check whether an upstream valine would have the same influence on alanine we analysed the mesostate population of alanine in V**A**-all. The obtained populations of A are nearly identical with the ones of G**A**-all. This result suggests that at least for alanine the apparent influence of upstream neighbours is nearly eliminated by the averaging over all downstream neighbours. I checked this supposition by calculating the mesostate populations for all-X-all for all X-residues investigated in this study. For most of the later, the differences between all-X-all and GX-all were very small (< 5%). Only X=D showed a slight enhancement of pPII at the expense of type I/II β-turn.

In order to illustrate the possible influence of averaging over downstream neighbours I performed the following very simple calculation. It was assumed that the target residue in XY-all can adopt only pPII and β-strand conformations. The respective mole factions were calculated as follows:

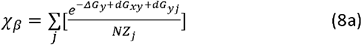

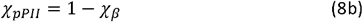

where the partition sum Z_j_ is written as:

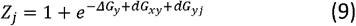

Δ*G*_*y*_ denotes the intrinsic Gibbs free energy difference between pPII and β-strand of residue Y, *dG*_*xy*_ is the free energy change caused by interaction with the upstream residue X and *dG*_*xj*_ represents the free energy change caused by interaction with the *j*^*th*^ downstream residue. N=20 is the number of residues. The distribution of interaction energies *dG*_*xj*_ was heuristically described by a Gaussian function:

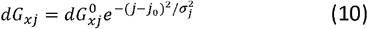

*j*_*0*_ is the position of the maximum the choice of which is arbitrary, σ_j_ is the standard deviation. All free energies are written in units of RT. Two scenarios, one with a high (0.67) and the other one with a low intrinsic pPII propensity (0.4) were considered. The correspond free energy differences are 0.7 and −0.4. Next, a dG_xy_ value of 0.3 was added to account for the general tendency of downstream neighbours. It reduced the corresponding pPII population to 0.59 and 0.33 (*vide infra*). In a third step, the Gaussian term in eq. (10) was added. A variation of the free energy term and of the standard deviation revealed that 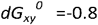, *j*_*0*_=5 and σ_j_=5 reproduced the assumed intrinsic propensities for both rather different scenarios. The results of this very heuristic calculation show that the averaging of Ramachandran plots of a distinct Y-residue over all natural downstream neighbours can practically neutralize the weaker nearest neighbour interactions with the upstream residue X. The reader should note that the rather large standard deviation of the utilized Gaussian distribution reflects an averaging over a relatively wide range of negative free energies.

The results of the above analysis suggest an asymmetry between nearest neighbour interactions in the coil library in that the interactions with upstream neighbours dominate over the ones with downstream neighbours. This notion is corroborated further by the observation that out of 26 residue pairs for which the coil library data indicate a nearest neighbour induced increase of the pPII population, 20 are of the all-**X**Y type, while 6 out of 7 neighbours that reduce the pPII population of the target residue are at the upstream position (X**Y**-all type).

The above unbalance between interactions with down- and upstream neighbours is still present in GXYG peptides, but to a lesser extent. In most peptides, nearest neighbour interactions reduce the pPII population. In 12 out of 19 cases this involves the respective upstream neighbours. pPII is enhanced in 14 cases, from which 9 involve downstream neighbours.

Another peculiarity of the coil library distributions is the rather large separation between the visible pPII and β-strand peaks in nearly all distributions. While the respective distances are often larger in GXYG than in respective GXG/GYG plots, they are not really close to the ones in the coil library. Figure S19 illustrates this point by depicting the ϕ dependence of the probability density for the serine residue in all-**S**L at ψ=150°. The plot actually reveals the existence of an additional very shallow maximum at 110° and of course the aforementioned dominance of pPII which is closer to the one in the corresponding GSLG plot (Figure S11). Similar tendencies can be seen in Ramachandran plots of the coil library used by the Sosnick group.(58) Interestingly, a very detailed coil library investigation of Jiang et al. revealed that the distance between the two basins depends on the χ_1_ angle of the side chain, where the g^-^ rotamer seems to produce the largest distance (their plots also represent an average over all neighbours in the library).(89) Hence, the discrepancy between coil library and peptide basin position noted in this study might actually reflect different populations of side chain rotamers. Needless to say, that this would further complicate the picture. Sosnick and coworkers actually provided computational evidence for coupling between side chain and backbone dynamics.(49)

In view of the differences between the neighbour dependence of residues in GXYG peptides and respective coil library dimers the question arises which of the two data sets are more useful for the identification of coil states. Being able to do so is necessary for the identification of residual structures in unfolded and disordered proteins.(59,60,90,91) The availability of a large data set for all natural amino acid residues in the coil library of Ting et al. is a clear advantage over the limited number of GXYG peptides that have thus far been investigated. The comparison of coil library and GXYG distributions has thus far revealed that averaging over neighbours might artificially diminish nearest neighbour effects and of course eliminates the side chain specificity of such interactions for a target residue. It is frequently believed that the differences caused by the latter are statistically insignificant. To check whether such a notion has merits I had a closer look at the reported ^3^J(H^N^H^Cα^) constants of two IDPs that are generally thought to be in an ideal coil state, namely the amyloid peptide Aβ_1-40_(92) and α-synuclein (92). Table 2 list the J-coupling constants of valine, phenylalanine, glutamic acid and alanine. The listed coupling constants of α-synuclein were taken from the C-terminal segment between residue 90 and 140, which does not exhibit any transient secondary structures. Note that experimental uncertainties of ^3^J(H^N^H^Cα^) are very small and do rarely exceed 0.1 Hz.(57,70,72) Since the values for α-synuclein had to be taken from a graphic representation, the corresponding error might be slightly larger. The data listed in Table 2 show that the coupling constant of valine varies within a range of 1.2 Hz. The corresponding intervals of alanine, glutamic acid and phenylalanine are 0.8, 1.4 and 0.7 Hz. Interestingly, while coupling constant values of valine scatter around the corresponding GVG value, the coupling constants for A and E are mostly lower, in line with an earlier observation for the tau protein.(56,91) This might reflect a higher degree of sampling of right-handed helical conformations. The Ting et al. library value for the corresponding all-F-all distributions is close to the average value of F (7.46 Hz). For all-A-all and all-V-all the Ting library values are significantly larger than the average (5.2 and 7.55 Hz, respectively).

**Table 2.**
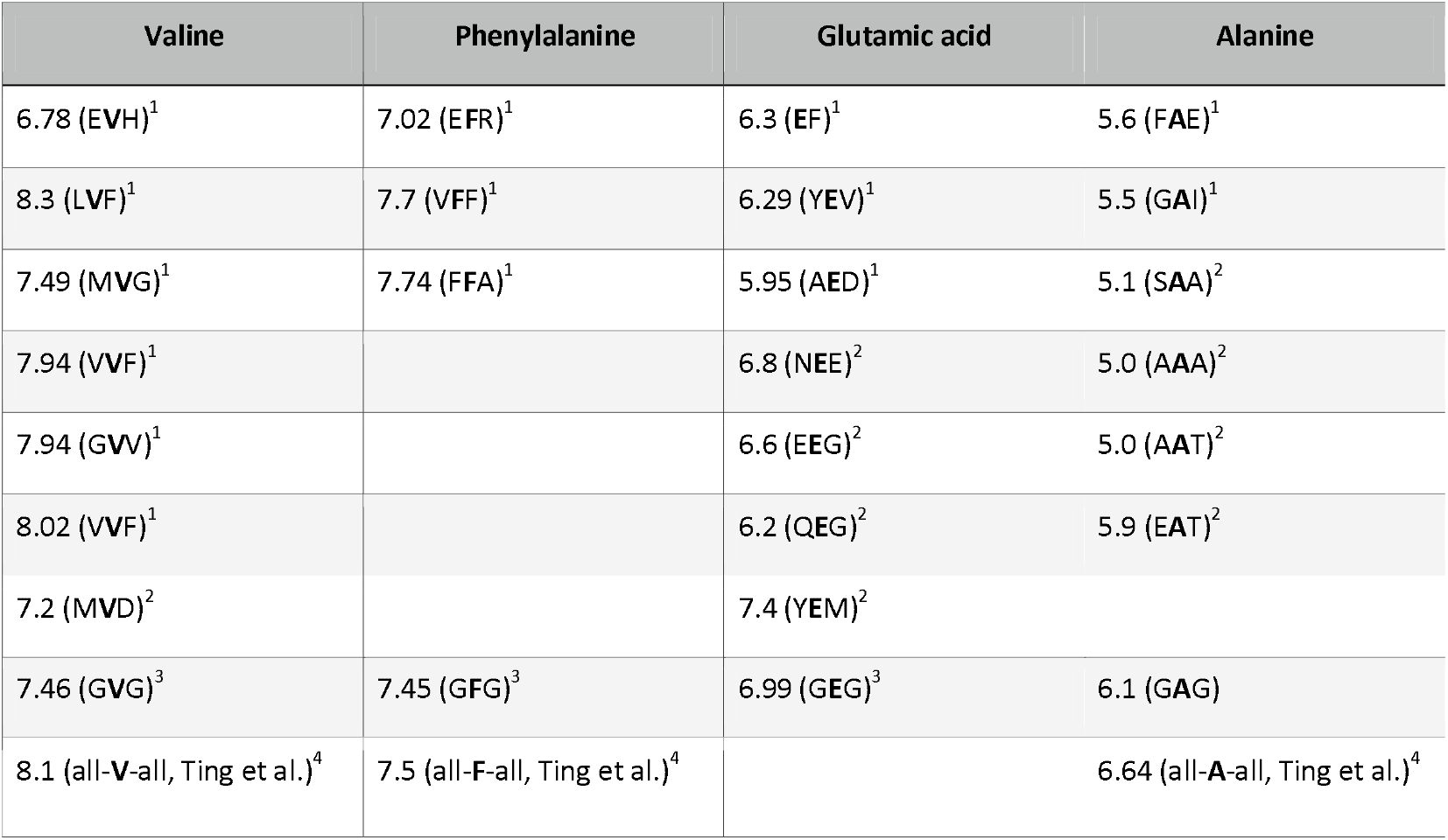
List of ^3^J(H^N^ H^Cα^) constants in units of Hz of the indicated amino acid residue in ^1^Aβ_1-40_,(93)^2^ α-synuclein (residues 89-140),(92)^3^ GXG peptides(70–72) and the coil libraries of ^4^Ting et al.(63) and ^5^Shen et al.(60) Values for the latter were obtained by averaging over both neighbours.

### Summary and Outlook

This paper presents a comparison of the influence of nearest neighbours on a selected number of amino acid residues in cationic tetra-peptides and the coil library of Ting et al. The way how the latter has been obtained allowed a direct comparison of amino acid pairs and the different influence of upstream and downstream neighbours. However, while these pairs are flanked by glycines in the tetra-peptides, averages were taken over all neighbours of the consider pairs in the coil library. As a consequence, either the influence of the upstream or of the downstream neighbour is explicitly taken into account. Ramachandran plots were compared by means of different metrics; i.e. mesostate populations, Hellinger distances, configurational entropy and several scalar J-coupling constants. Differences between GXYG and coil library distributions are noteworthy. Generally, nearest neighbour interaction induced changes are more pronounced in GXYG. For quite a noticeable number of pairs, the neighbours reduce the pPII propensity of amino acids in GXYG, while they increase it in the coil library. The averaging over either upstream or downstream neighbours for the construction of coil libraries reduce the specific influence of opposite neighbours. This result led to the conclusion that the residue specificities of nearest neighbour interactions should not be ignored. As a consequence, the identification of the coil state of a protein and/or an IDP of IDR is a complicated task. If one took short peptides as a starting point, one would have to investigate a total of 8000 GXYZG peptides just for a thorough assessment of nearest neighbour interactions. The exploration of non-nearest neighbour interactions would require an even larger number of hexa- and even hepta-peptides.

In principle, conducting molecular dynamics simulations could be a promising alternative venue. However, moving along this path requires combinations of force field and water models, which can reproduce intrinsic propensities as well as nearest neighbour interactions. Particularly regarding the latter, current approaches are still in their infancy.(44,45,51,54,94) The GXYG peptides used for this study should be used as benchmarks for the necessary force field development.

Why is it necessary to gain a comprehensive picture of nearest neighbour interactions? First, it would be necessary to obtain an accurate, residue specific picture of the statistical coil state of proteins, which can be used as a baseline for identifying the occurrence transient secondary structures. In the field of NMR spectroscopy, the Blackledge group developed the so-called flexible meccano program, which they used to extract a set of possible conformational ensembles for a given polypeptide/protein sequence by exploiting conformational distributions of amino acid residues in a restricted coil library.(90,95–97) Such ensembles are then used as a starting point for the structure analysis of IDPs by means of the ASTEROIDs algorithm (a selection tool for ensemble representations of intrinsically disordered states), which adjusts the conformational ensemble to constraints imposed by secondary chemical shifts, J-coupling constants (mostly ^3^J(H^N^H^C⍰^)), residual dipole coupling and experimentally obtained radii of gyration. From the description of flexible meccano in the literature (97) it is obvious that nearest neighbour interactions are solely considered by averaging. As argued in an earlier paper, this might lead to misidentification of transient secondary structures in IDPs such as the tau protein.(56) Second, correlations between conformational dynamics of residues reduce the configurational entropy of unfolded proteins to a significant extent.(46) Being able to quantify the latter is essential for an understanding of the thermodynamics of protein folding and disorder to order transitions. The effect of correlation dynamics on the entropy of GXYG is illustrated in Table S4. It compares the sum of individual configurational entropies of a selected set of X and Y with the total configurational entropy of XY calculated with an interaction model described in refs.(52,55) Most of the obtained values (GAVG and GALG being an exception) are negative, indicating entropy reduction. The majority of the respective room temperature Gibbs energy values are in 10^2^ J/mol range. Such values could become much larger for longer peptides and even proteins. GFDG stands out with an entropy reduction of 1.54 kJ/mol. Generally, the values are lower than the ones obtained from MD simulations of Baxa et al., who reported NNI reported entropy changes in the range of −1.5 and 2.1 kJ/residue for the unfolded state of ubiquitin.(49,50). However, their results and the ones presented in this study strongly suggest that the assumption of additivity of thermodynamic parameters (i.e. conformational entropy, free Gibbs energy of solvation, etc.) can now longer be considered as valid.

## Author contributions

RSS is the sole author of this article. He performed all the data analyses described therein and wrote the paper.

## Data availability

All data analysed in this article have been published and are therefore available. Reported secondary data are either listed in the Tables in the main manuscript and the Supporting Information or will be available from the author upon request.

## Acknowledgement

I would like to thank Dr. Roland L. Dunbrack from the Institute for Cancer Research, Fox Chase Cancer Center, Philadelphia, for providing me access to the files containing the data points of the Ramachandran plots reported in Ting et al. Dr. Siobhan E. Toal from Rowan University has checked the writing that concerned the content of ref.(57)

## Footnotes

† In Ting et al., H was multiplied by 100 to express it in units of %.

