## Supplemental tables S1-S2, supplemental figures S1-S19 for "Nearest Neighbour Interactions between Amino Acid Residues in Short Peptides and Coil Libraries"

*Reinhard Schweitzer-Stenner**

Department of Chemistry, Drexel University, 3141 Chestnut Street, Philadelphia, PA 19104, USA

**Contact:**

To be submitted to *RSC Advances*

Supporting Information

**Comparison with Ramachandran parameters reported by Toal et al.**

The J-coupling parameter values in Table S1 reveal that the fitting procedure did not yield as good a reproduction of the very high ^3^J(H^Cα^C’) values (> 3.0 Hz) measured for some GXYG peptides as the earlier procedure adopted by Toal et al.(1) However, I consider the current fitting parameters as more reliable for the following reason. The only way to obtain such large values for the coupling constants would require a substantial sampling of basins in the upper right quadrant of the Ramachandran plot with φ-values in the range between 60^o^ and 100^o^, where the Karplus curve for this J-coupling parameter significantly exceeds the value of 3 Hz.(2,3) In the upper left quadrant which is predominantly sampled by our peptides, only DFT-based Karplus curves for alanine residues exceed 3.0 Hz at the respective maximum position (φ=-120^o^).(4) Such a modeling would not allow the reproduction of the amide I’ profiles. A closer inspection of the six peptides for which such large ^3^J(H^Cα^C’) values had been obtained reveal that five of them contained serine. Interestingly, these values were obtained for serine itself as well as for its respective neighbors A, V and K. One may therefore wonder whether the specific Karplus curves for these residues would have a more pronounced maximum in the upper left quadrant region (we have to remember here that the parameters of empirical Karplus curves result from an average over many different residue types). The above DFT calculation for alanine suggests that residue specificity might lead to differences from empirical Karplus curves in this region. It is very likely that the seemingly better theoretical ^3^J(H^Cα^C’) values reported in our earlier study reflect an overestimation caused by the low-resolution scanning of the Ramachandran plot (with increments varying between 6^o^ and 8^o^) and the fixed Ramachandran sub-space used for all GXYG peptides investigated (cf. the Theory section of the main manuscript). As stated in the later, the current fitting of J-coupling constants used the standard resolution of 2^o^ of our Gaussian model.

Generally, the differences between the mole fractions displayed in Figures S3-S6 and corresponding values reported by Toal et al.(1) are small or moderate (Table S2). This notion applies in particular to the alanine series, where the hierarchy of pPII reduction by its neighbors was reproduced. The reproduction of the very pronounced influence of valine in G**A**VG should be emphasized in this context. As in Toal et al., the influence of unlike nearest neighbors on the conformational propensities of serine and lysine is small. For the leucine series, only the fractions of GL**L**G deviates significantly from the ones in Toal et al. in that the β-strand propensity is significantly enhanced at the expense of pPII. The aspartic acid series has changed moderately in that pPII propensities are now slightly lower in G**D**VG and G**DL**G. The valine series are again very similar with the exception of GD**V**G, where the β-strand propensity is now more enhanced at the expense of pPII. Taken together, it is safe to state that the core message of Toal et al., namely the significant influence of neighbors on the mole fraction of alanine and aspartic acid, remains valid. Interestingly, as also shown in Figures S3-S7, NNIs become more pronounced in GDDG and also for phenylalanine containing peptides. The latter results emerged from the more recent work of Milorey and colleagues.(5–7) In addition to the changes of mole fractions, the new analysis also yielded some changes in the position of Gaussian sub-distribution, which can be inferred from Table S2. As already indicated by the data of Toal et al. NNI change such positions even in cases where the respective change of mole fractions is rather small (e.g. for most peptides of the leucine series).

**Table S1.** List of (Gaussian) parameters used to construct a (local, restricted) random coil like distribution. The first four columns list the locations and widths of the Gaussian sub-distributions in the Ramachandran space. The first column lists the mole fractions of the respective sub-distributions. Taken from ref. (8). Copyright by the Royal Society of Chemistry 2014

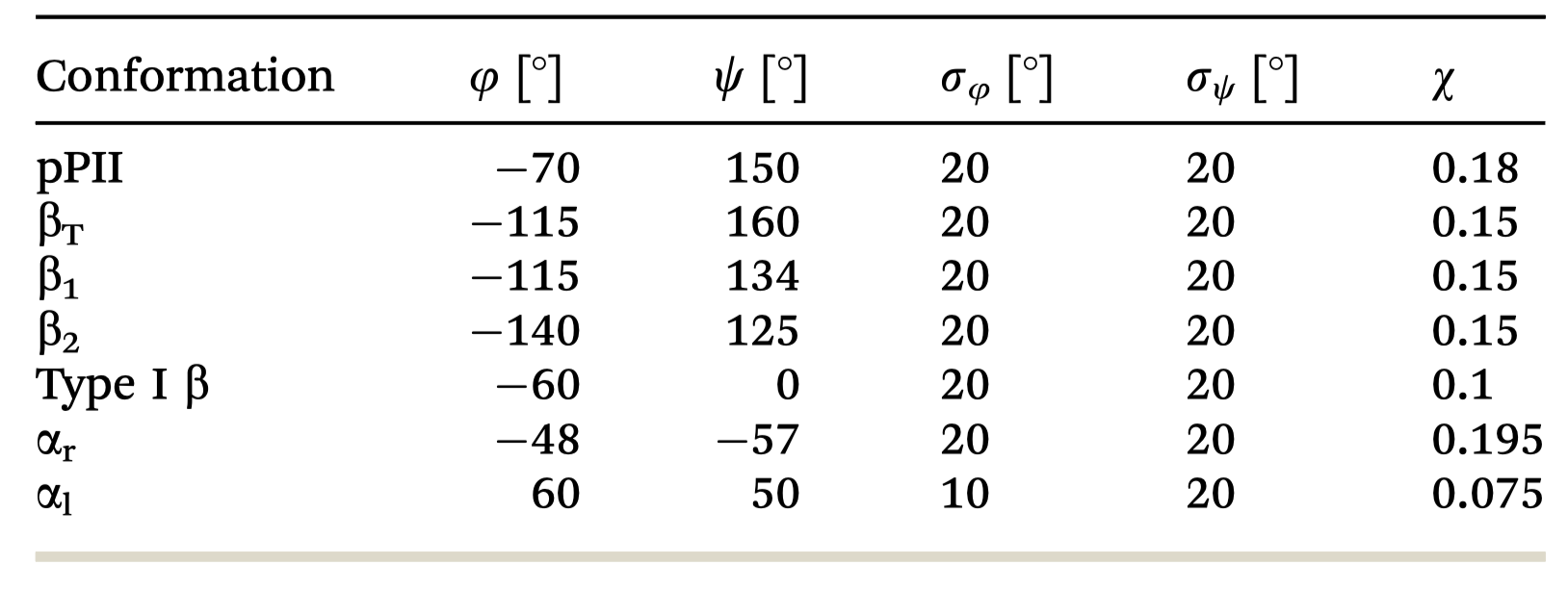

**Table S2.** List of new and old calculated J-coupling constant of GXYG peptides compared with the experimental values reported by Toal et al.(1) The respective X and Y residues for which these coupling constants are reported are typed in bold. If not indicated otherwise the reported coupling constants are of the ^3^J type. All values are listed in units of Hz. The last column lists the reduced χ^2^ value of the new fits to the experimental coupling constants. The respective uncertainties of the ^3^J-constants were calculated as a superposition of experimental uncertainties and errors produced by the uncertainties of the used Karplus parameters as reported by Wand and Bax.(2) Since the values for the uncertainties of the Karplus parameters used for the ^1^J-constant are not known, the considered error reflects solely the experimental uncertainty. This can lead to a significant overestimation of this parameter’s contribution to the reduced chi-square value. Generally, the thus calculated errors are smaller than the root mean square values taken from the data scattering in J-coupling plots.(3)

|  | H^N^H_α_  new | H^N^H_α_  old | H^N^H_α_  exp | H^N^C’  new | H^N^C’  old | H^N^C’  exp | H_α_C’  new | H_α_C’  old | H_α_C’  Exp | H^N^C_β_  new | H^N^C_β_  old | H^N^C_β_  exp | ^1^J(NC_α_)  new | ^1^J(NC_α_)  old | ^1^J(NC_α_)  exp | χ^2^_r_ |
| --- | --- | --- | --- | --- | --- | --- | --- | --- | --- | --- | --- | --- | --- | --- | --- | --- |
| DA | 7.46 | 7.18 | 7.22 | 1.09 | 1.08 | 0.97 | 2.76 | 3.0 | 3.15 | 1.32 | 1.5 | 1.54 | 11.84 | 11.75 | 11.81 | 1.4 |
| DA | 6.48 | 6.62 | 6.63 | 1.14 | 1.24 | 1.2 | 2.33 | 2.50 | 2.19 | 1.70 | 1.61 | 1.98 | 11.58 | 11.19 | 11.37 | 1.3 |
| DK | 6.41 | 6.31 | 6.19 | 0.87 | 0.84 | 0.5 | 2.50 | 2.79 | 2.7 | 1.94 | 1.76 | 1.47 | 11.79 | 11.73 | 11.93 | 6.9 |
| DK | 6.66 | 6.77 | 6.63 | 0.96 | 1.1 | 1.03 | 2.0 | 2.2 | 2.09 | 1.74 | 1.60 | 1.54 | 11.36 | 10.9 | 11.27 | 1.0 |
| DL | 7.28 | 6.90 | 6.97 | 1.05 | 0.92 | 0.9 | 2.25 | 2.65 | 2.75 | 1.40 | 1.52 | 1.57 | 11.57 | 11.3 | 11.34 | 6.8 |
| DL | 6.84 | 6.81 | 6.78 | 0.95 | 0.90 | 0.84 | 2.12 | 2.45 | 2.45 | 1.67 | 1.60 | 1.66 | 11.52 | 10.91 | 10.58 | 2.8 |
| DV | 7.32 | 6.59 | 7.08 | 0.88 | 0.91 | 0.87 | 2.77 | 3.0 | 3.0 | 1.53 | 1.70 | 1.54 | 11.72 | 11.75 | 11.90 | 0.9 |
| DV | 8.03 | 8.23 | 8.25 | 0.78 | 0.81 | 0.81 | 2.63 | 2.80 | 2.80 | 1.37 | 1.30 | 1.28 | 11.30 | 11.19 | 11.23 | 1.2 |
| SA | 7.1 | 7.4 | 6.86 | 0.99 | 1.0 | 0.97 | 2.74 | 3.2 | 3.26 | 1.55 | 1.8 | 1.79 | 11.70 | 12.2 | 12.2 | 5.7 |
| SA | 6.66 | 6.20 | 6.35 | 0.96 | 0.8 | 0.79 | 2.7 | 3.2 | 3.2 | 1.77 | 2.0 | 2.05 | 11.11 | 10.9 | 11.3 | 7.3 |
| SK | 6.90 | 6.60 | 6.62 | 0.96 | 0.9 | 0.88 | 2.94 | 3.3 | 3.52 | 1.67 | 1.8 | 1.86 | 11.69 | 12.2 | 12.53 | 2.32 |
| SK | 7.22 | 7.27 | 7.22 | 1.03 |  |  | 2.96 | 3.2 | 3.52 | 1.48 | 1.6 | 1.54 | 11.36 | 11.0 | 11.3 | 6.0 |
| SL | 6.89 | 7.3 | 6.86 | 0.89 | 1.0 | 0.44 | 2.54 | 2.4 | 2.48 | 1.60 | 1.5 | 1.79 | 11.71 | 12.1 | 12.2 | 1.7 |
| SL | 7.22 | 7.3 | 7.34 | 0.94 | 0.8 | 0.66 | 2.25 | 2.4 | 2.59 | 1.51 | 1.60 | 1.60 | 11.69 | 10.8 | 11.15 | 3.3 |
| SV | 7.02 | 6.9 | 6.95 | 0.87 | 0.9 | 0.87 | 2.86 | 3.3 | 3.52 | 1.67 | 1.90 | 1.92 | 11.80 | 12.2 | 12.2 | 7.5 |
| SV | 8.0 | 8.2 | 8.43 | 0.77 | 0.8 | 0.76 | 3.17 | 3.1 | 3.52 | 1.33 | 1.5 | 1.41 | 11.20 | 11.3 | 11.30 | 4.3 |
| AL | 6.58 | 6.75 | 6.80 | 0.98 | 1.3 | 1.36 | 2.05 | 2.0 | 2.08 | 1.77 | 2.30 | 2.29 | 11.09 | 11.30 | 11.32 | 5.1 |
| AL | 7.47 | 7.36 | 7.4 | 0.75 | 0.85 | 0.53 | 2.28 | 2.32 | 2.29 | 1.53 | 1.75 | 1.66 | 11.37 | 11.34 | 11.40 | 2.5 |
| KL | 6.76 | 6.40 | 6.44 | 1.06 | 1.01 | 0.96 | 2.08 | 2.22 | 2.20 | 1.63 | 1.69 | 1.73 | 11.32 | 11.32 | 11.44 | 1.6 |
| KL | 7.27 | 7.0 | 7.10 | 0.83 | 0.75 | 0.51 | 2.27 | 2.39 | 2.37 | 1.57 | 1.72 | 1.73 | 11.29 | 11.54 | 11.26 | 5.4 |
| LL | 7.78 | 7.60 | 7.62 | 0.73 | 0.88 | 0.77 | 2.89 | 3.20 | 3.52 | 1.43 | 1.52 | 1.54 | 10.58 | 10.90 | 10.65 | 7.9 |
| LL | 8.17 | 8.85 | 9.08 | 0.79 | 0.9 | 0.84 | 2.71 | 3.44 | 3.52 | 1.21 | 1.80 | 1.73 | 11.18 | 11.20 | 11.16 | 15.2 |
| VL | 7.77 | 8.46 | 8.46 | 0.76 | 0.75 | 0.77 | 2.54 | 2.50 | 2.53 | 1.40 | 1.60 | 1.54 | 11.07 | 11.1 | 11.07 | 2.4 |
| VL | 7.92 | 8.10 | 8.15 | 0.63 | 0.63 | 0.58 | 2.24 | 2.40 | 2.37 | 1.42 | 1.49 | 1.54 | 11.38 | 11.45 | 11.24 | 0.8 |
| AV | 6.42 | 6.60 | 6.60 | 1.17 | 1.10 | 1.27 | 2.41 | 2.50 | 2.56 | 1.71 | 2.21 | 2.40 | 11.20 | 11.43 | 11.45 | 5.5 |
| AV | 7.21 | 7.19 | 7.15 | 0.92 | 1.08 | 1.16 | 2.09 | 2.29 | 2.35 | 1.52 | 1.56 | 1.5 | 11.44 | 11.29 | 11.33 | 2.7 |
| KV | 7.37 | 7.24 | 7.21 | 0.95 | 0.90 | 0.87 | 2.78 | 2.70 | 2.72 | 1.46 | 1.91 | 1.84 | 11.23 | 11.33 | 11.23 | 4.3 |
| KV | 7.33 | 7.40 | 7.42 | 0.88 | 1.0 | 0.99 | 2.39 | 2.10 | 2.06 | 1.51 | 1.54 | 1.52 | 11.48 | 11.55 | 11.46 | 2.5 |

**Table S3** List of statistical weights and positions of Gaussian sub-distributions obtained from analyses of J-coupling constants and amide I’ profiles of the indicated tetra-peptides (the glycine residues have been omitted in the first row). Earlier results reported by Toal et al.(1) are given in parenthesis. Conformation 1: pPII, conformation 2: β-strand, conformation 3: right-handed helical. Conformations 4 and 5 are residue dependent, they predominantly encompass left-handed helical, asx- and γ-turns. A solid red frame indicates that corresponding values obtained from the new and old analysis are very similar. Red dashed frames indicate moderate agreement.

|  | χ_1_ | (φ_1_,ψ_1_) | χ_2_ | (φ_2_,ψ_2_) | χ_3_ | (φ_3_,ψ_3_) | χ_4_ | (φ_4_,ψ_4_) | χ_5_ | (φ_5_,ψ_5_) |
| --- | --- | --- | --- | --- | --- | --- | --- | --- | --- | --- |
| DA | 0.39  (0.24) | -82,175  (-78,175) | 0.38  (0.48) | -132,175  (-132,175) | 0.09  (0.09) | -50,0  (-50,0) | 0.14  (0.19) | 65,150  (65,150) |  |  |
| DA | 0.62  (0.62) | -72,165  (-69,155) | 0.23  (0.23) | -115,165  (-115,155) | 0.05  (0.05) | -30,-30 | 0.1  (0.1) | 50,-50  (50,-50) |  |  |
| DK | 0.4  (0.4) | -70,175  (-75,175) | 0.34  (0.45) | -115,175  (-130,175) | 0.08  (0.08) | -40,0  (-50,0) | 0  (0.08) | 60,0  (-50,0) | 0.18  (0.07) | 100,150  (65,130) |
| DK | 0.5  (0.46) | -66,150  (-66,148) | 0.38  (0.42) | -115,150  (-115,135) | 0.05  (0.05) | -65,-35  (-65,-35) | 0.03  (0.03) | 70,45  (70.45) | 0.04  (0.04) | -60,110  (-60,110) |
| DL | 0.45  (0.4) | -75,158  (-75,145) | 0.43  (0.43) | -130,158  (-130,145) | 0.07  (0.07) | -50,0  (-50,0) | 0.05  (0.1) | 80,60  (65,130) |  |  |
| DL | 0.49  (0.49) | -67,155  (-75,145) | 0.42  (0.43) | -110,155  (-130,145) | 0.05  (0.05) | -50,-40  (-50,-40) | 0.04  (0.04) | 60,40  (-65,112) |  |  |
| DV | 0.4  (0.45) | -75,170  (-78,150) | 0.4  (0.4) | -115,170  (-130,145) | 0.07  (0.07) | -50,0  (-50,0) | 0.15  (0.10) | 65,130  (65,130) |  |  |
| DV | 0.15  (10.30) | -74,165  (-74,145) | 0.65 (0.50) | -120,145  (-120,140) | 0.1  (0.1) | -60,-30  (-60,-30) | 0.1  (0.1) | -95,130  (-95,130) |  |  |
| SA | 0.37  (0.37) | -70,172  (-75,152) | 0.4  (0.40) | -115,170  (-118,150) | 0.09  (0.1) | -50,0  (-50,0) | 0.08  (0.06) | 60,0  (70,0) | 0.07  (0.07) | 70,160  (70,160) |
| SA | 0.4  (0.5) | -69,155  (-69,150) | 0.31  (0.31) | -115,155  (-115,140) | 0.07  (0.07) | -60,-30  (-60,-30) | 0.17  (0.07) | 70,60  (60,50) | 0.05  (0.05) | -50,50  (-50,50) |
| SK | 0.36  (0.31) | -70,170  (-75,165) | 0.39  (0.44) | -103,170  (-103,165) | 0.05  (0.1) | -50,-30  (-50,-15) | 0.05  (0.05) | 65,30  (70,0) | 0.15  (0.04) | 65,145  (75,145) |
| SK | 0.37  (0.42) | -66,165  (-66,145) | 0.43  (0.49) | -115,165  (-125,135) | 0.03  (0.03) | -70,-70  (-65,35) | 0.17  (0.06) | 65,45  (70,45) |  |  |
| SL | 0.37  (0.37) | -72,165  (-74,165) | 0.42  (0.42) | -112,165  (-132,155) | 0.1  (0.1) | -50,-15  (-50,-15) | 0.05  (0.05) | 70,0  (70,0) | 0.06  (0.06) | 65,145  (65,160) |
| SL | 0.42  (0.48) | -70,165  (-70,152) | 0.46  (0.40) | -120,165  (-120,142) | 0.07  (0.03) | -50,-50  (-50,-50) | 0.  (0.04) | (-70,0) | 0.05  (0.05) | 70,20  (70,45) |
| SV | 0.37  (0.47) | -68,165  (-70,155) | 0.39  (0.41) | -110,165  (-110,155) | 0.05  (0.05) | -50,-15  (-50,-15) | 0.20 (0.07) | 70,160  (65,160) |  |  |
| SV | 0.2  (0.28) | -80,152  (-74,152) | 0.54 (0.51) | -120,147  -120,140 | 0.05  (0.05) | -60,-30  (-60,-30) | 0.11 (0.06) | 60,0  (60,0) | 0.1  (0.1) | 95,130  (95,130) |
| AL | 0.55  (0.65) | -69,145  (-69,145) | 0.29  (0.29) | -120,145  (-115,140) | 0.06  (0.06) | -60,30  (-55,30) | 0.05  (0) | -70,-40 | 0.05  (0) | 60,40 |
| AL | 0.42  (0.42) | -78,150  (-70,150) | 0.46  (0.46) | -120,150  (-120,140) | 0.03  (0.03) | -70,10  (-70,10) | 0.03  (0.03) | -50,-50  (-50,50) | 0.06  (0.02) | 100,115  (100,115) |
| KL | 0.52  (0.52) | -66,150  (-66,145) | 0.38  (0.38) | -120,145  (-120,140) | 0.04  (0.04) | 70,30  (70,30) | 0.06  (0.06) | -65,35  (-65,35) |  |  |
| KL | 0.43  (0.43) | -70,142  (-70,142) | 0.47 (0.52) | -115,150  (-120,150) | 0.04  (0.05) | -60,-50  (-50,50) | 0.06  (0.0) | 70,60 |  |  |
| LL | 0.37  (0.42) | -80,130  (-76,145) | 0.42  (0.42) | 120,130  (-98,160) | 0.04  (0.04) | -60,-50  (-50,-50) | 0.02  (0.07)0 | -70,60  (-65,112) | 0.15  (0.03) | 70,50  (100,115) |
| LL | 0.25  (0.4) | -80,148  (-76,145) | 0.59  (0.49) | -120,148  (-98,160) | 0.02  (0.02) | -65,112  (-65,112) | 0.05 (0.05) | -50,-50  (-50,-50) | 0.09 (0.04) | 70,60  (100,115) |
| VL | 0.33  (0.33) | -80,145  (-74,152) | 0.48  (0.48) | -120,145  (-120,145) | 0.08 (0.11) | 75,0  (60,0) | 0.11 (0.08) | -60,60  (-60,60) |  |  |
| VL | 0.4  (0.4) | -85,150  (-70,145) | 0.48  (0.48) | -120,150  (-123,142) | 0.05  (0.05) | -60,-50  (-50,-50) | 0.05  (0.05) | -67,112  (-67,112) | 0.02  (0.02) | 100,105  (100,115) |
| AV | 0.4  (0.4) | -60,153  (-75,155) | 0.4  (0.48) | -120,153  (-120,155) | 0.05  (0.05) | -60,-30  (-60,-30) | 0.13  (0.05) | 80,-40  (50,-50) | 0.02  (0.02) | -60,100  (-60,100) |
| AV | 0.39  (0.39) | -72,155  (-72,155) | 0.45  (0.45) | -120,155  (-120,155) | 0.09  (0.09) | -60,-30  (-60,-30) | 0.05  (0.05) | -60,60  (-60,60) | 0.02  (0.02) | 95,115  (95,115) |
| KV | 0.3  (0.4) | -66,150  -66,145 | 0.48  (0.48) | -120,145  (-120,145) | 0.15  (0.05) | 75,30  (75,45) | 0.07  (0.07) | -65,35  (-65,35) |  |  |
| KV | 0.44  (0.44) | -75,155  (-75,155) | 0.43  (0.4) | -115,155  (-110.100) | 0.07  (0.07) | 65,35  (65,35) | 0.02  (0.02) | -60,-30  (-60,-30) | 0.04  (0.04) | -60,60  (-60,60) |

**Table S4.** List of the Gibbs energy contributions of the configurational entropy calculated as the sum over the two residues of GXYG peptides and as total entropy of the configuration produced by nearest neighbor interactions, which were calculated by an interaction model described by Schweitzer-Stenner et al.(5)

| Peptide | TΔS_conf_-TΔS_sum_[kJ/mol] | TΔS_conf_[kJ/mol] | TΔS_csum_[kJ/mol] |
| --- | --- | --- | --- |
| GAFG | -0.09 | 4.024 | 4.11 |
| GFAG | -0.525 | 3.59 | 4.12 |
| GAVG | 0.64 | 5.34 | 4.70 |
| GDAG | -0.68 | 4.16 | 4.84 |
| GALG | 0.22 | 4.24 | 4.03 |
| GDDG | -0.24 | 6.011 | 6.25 |
| GDVG | -0.85 | 5.17 | 6.03 |
| GDFG | -0.73 | 4.72 | 5.45 |
| GFDG | -1.54 | 3.91 | 5.45 |
| GRRG | -0.73 | 4.70 | 5.43 |

**Figure S1**. Visualization of the reduced number of data points in the Ramachandran plot used to reproduce the amide I’ profile of GxyG peptides. The data points overlay a Ramachandran plot alanine in GSLG.
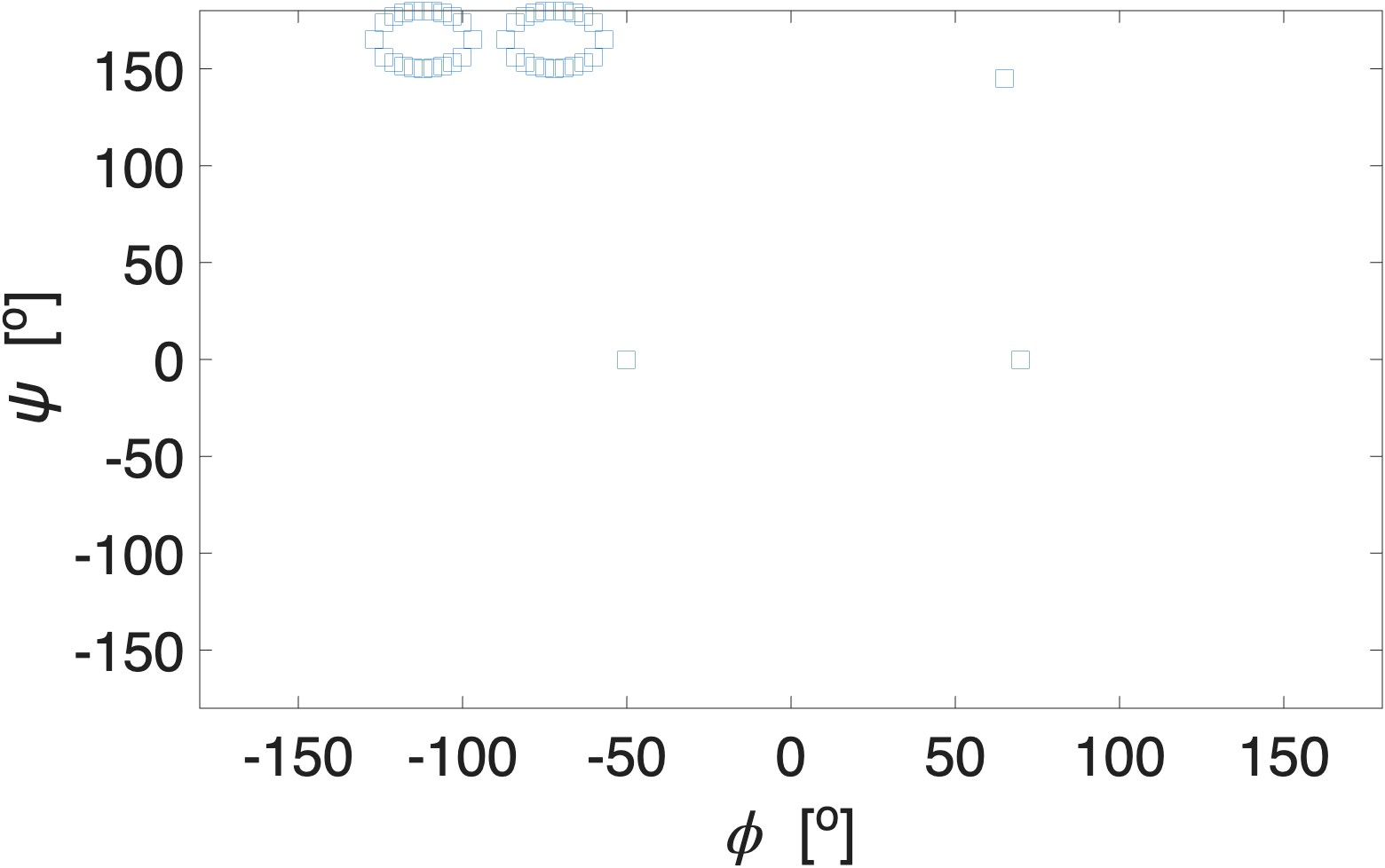

**Figure S2** Representation of mesostates in the Ramachandran plot. The labels aβ, pβ, βt, pPII, and α correspond to antiparallel β, parallel β, transitional β, polyproline II, and right-handed helix, respectively. The turn-supporting mesostates are marked blue (open access).

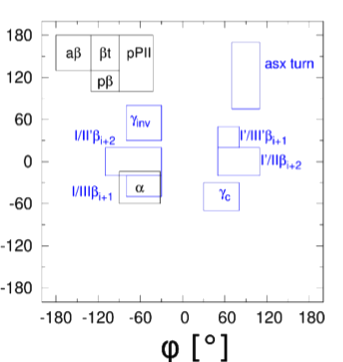

**Figure S3.** Profiles of the amide I’ band in the isotropic Raman, anisotropic Raman, FTIR and VCD spectrum of the indicated fully protonated tetrapeptides in D_2_O. The red data points represent experimental data. The blue line results from a simulation based on the fits to the respective J-coupling constants in Table S1. The broad band at 1625 cm^-1^ is assignable to the C=O stretching mode of the C-terminal carboxylate group. In order to account for this band in the simulated spectrum, a unnormalized Gaussian band has been add
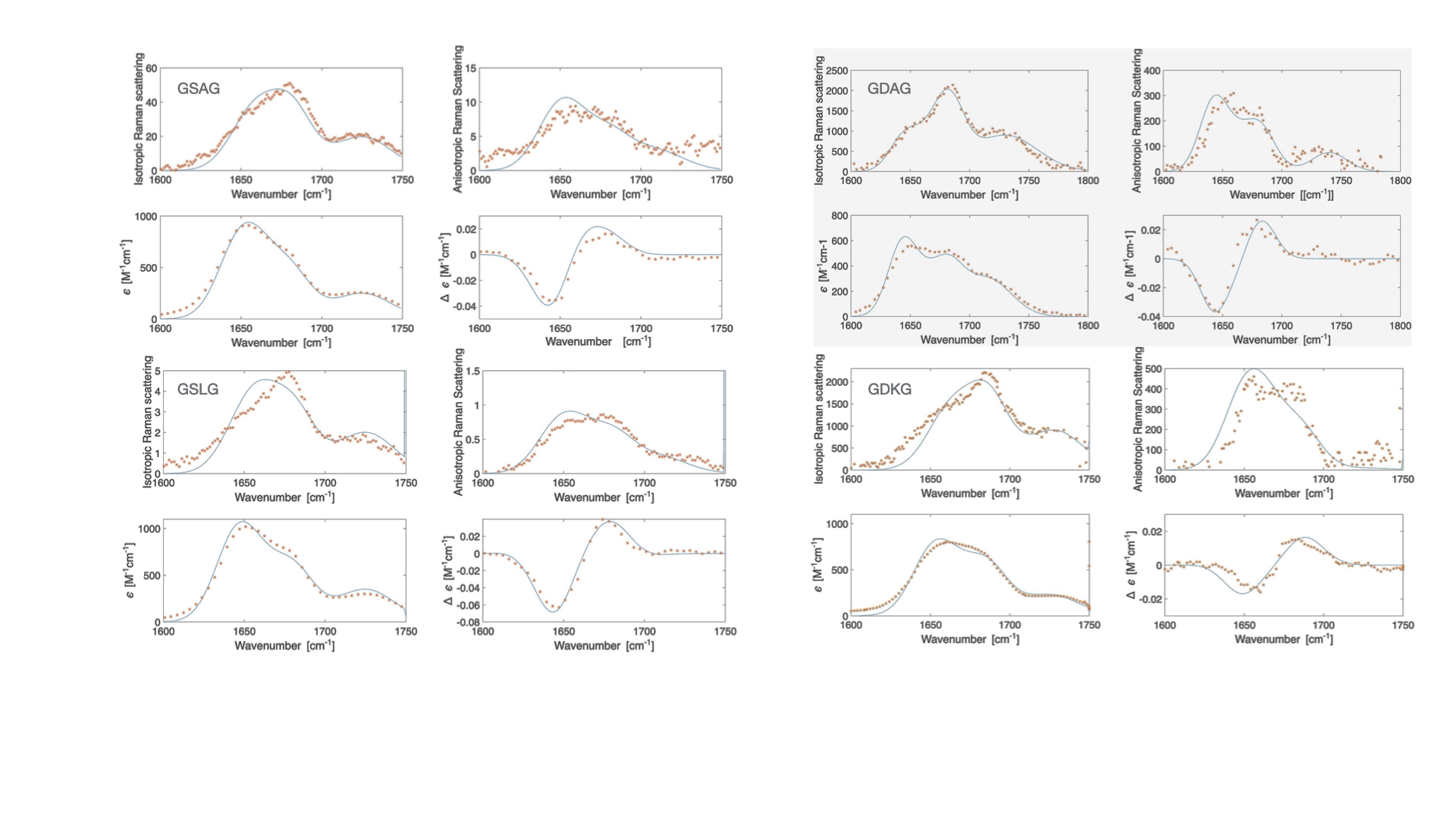
ed at the respective position. The different dispersions of the amide I’ reflect the specific spectroscopic changes due to the excitonic coupling between the amide I’ modes of the three peptide groups. The coupling strength and the redistributions of intensities as well as the rotational strength displayed in the VCD spectrum depends on the backbone dihedral angles φ and ψ of the non-terminal residues D and A. Details of the theory can be found in refs.(9,10) The experimental data have been taken from Toal et al.(1)

**Figure S4.** Mole fractions (statistical weights) of pPII, β-strand and of the combined turn-supporting conformations (including right- and left-handed helical) for a series of alanine (upper panel) and leucine (lower panel) containing peptides. The displayed mole fraction values were obtained from a re-analysis of GXYG data reported by Toal et al.(1) Values for GAFG and GFAG were taken from Schweitzer-Stenner et al.(5)

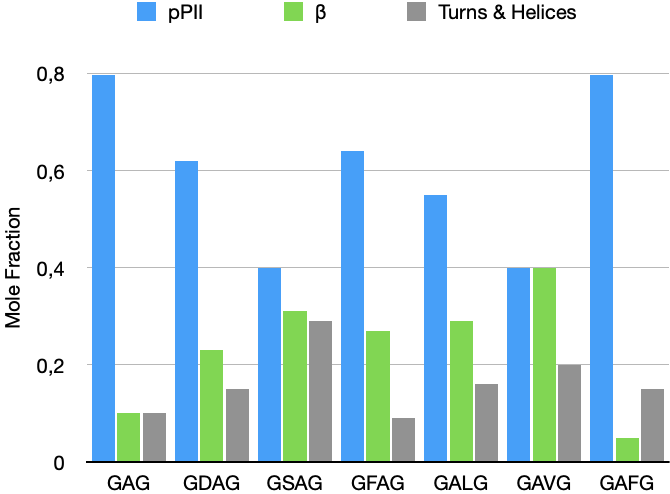

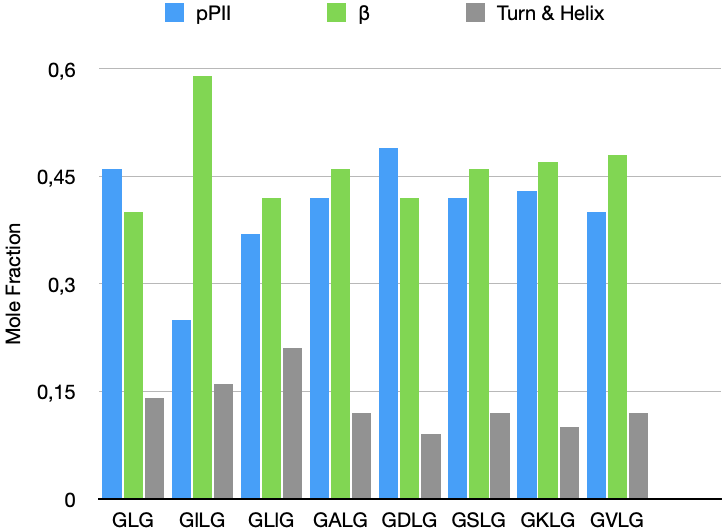

**Figure S5.** Mole fractions (statistical weights) of pPII, β-strand and of the combined turn-supporting conformations (including right- and left-handed helical) for a series of aspartic acid (upper panel) and serine (lower panel) containing peptides. The displayed mole fraction values were obtained from a re-analysis of GxyG data reported by Toal et al.(1) Values for GDFG and GFDG were taken from Milorey et al.(6) Note that the mole fractions for GDG reflect the result of a later re-analysis of the data reported in ref.(6) which revealed a higher pPII propensity than the one reported earlier.(11)

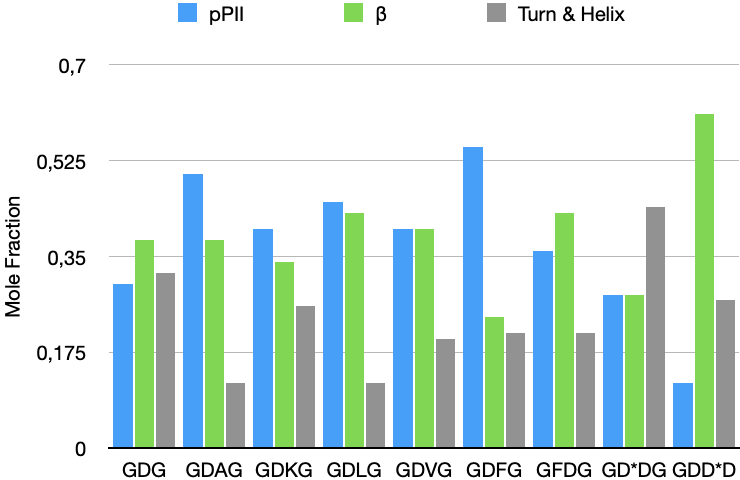

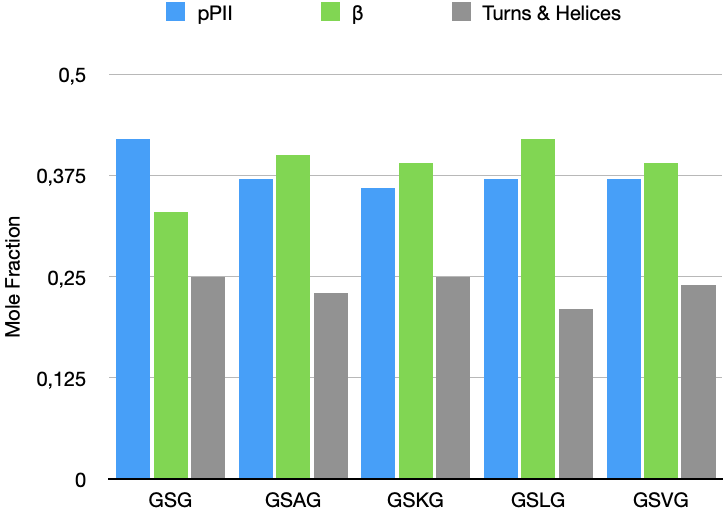

**Figure S6**. Mole fractions (statistical weights) of pPII, β-strand and of the combined turn-supporting conformations (including right- and left-handed helical) for a series of valine (upper panel) and lysine (lower panel) containing peptides. The displayed mole fraction values of valine containing peptides were obtained from a re-analysis of GXYG data reported by Toal et al.(1) The values for phenylalanine containing peptides and for G**K**KG and GK**K**G were taken from Schweitzer-Stenner et al.(5)

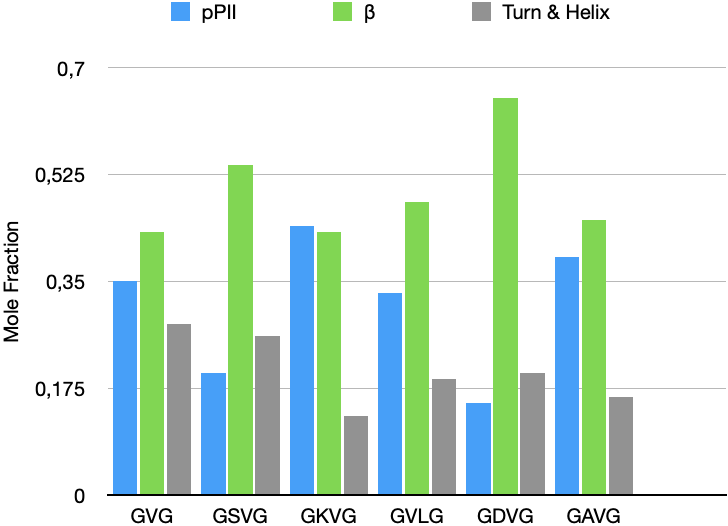

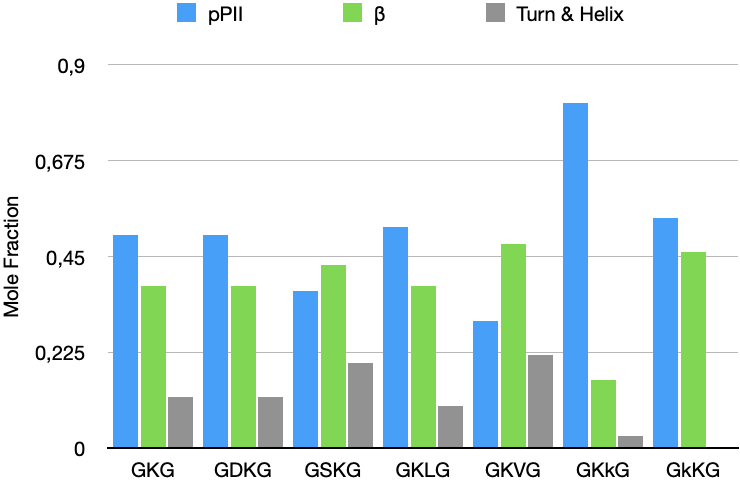

**Figure S7**. Mole fractions (statistical weights) of pPII, β-strand and of the combined turn-supporting conformations (including right- and left-handed helical) for a series of phenylalanine containing peptides which were taken from Schweitzer-Stenner et al.(5)

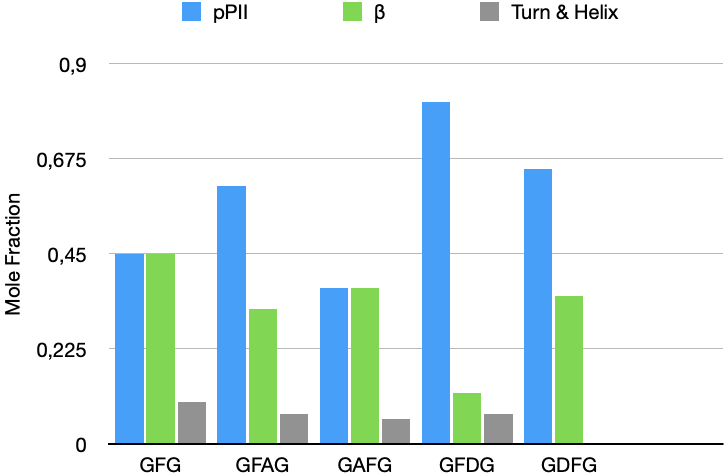

**Figure S8.** Gibbs energy diﬀerence between pPII and β-strand as a function of the pPII fraction calculated for diﬀerent fractions occupying the region above ψ = 100^o^ in the right-hand half of the Ramachandran plot**.** Taken from ref.(8) Copyright by the Royal Society of Chemistry 2023

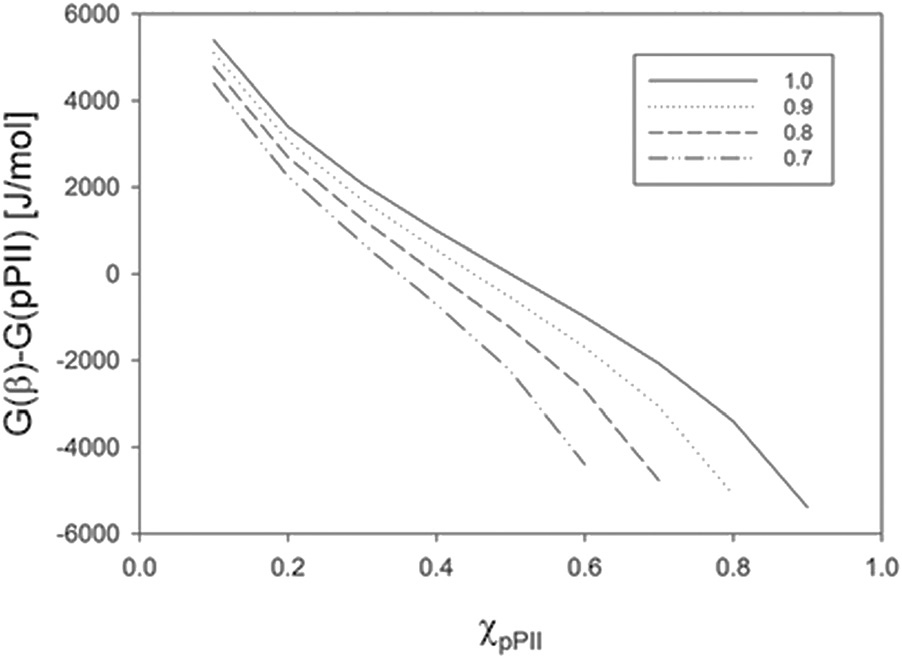

**Figure S9.** Ramachandran plots of alanine residues in GAG, all-**A**G, GD**A**G, D**A**-all, GS**A**G, S**A**-all, G**A**FG, all-F**A**, GF**A**G and F**A**-all. Corresponding plot for valine and leucine as neighbors of alanine are shown in Figure 1 of the main manuscript. All explored peptides were in their cationic state. The plots for the peptides were constructed with a Gaussian model and the parameters listed in ref.(12) for GAG, in Table S2 for GD**A**G, GS**A**G and for G**A**LG, and in ref.(5) for GF**A**G and G**A**FG. The plots for the coil library segments were built with files obtained from the website of the Dunbrack group with the kind permission of Dr. Dunbrack.(13)

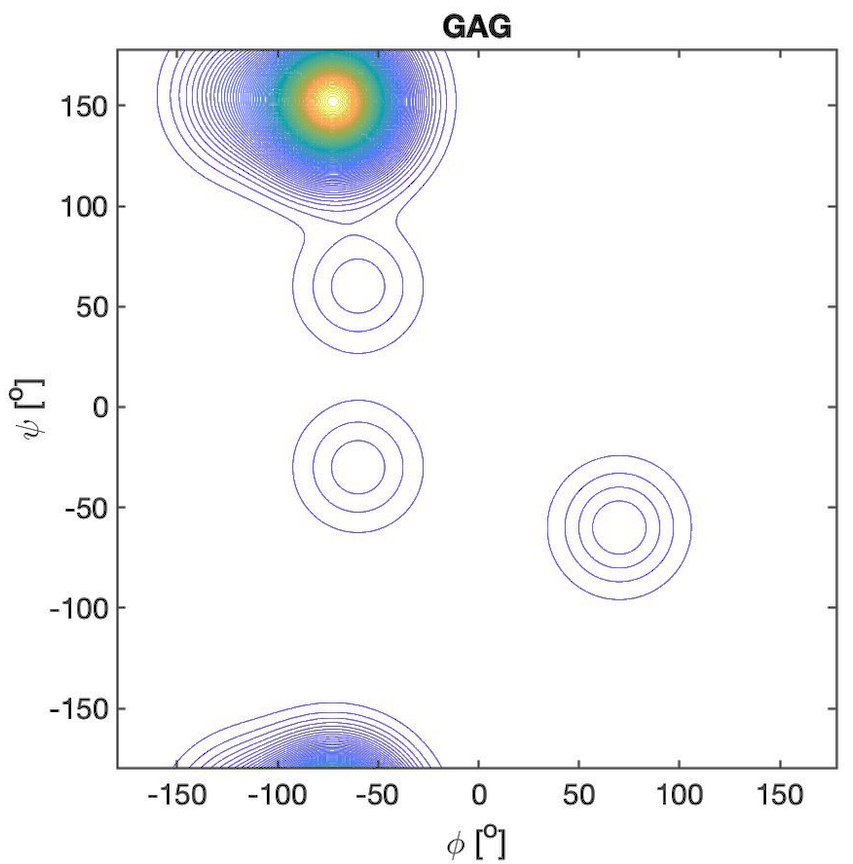

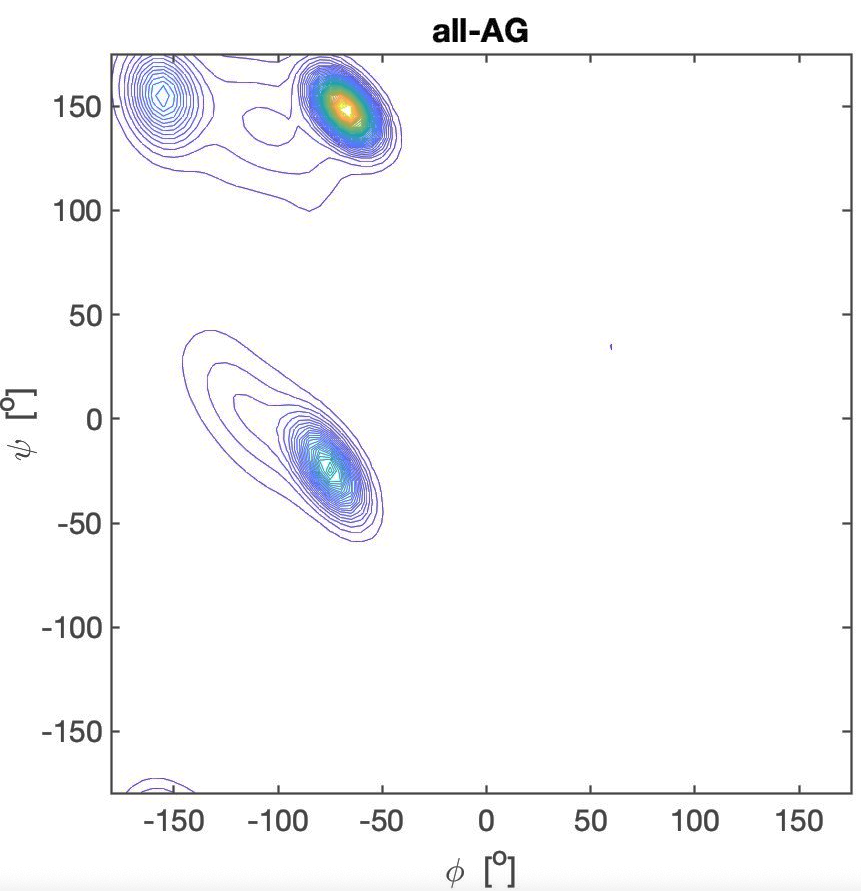

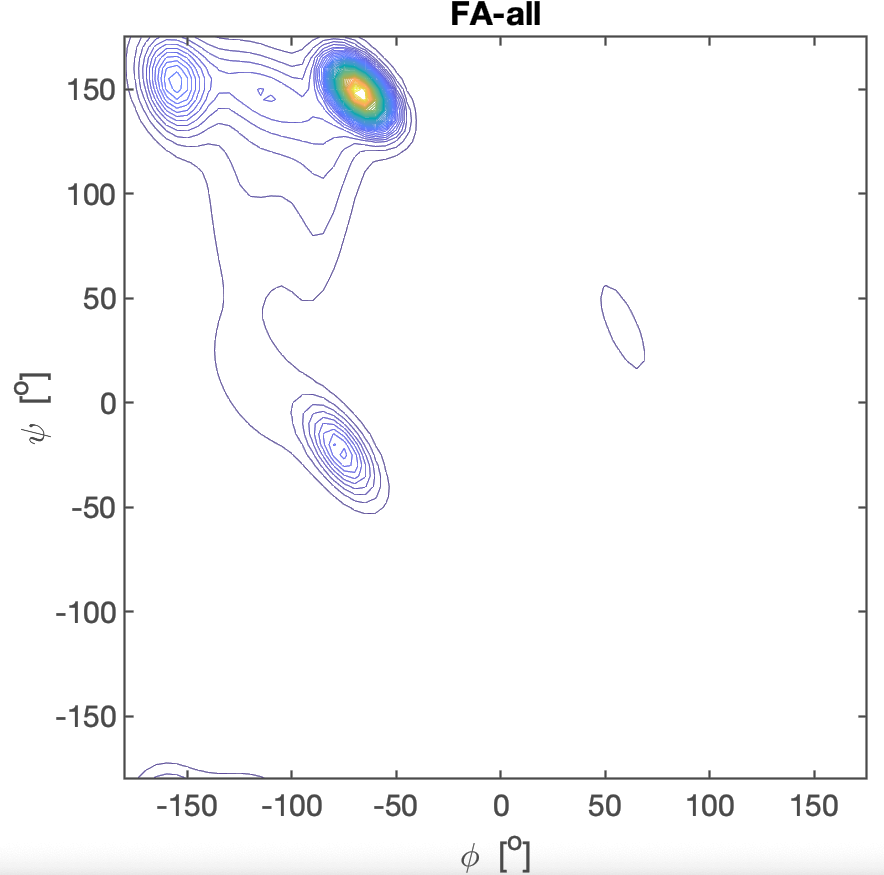

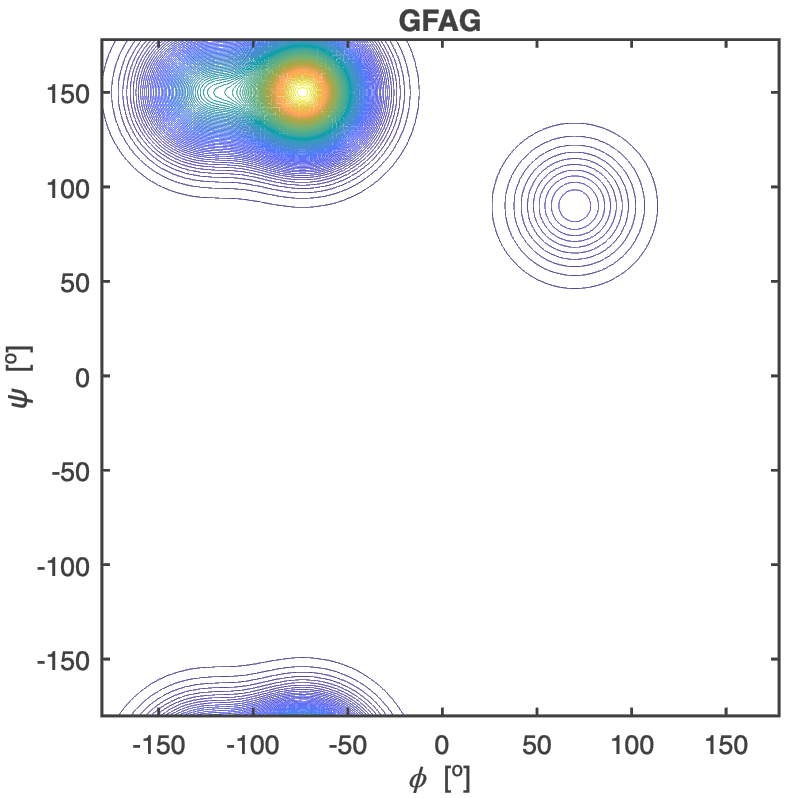

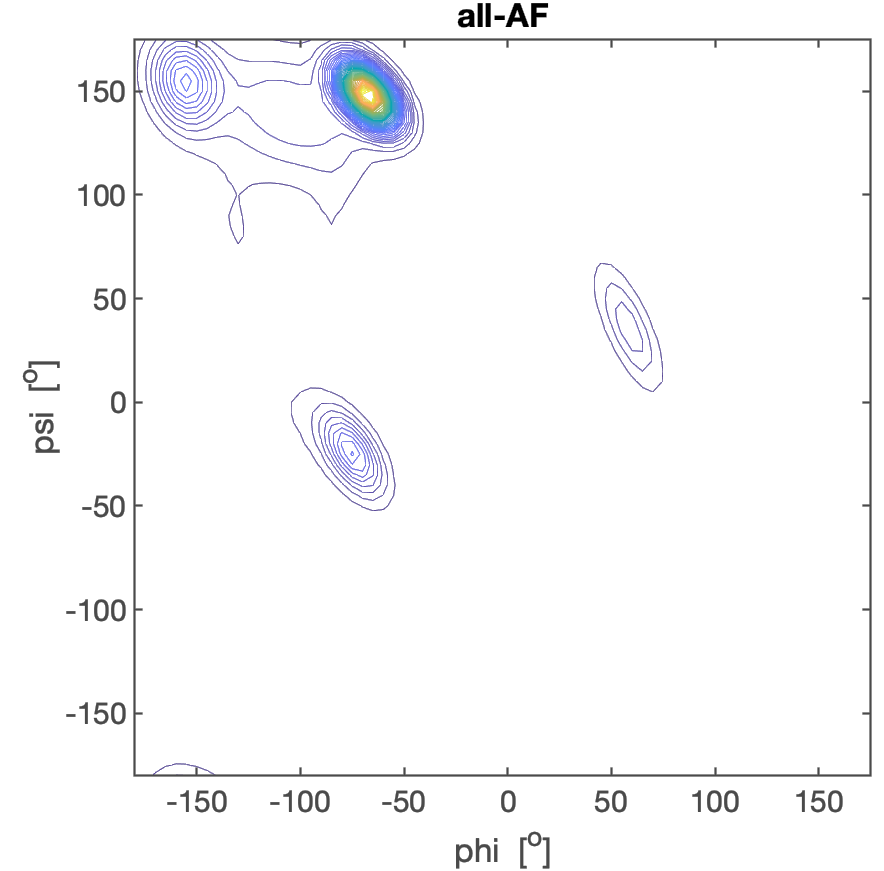

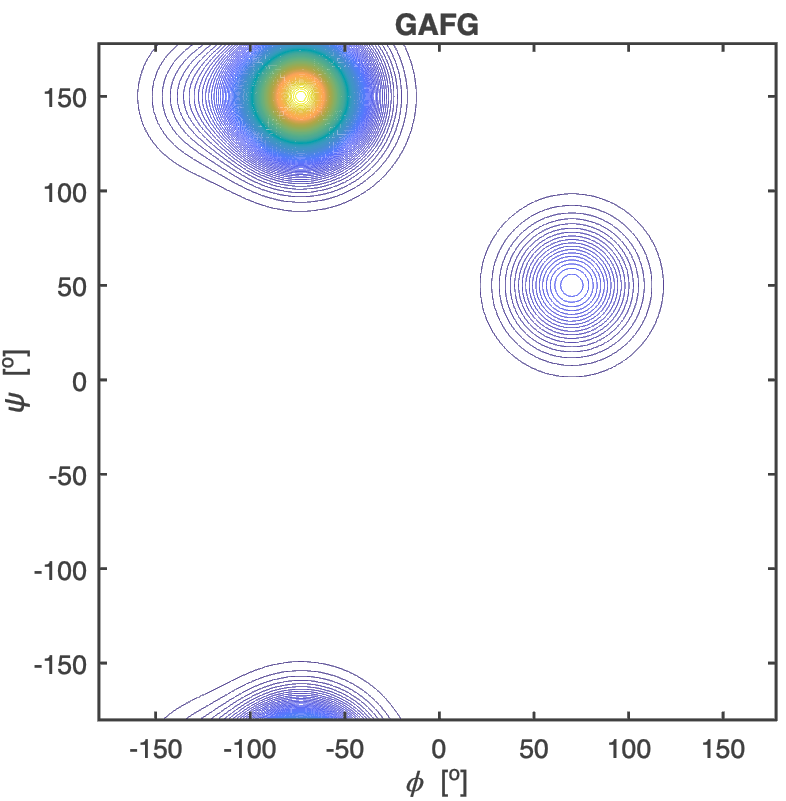

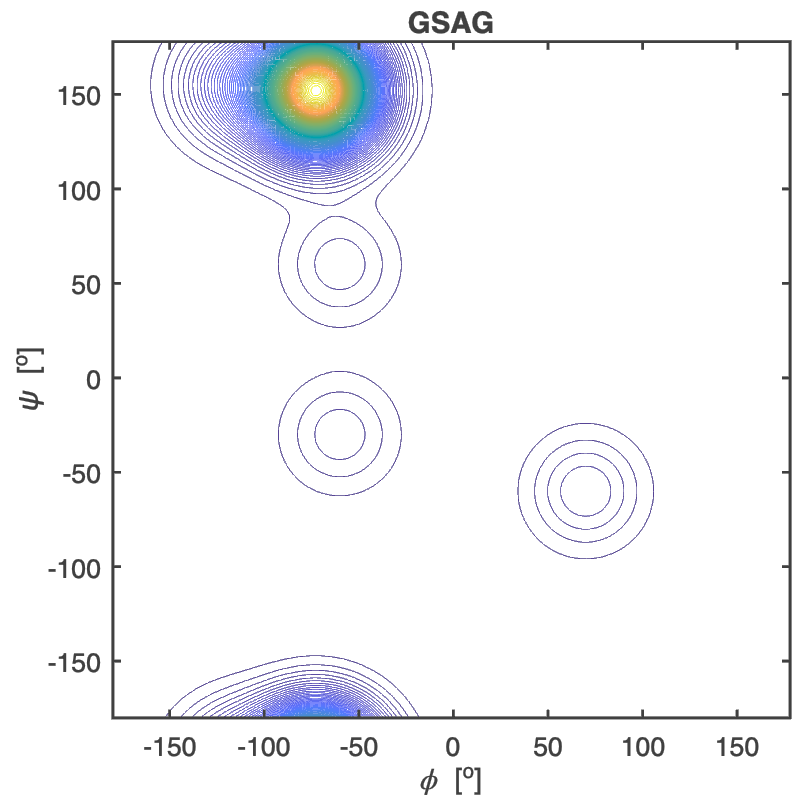

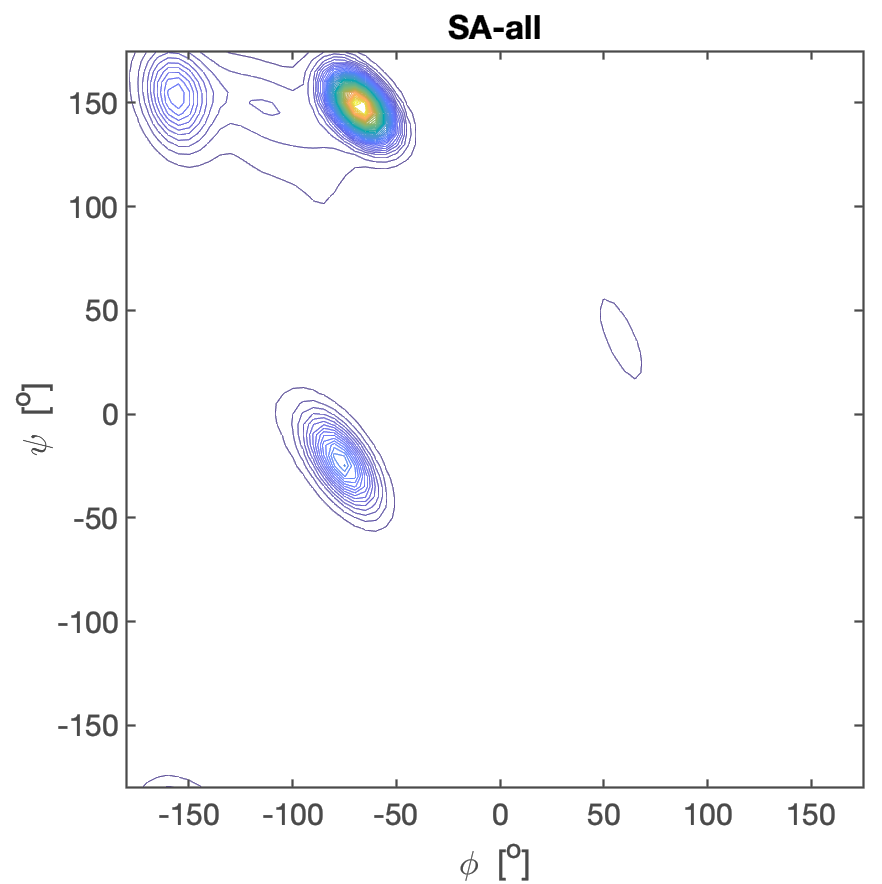

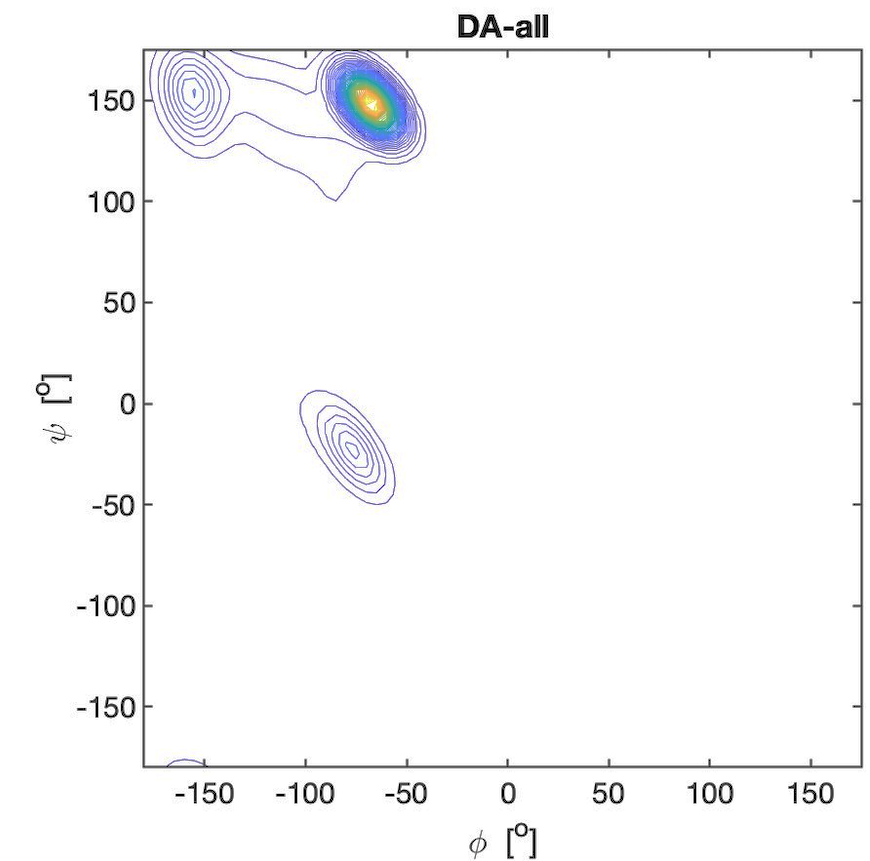

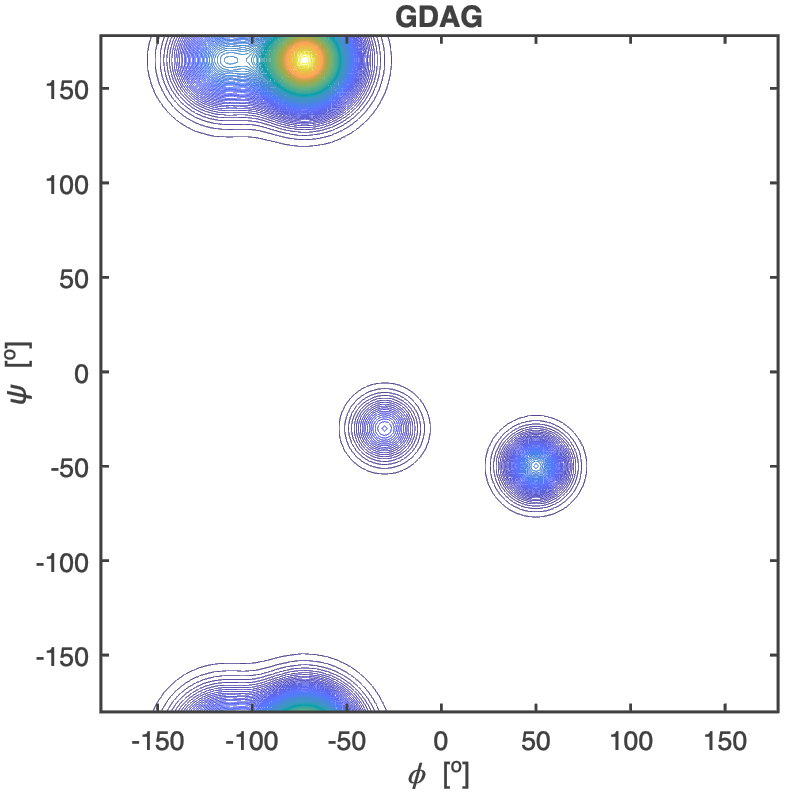

**Figure S10.** Ramachandran plots of alanine residues in GLG, all-**L**G, Gd**L**G, D**L**-all, Gs**L**G, S**L**-all, G**L**LG, all-**L**L, Gv**L**G, V**L**-all, Ga**L**G and A**L**-all. All explored peptides were in their cationic state. The plots for the peptides were constructed with a Gaussian model and the parameters listed in ref. for GLG,(12) in Table S2 for GD**L**G, GS**L**G, GL**L**G, GV**L**G and for GA**L**G. The plots for the coil library segments were built with files obtained from the website of the Dunbrack group with the
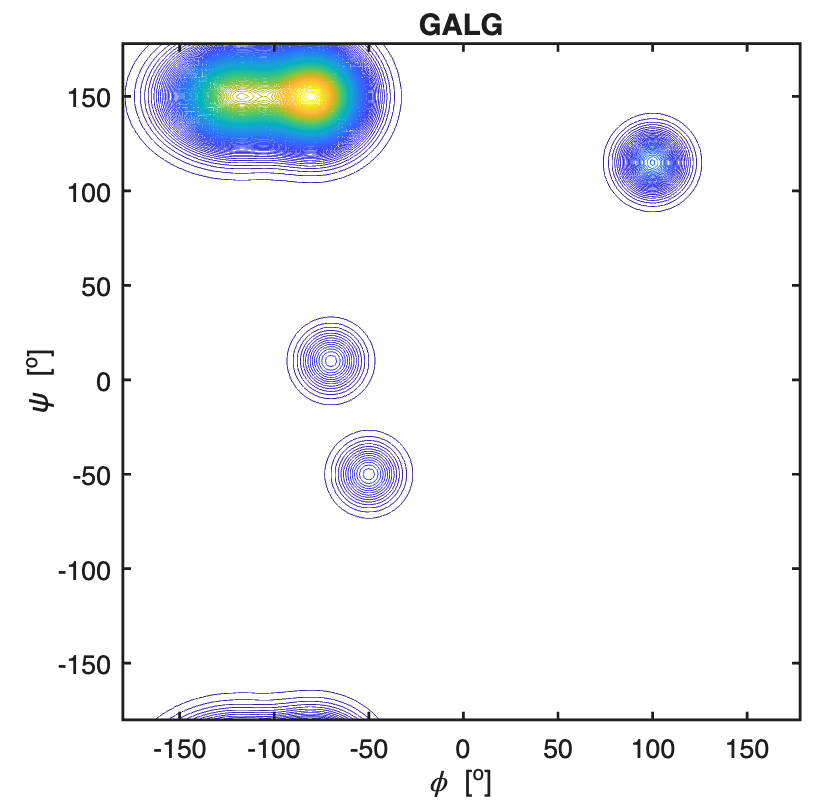

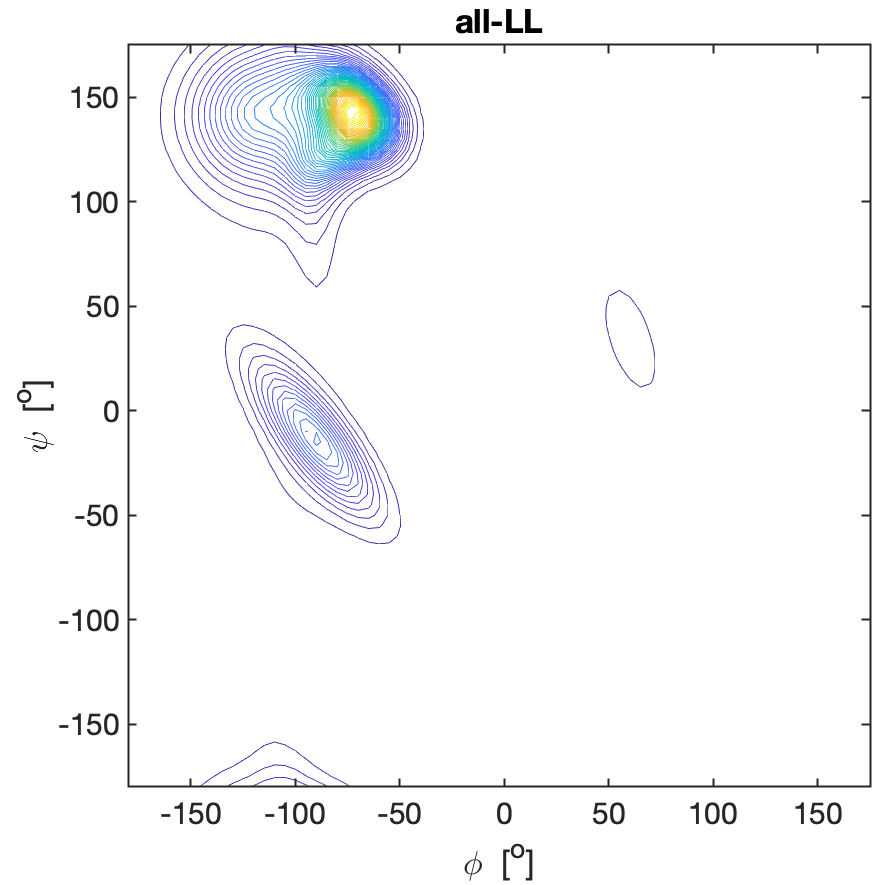

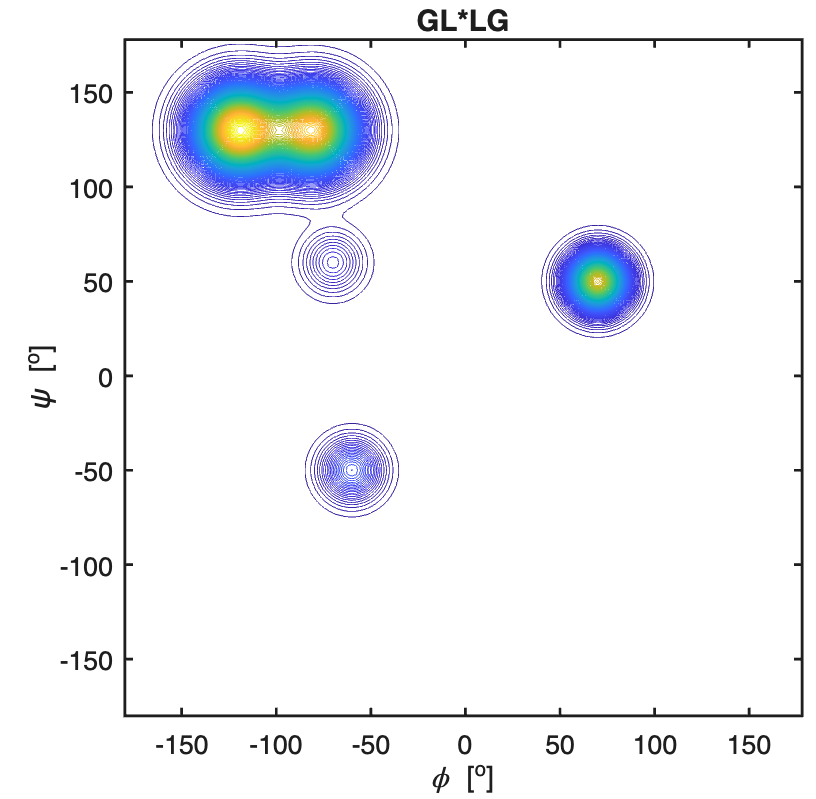

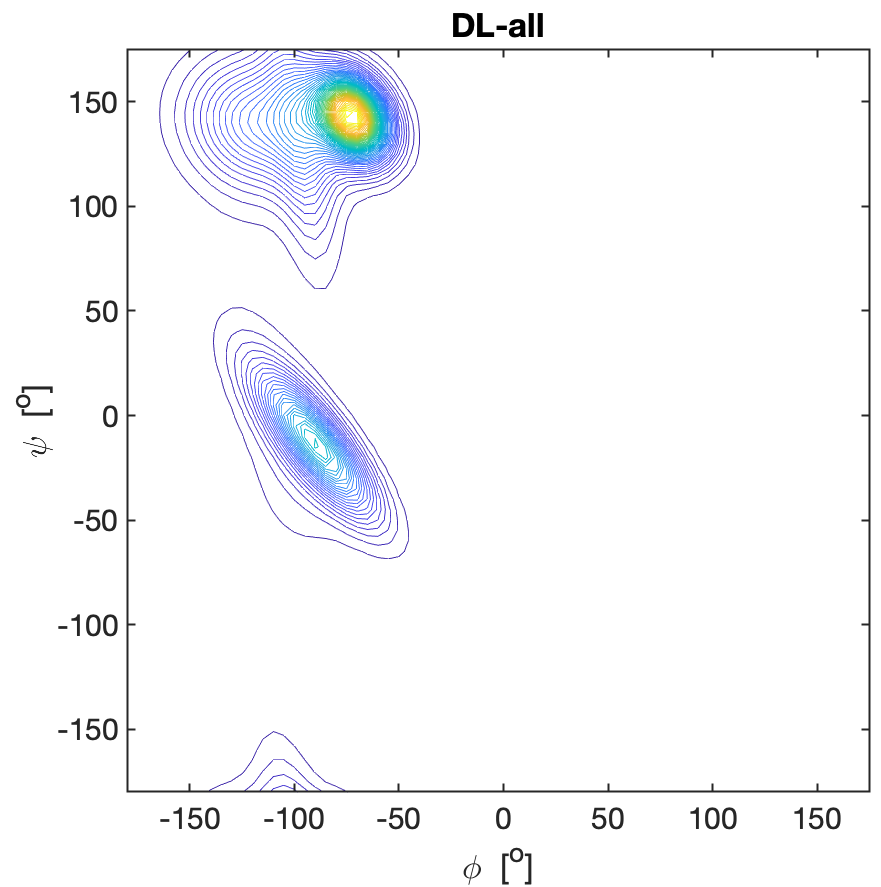

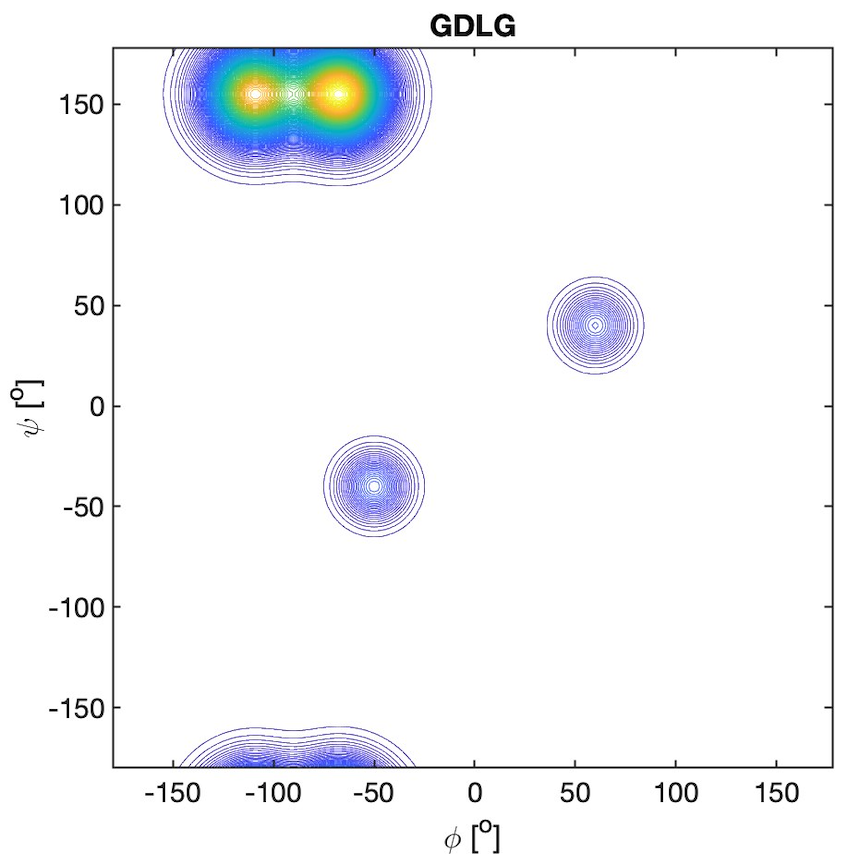

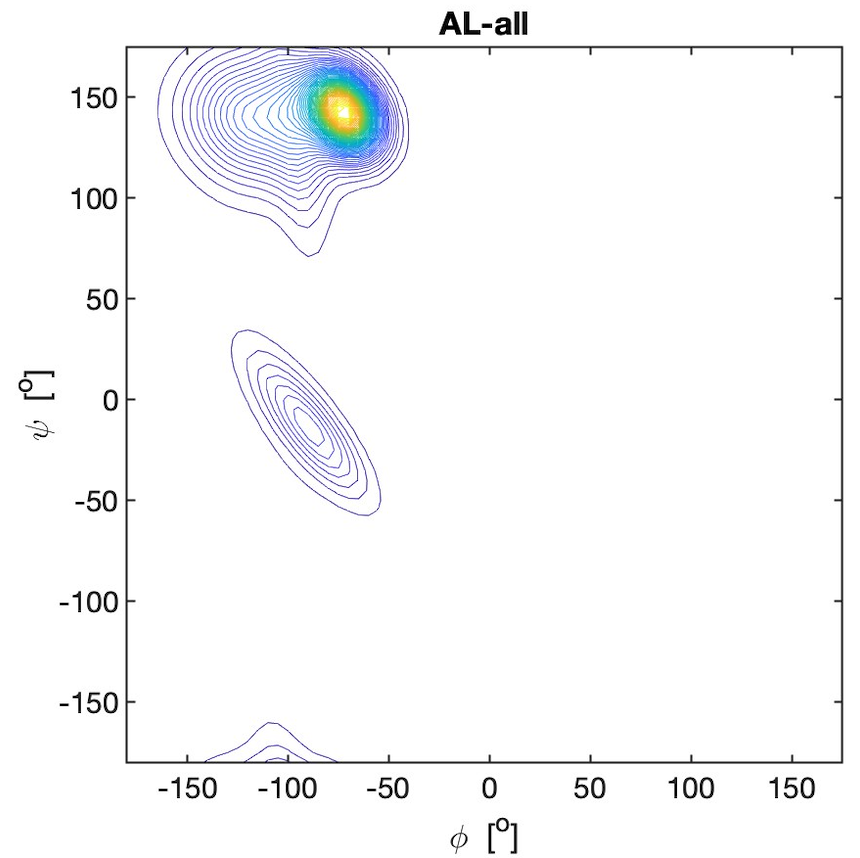

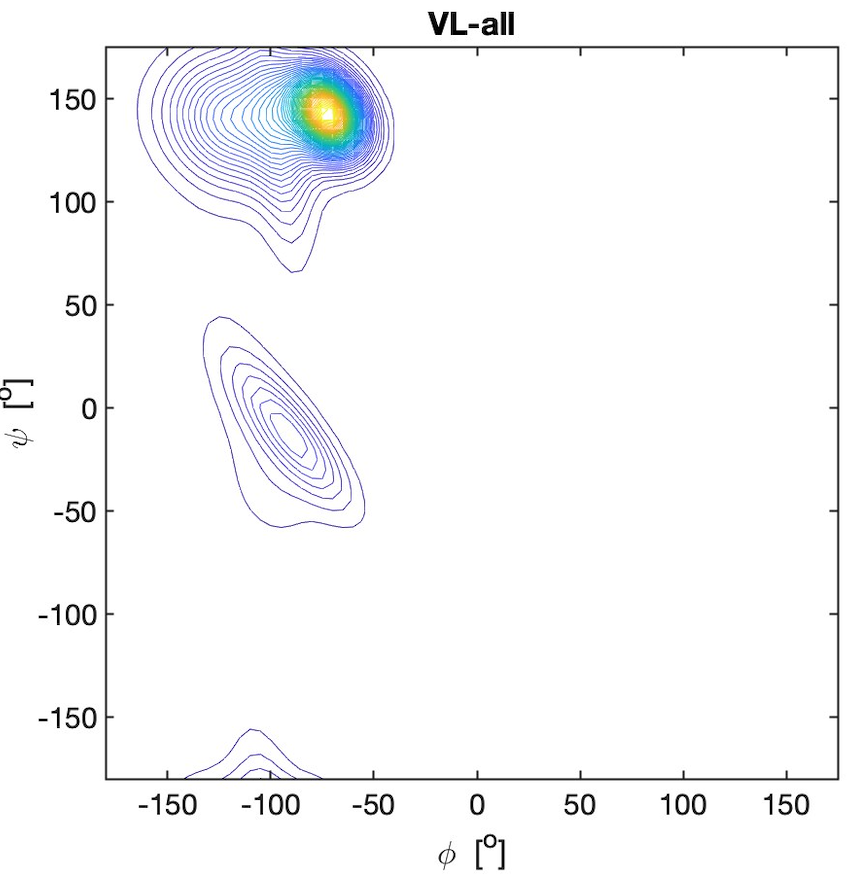

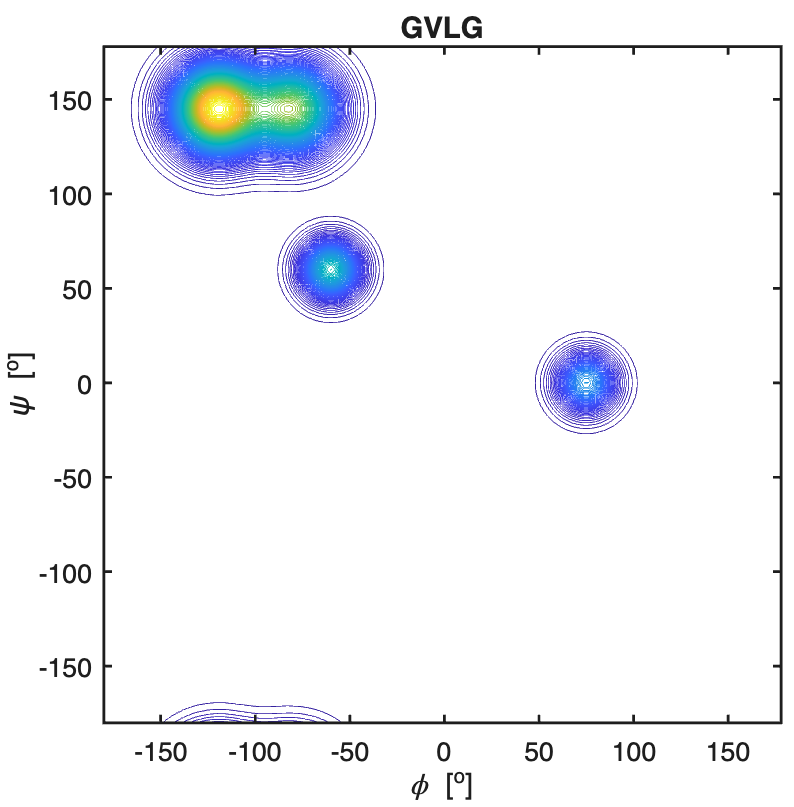

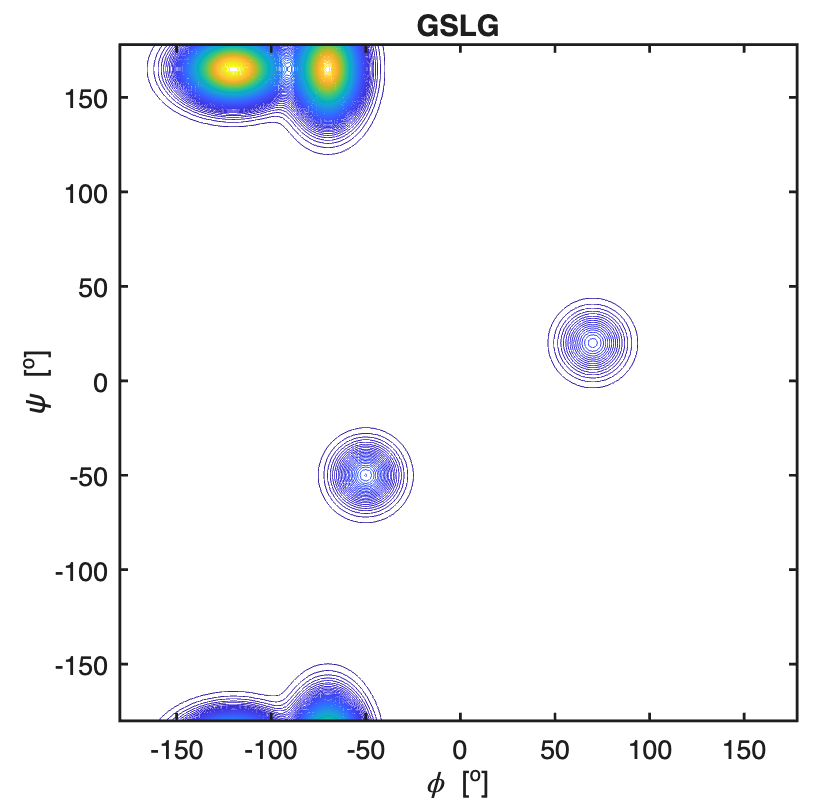

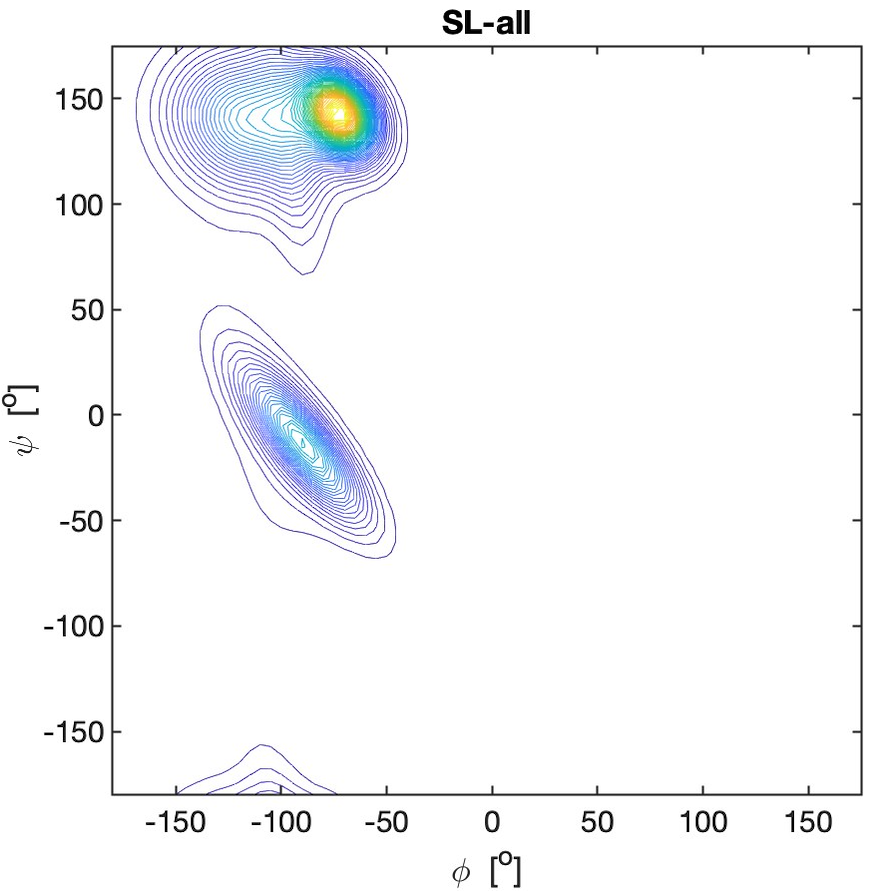

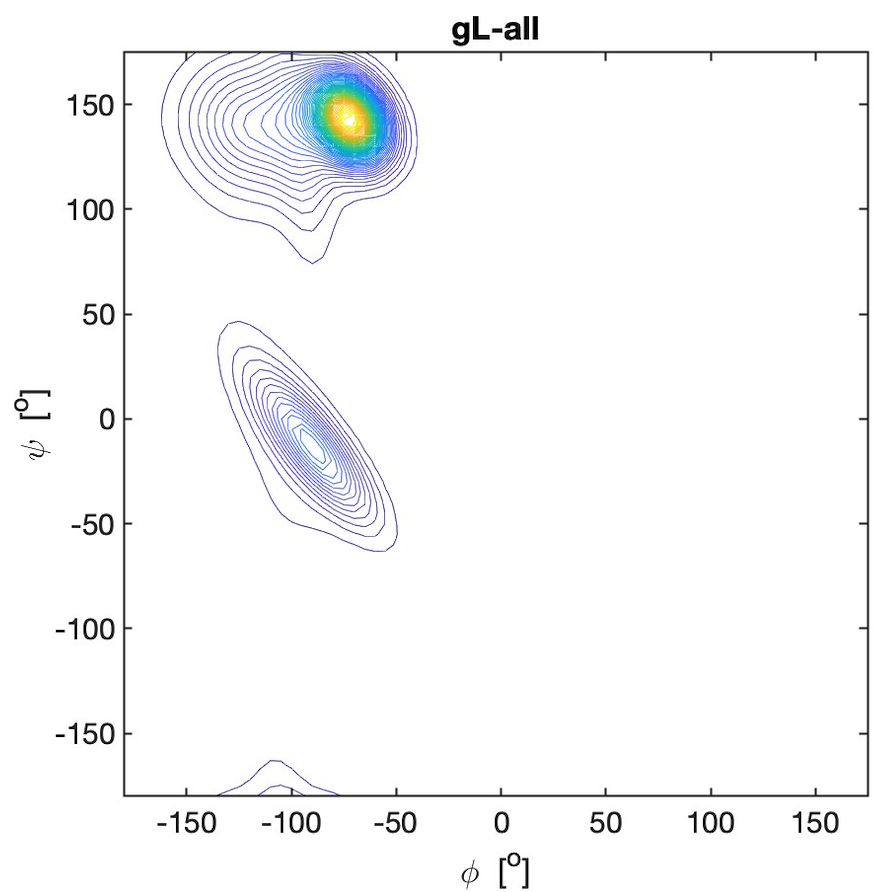
kind permission
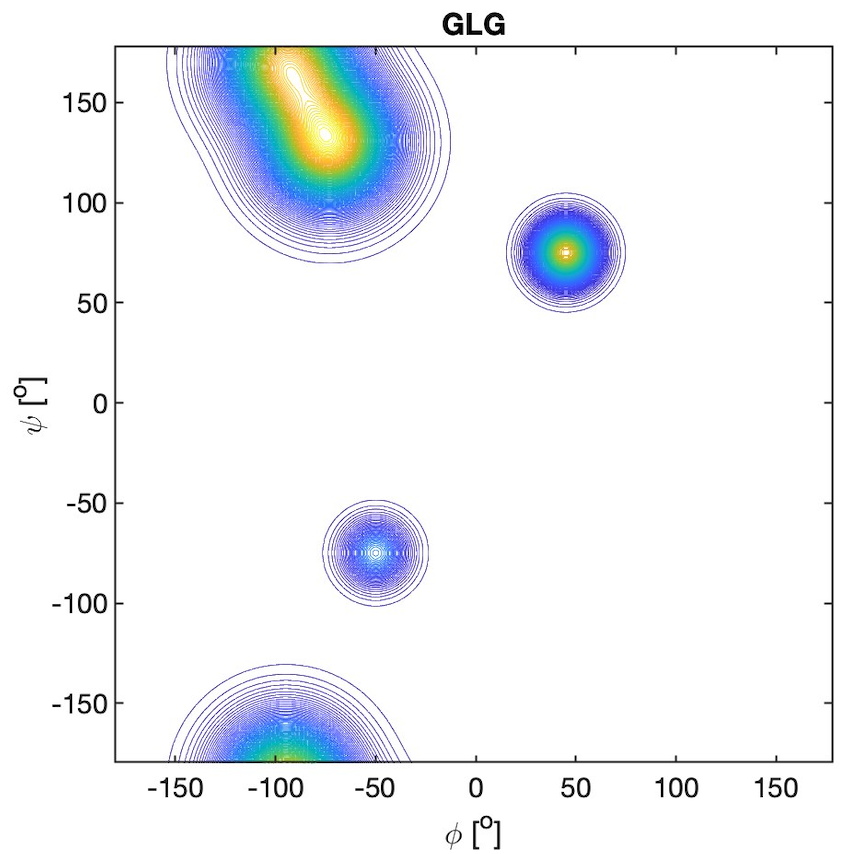
of Dr. Dunbrack.(13)

**Figure S11.** Ramachandran plots of aspartic acid residues in GDG, all-**D**G, G**D**AG, all-**D**A, G**D**KG, all-**D**K, G**D**LG, all-**D**L, G**D**FG, all-**D**F, GF**D**G, F**D**-all, G**D**VG, and all-**D**V, Corresponding plots for aspartic acid as upstream and downstream neighbor of aspartic acid are shown in Figure 2 of the main manuscript. All explored peptides were in their cationic state. The plots for the peptides were constructed with a Gaussian model and the parameters listed in ref.(6) for GDG, in Table S2 for G**D**AG, G**D**KG, G**D**LG, and G**D**VG and in ref.(6) for G**D**FG and GF**D**G. The plots for the coil library segments were built with files obtained from the website of the Dunbrack group with the kind
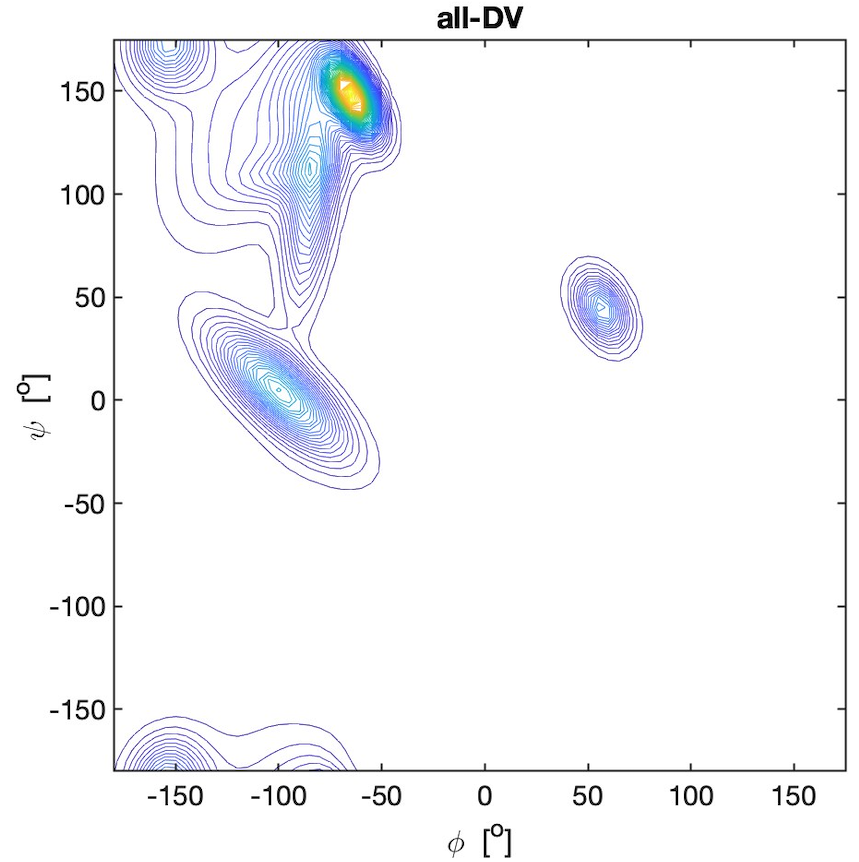

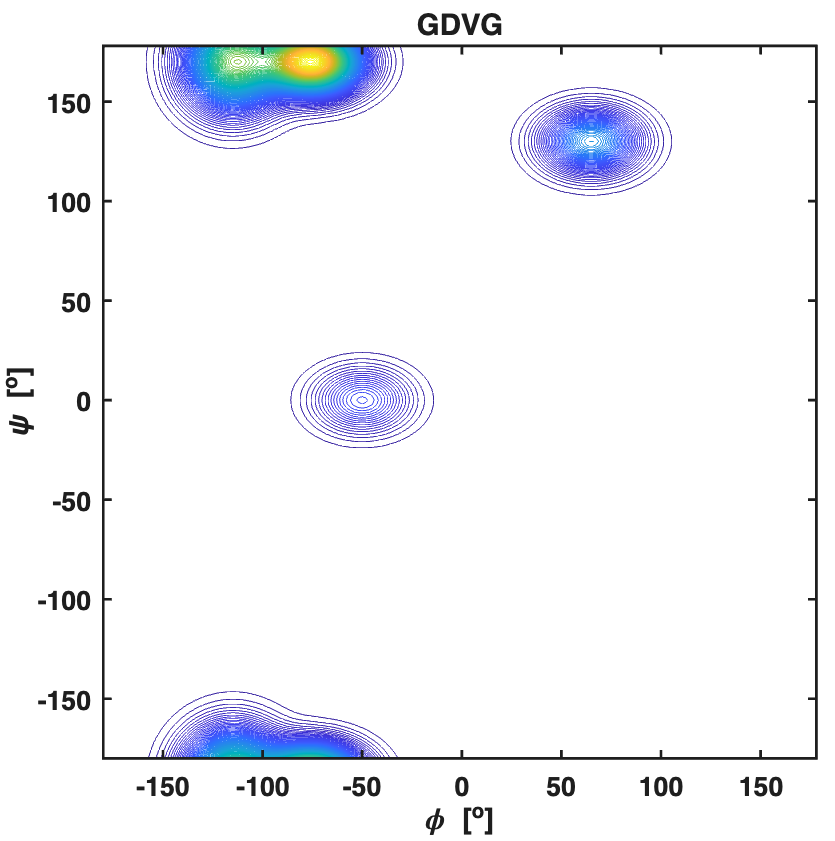

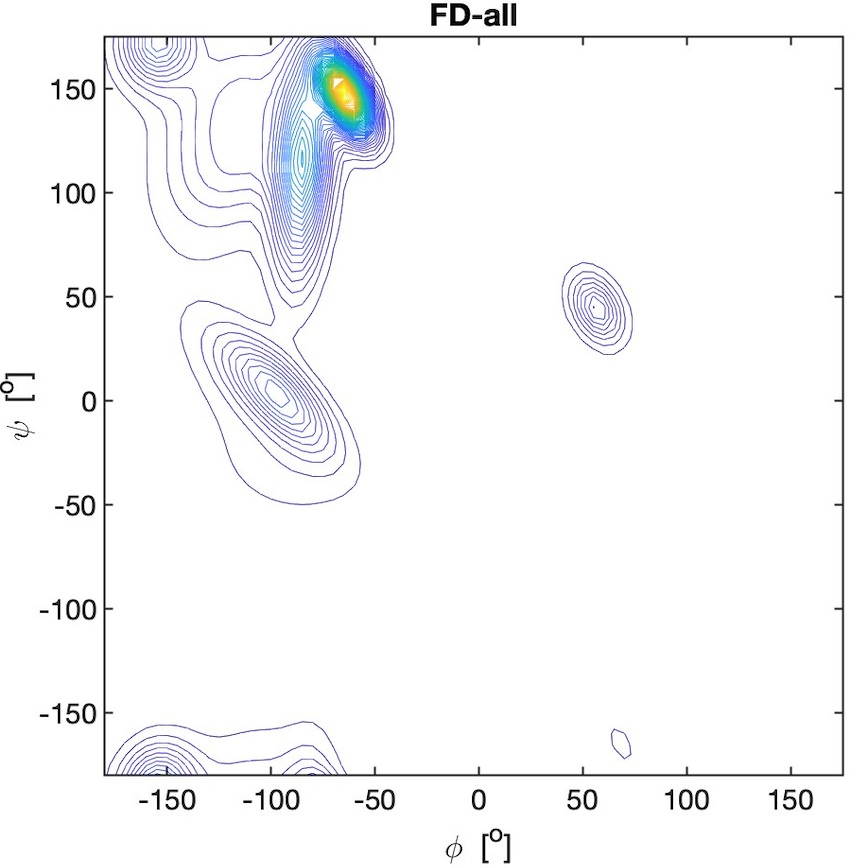

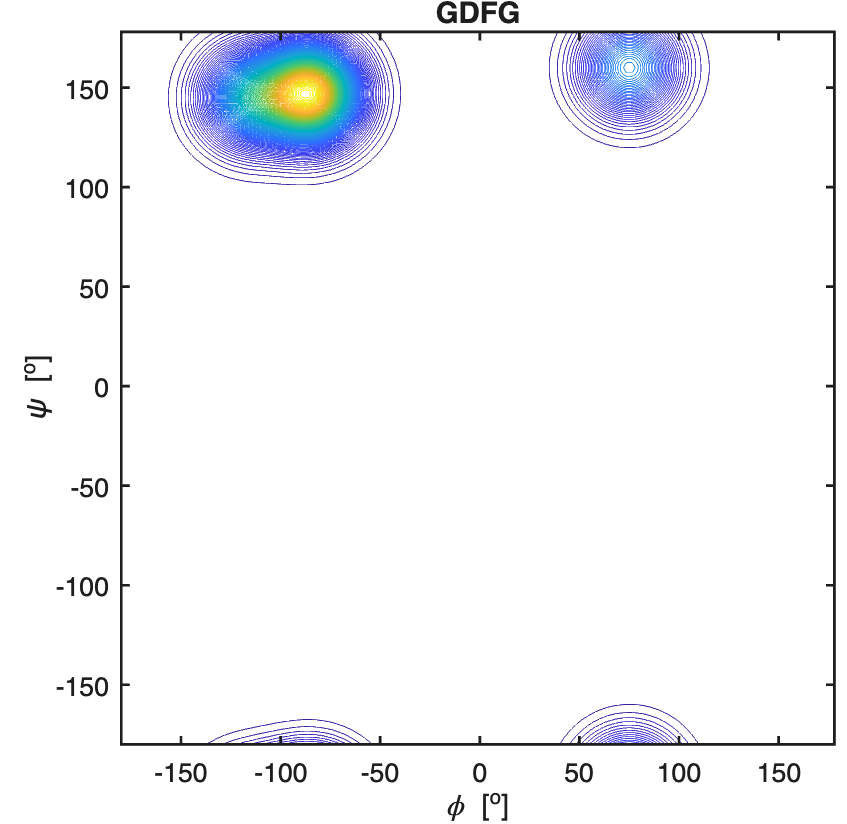

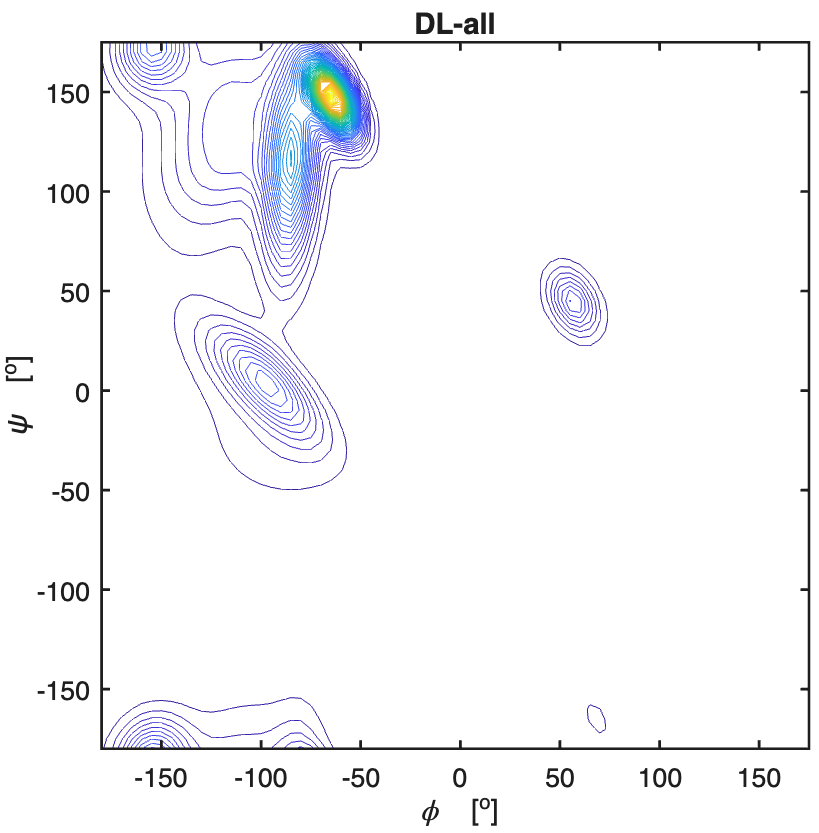

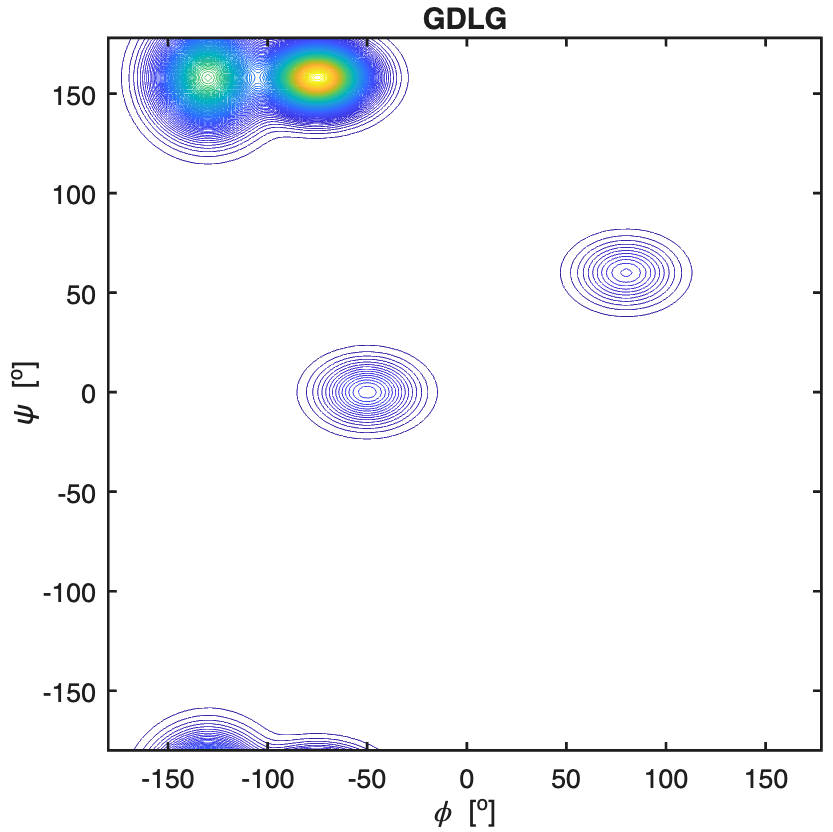

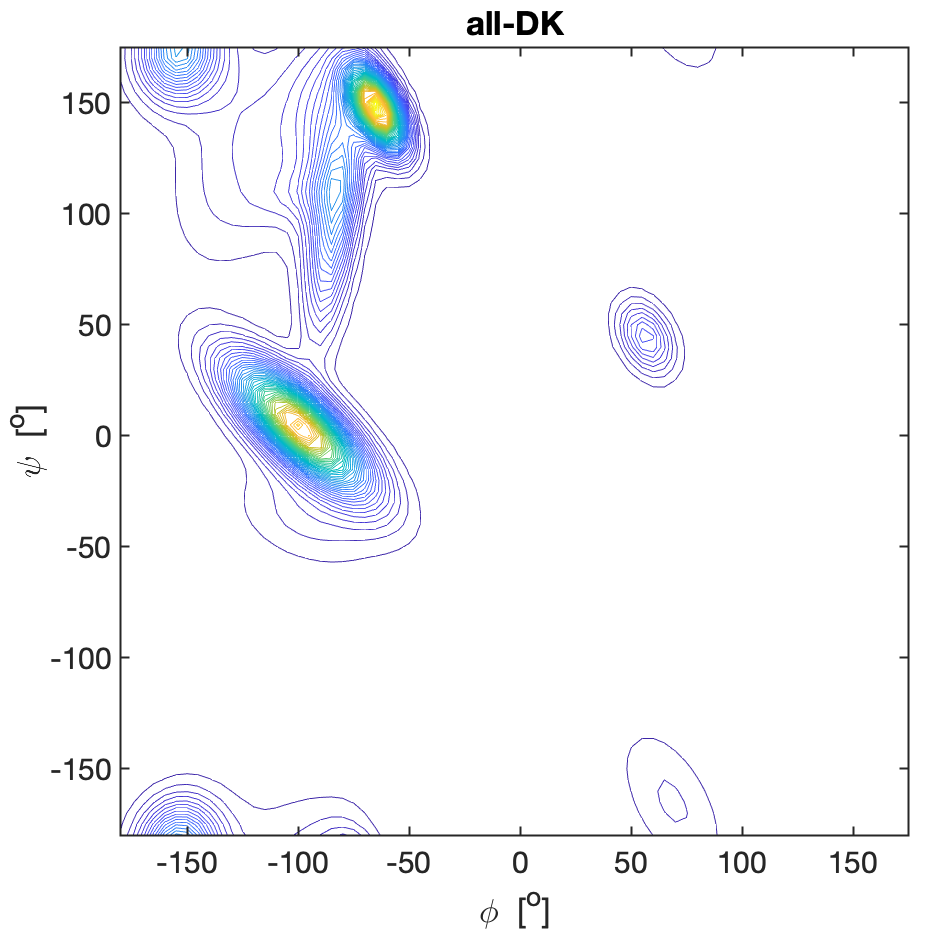

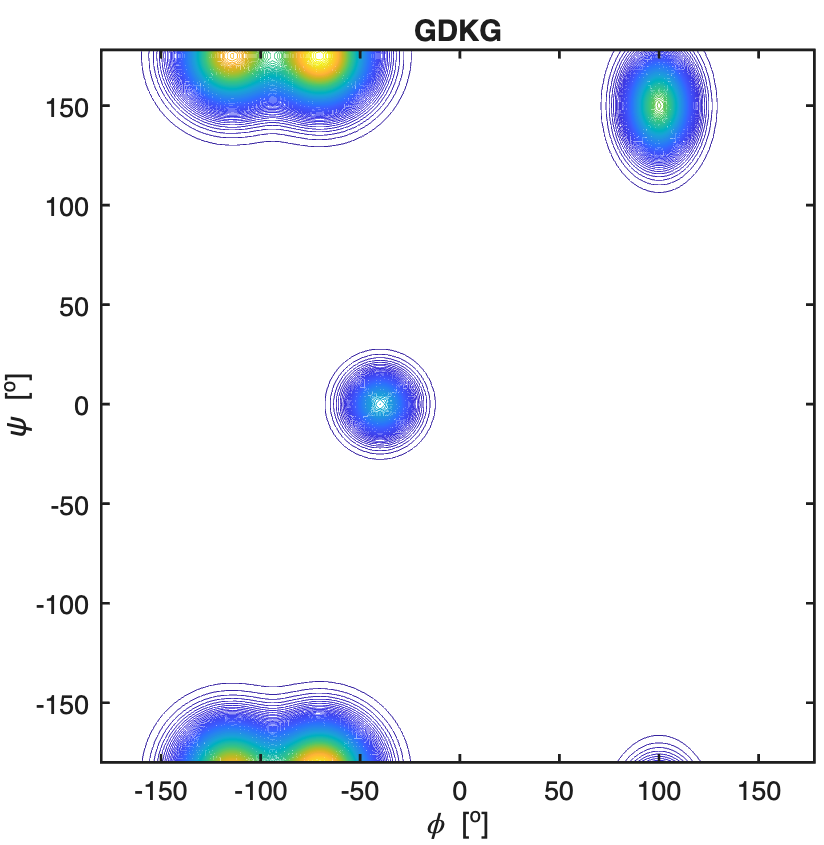

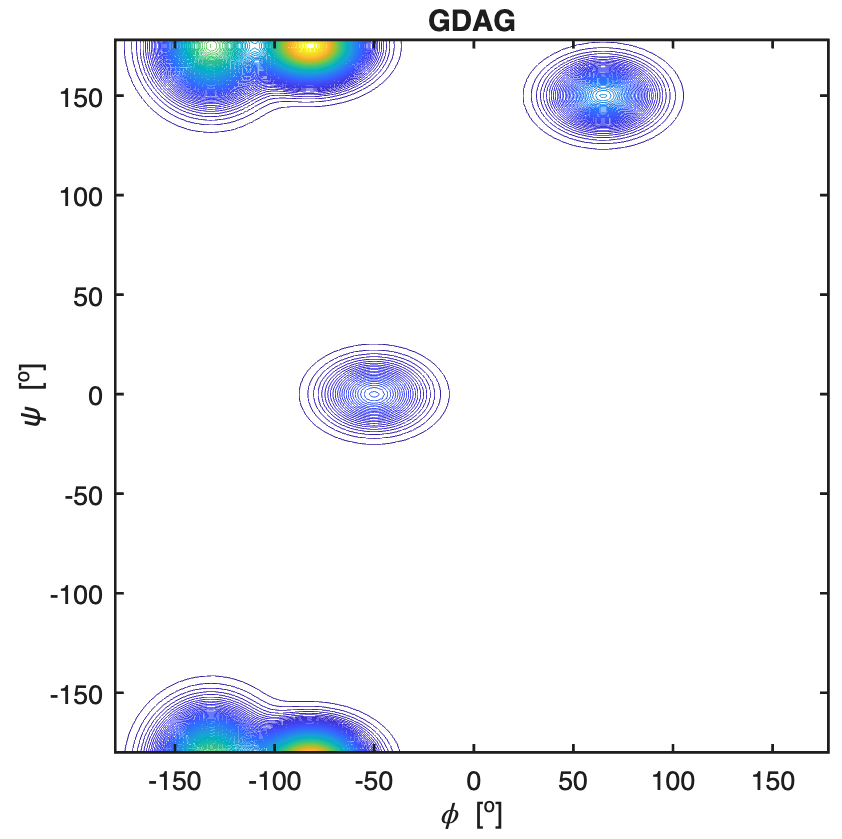

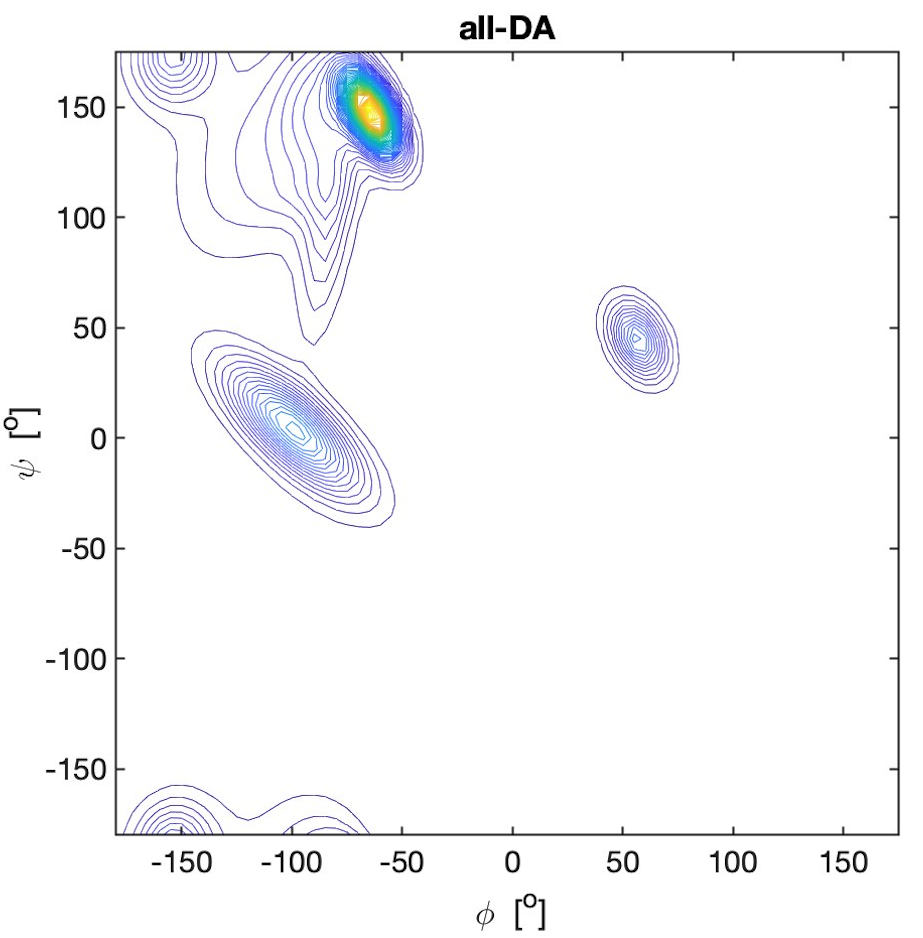

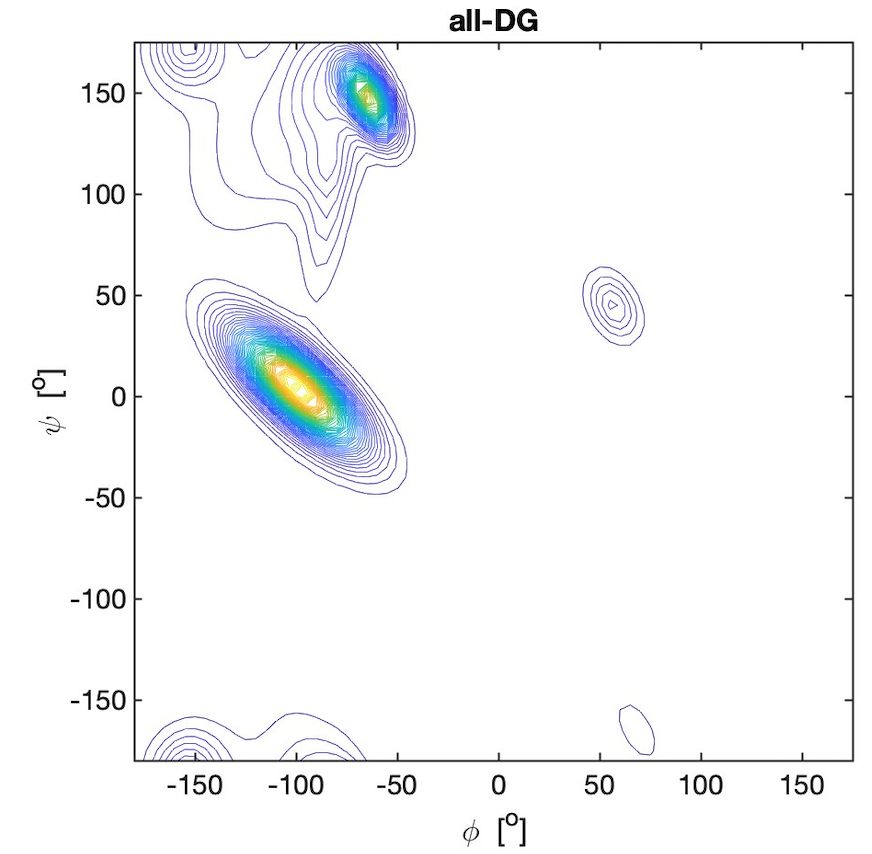
permission of Dr.
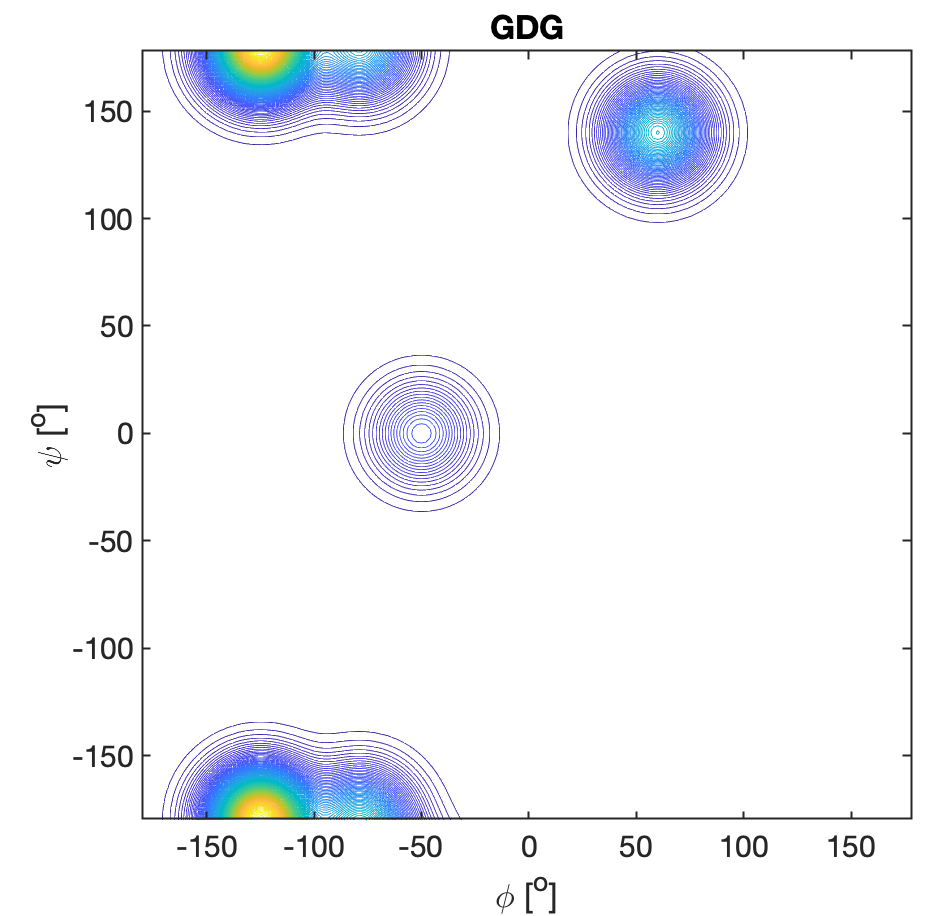
Dunbrack.(13)

**Figure S12.** Ramachandran plots of serine residues in GSG, all-**S**G, G**S**AG, all-**S**A, G**S**KG, all-**S**K, G**S**LG, all-**S**L, G**S**VG, and all-**S**V. All explored peptides were in their cationic state. The plots for the peptides were constructed with a Gaussian model and the parameters listed in ref.(12) for GSG and in Table S2 for the tetra-peptides. The plots for the coil library segments were built with files obtained from the website of the Dunbrack group with the kind permission of Dr.
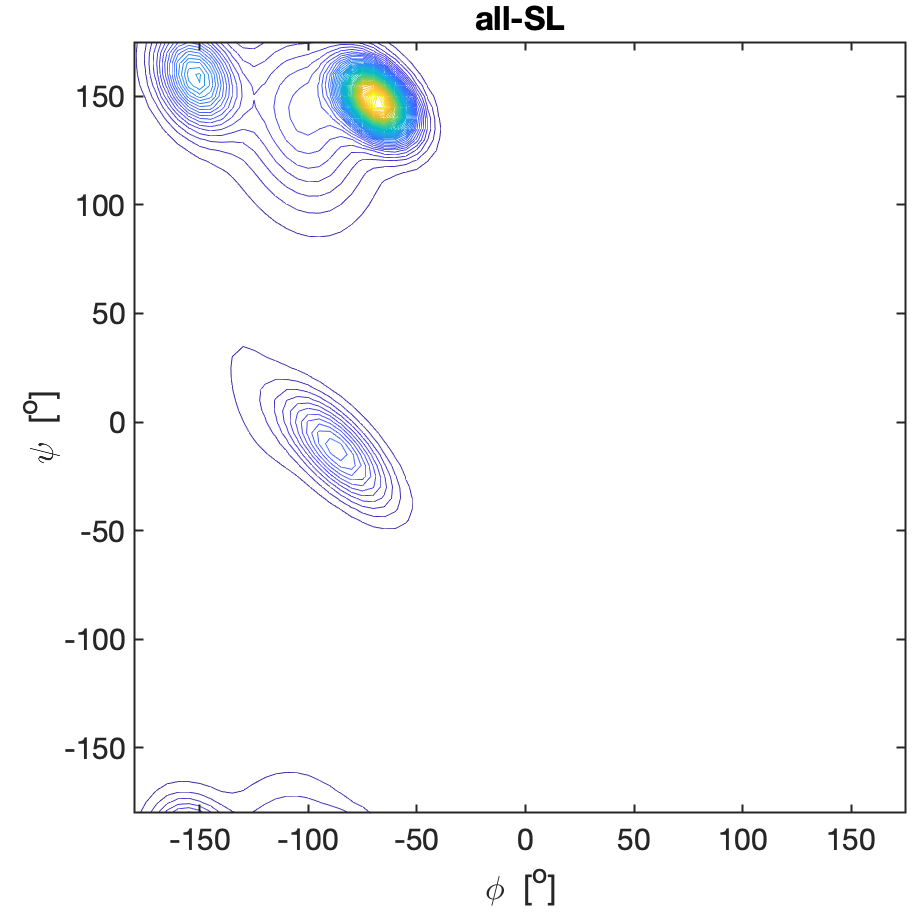

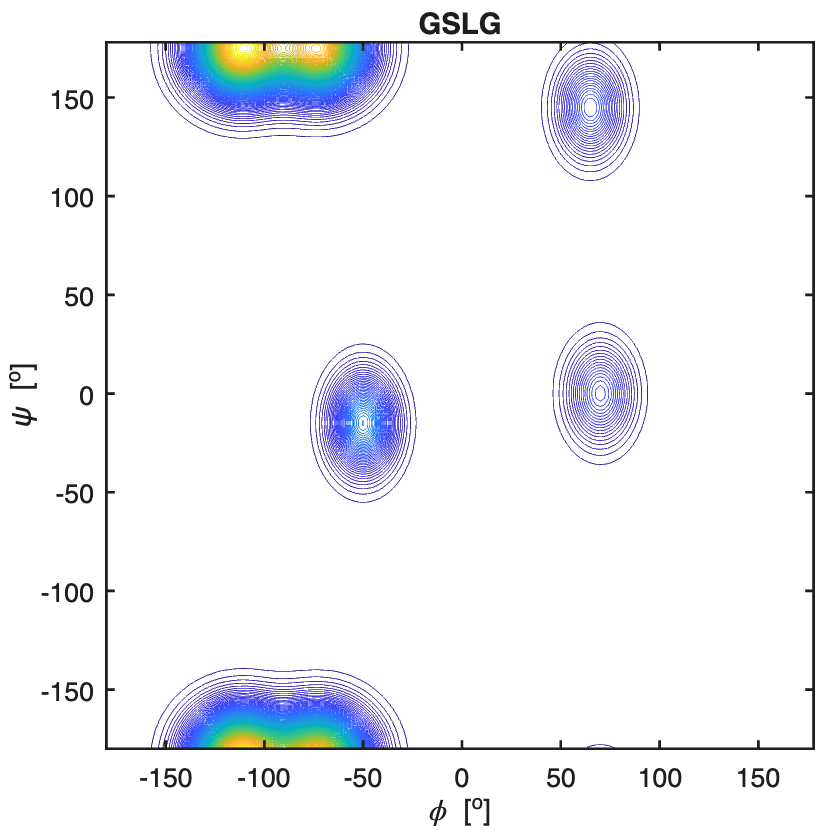

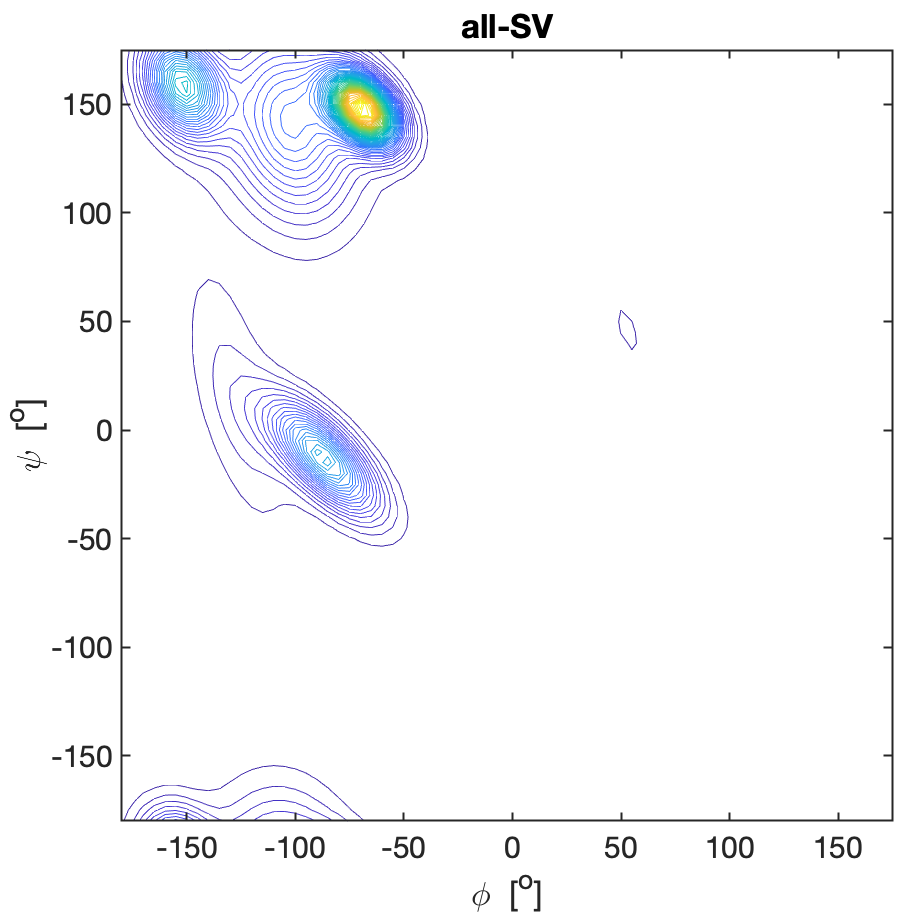

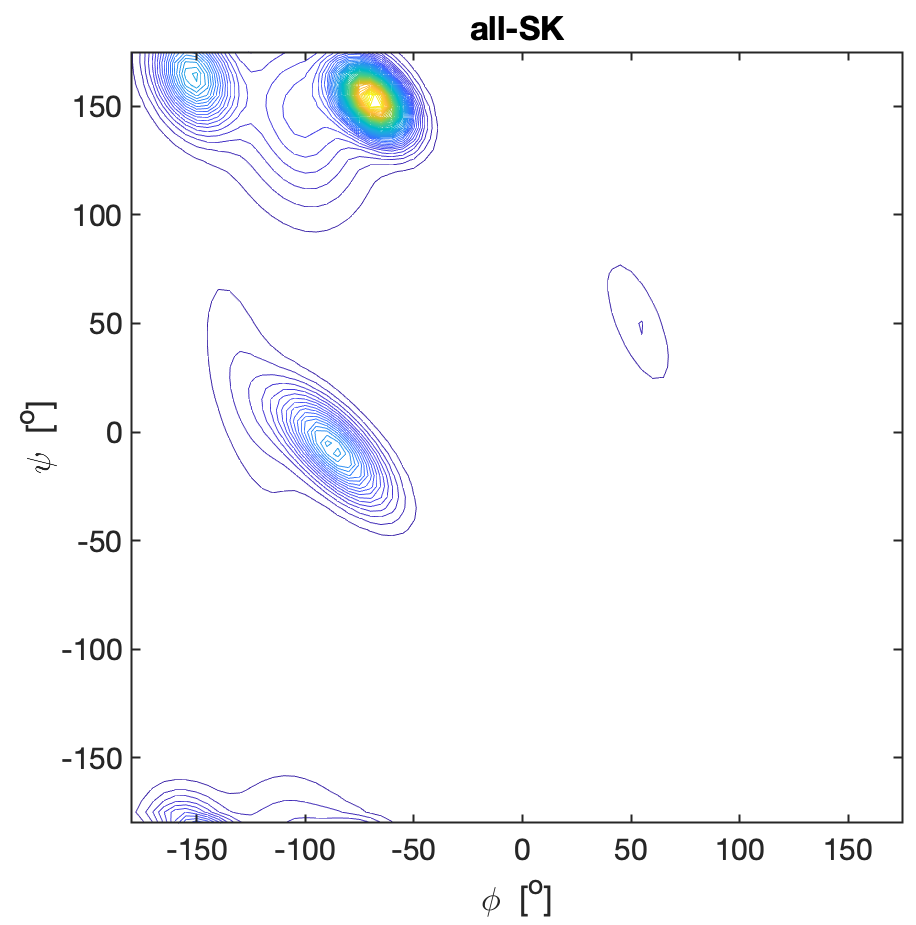

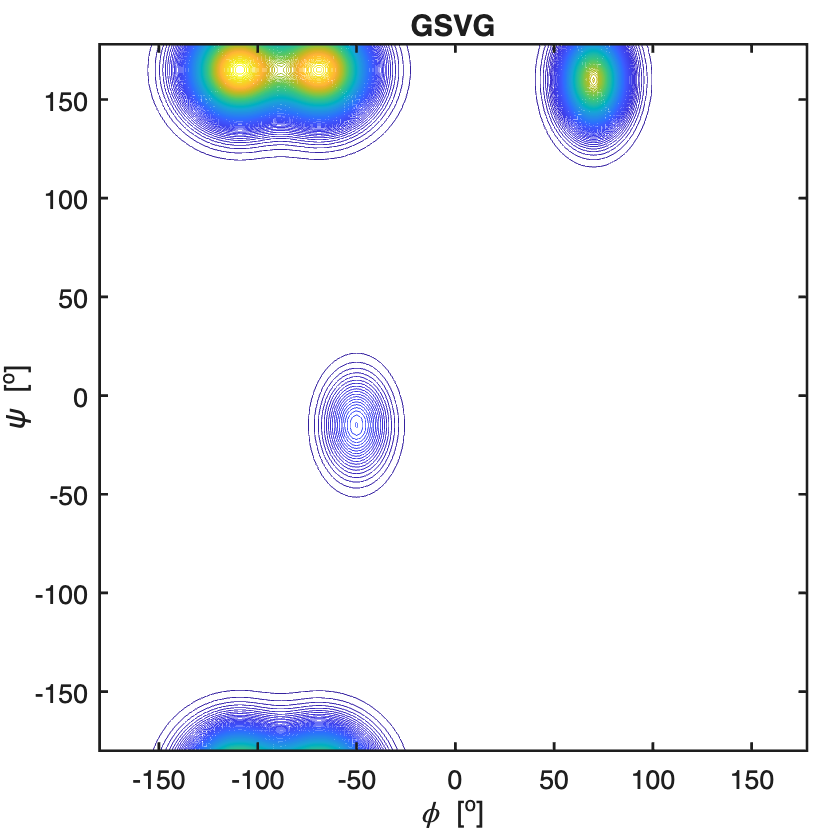

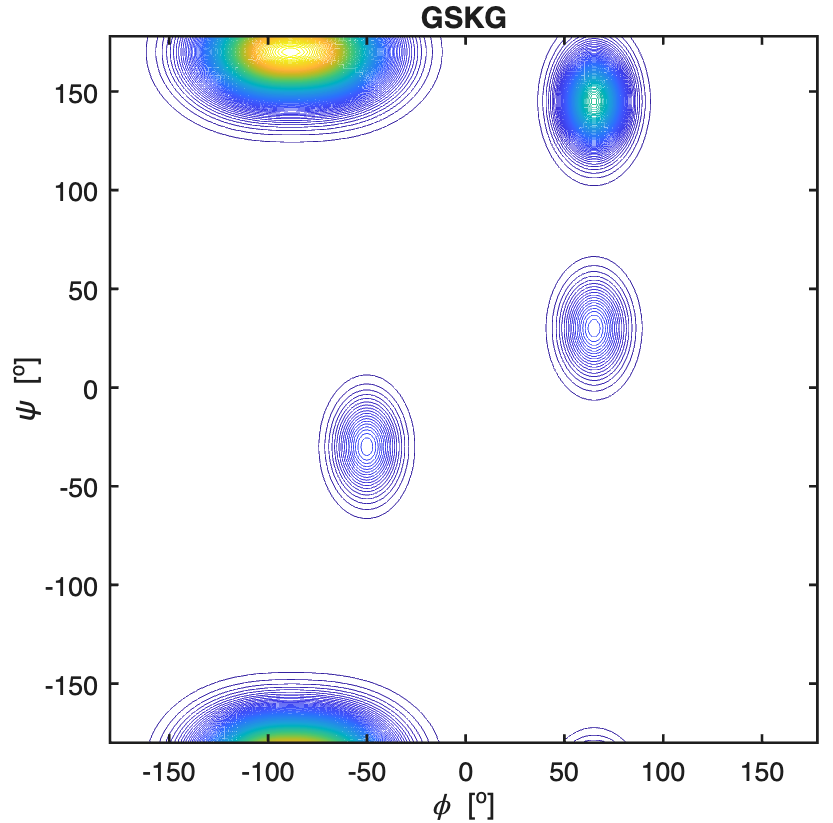

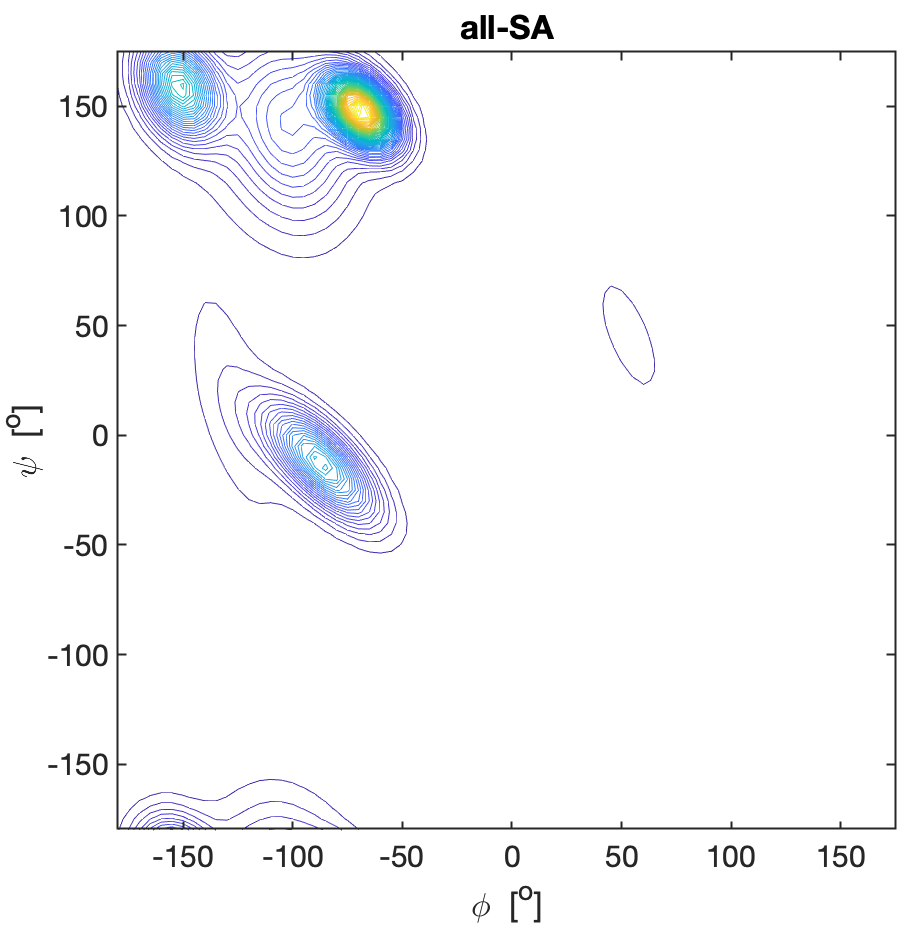

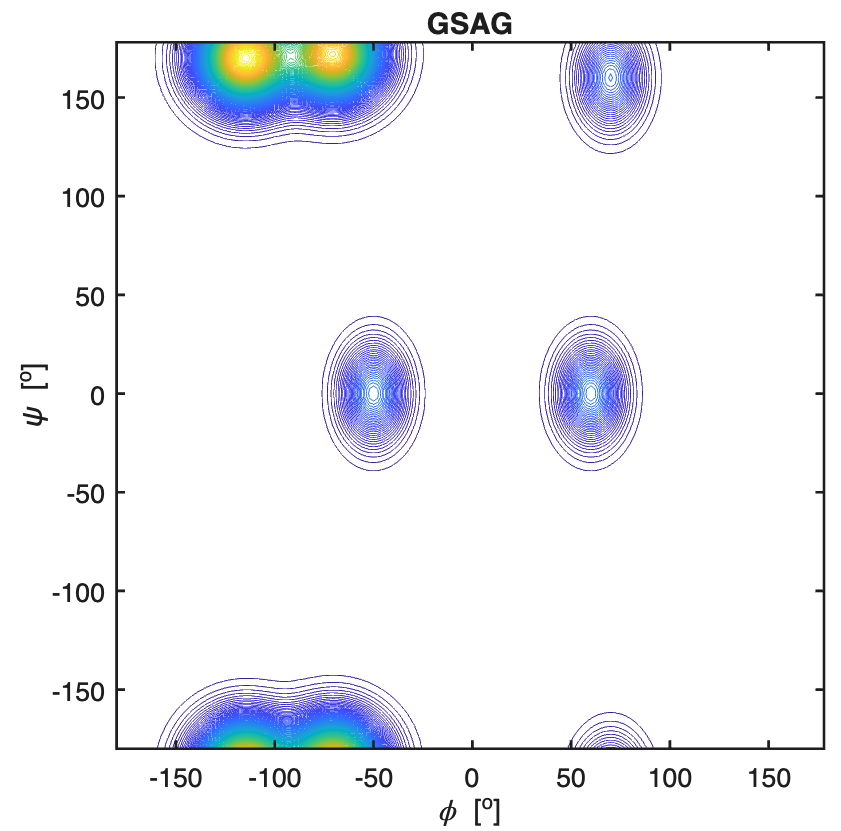

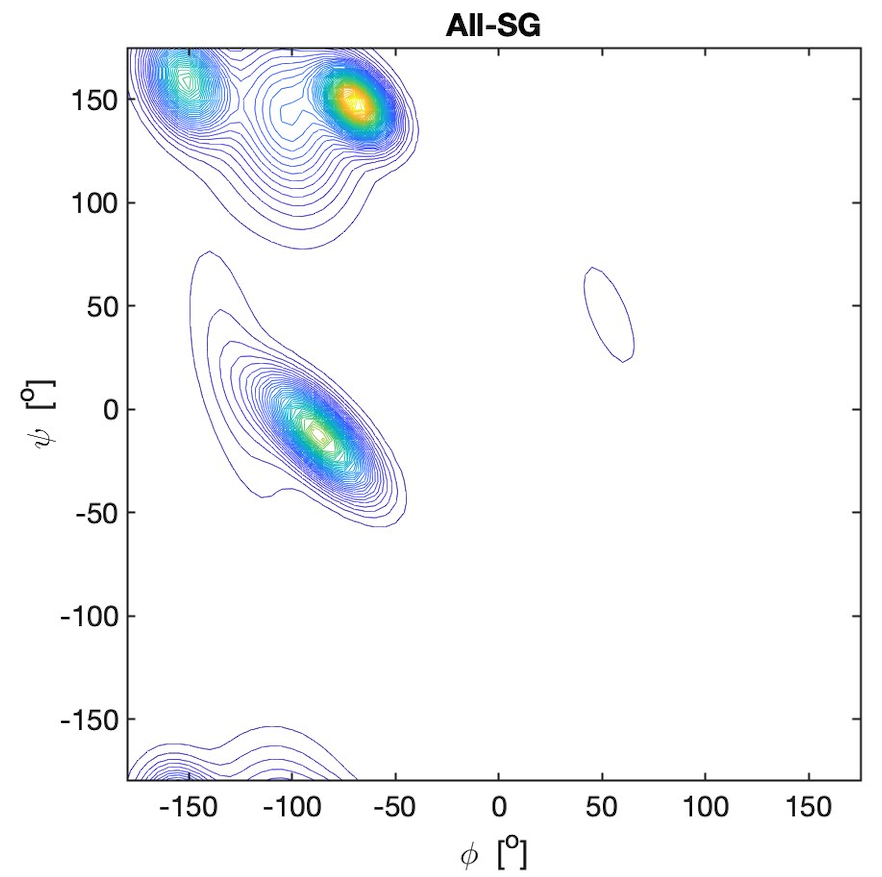

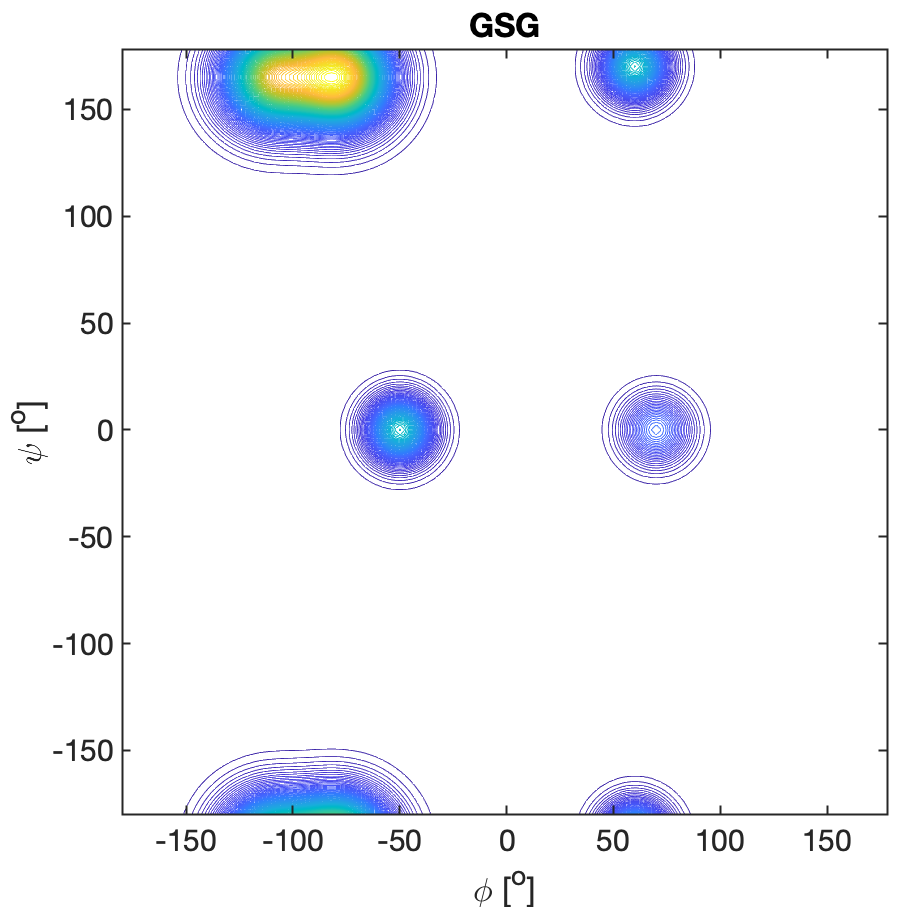
Dunbrack.(13)

**Figure S13.** Ramachandran plots of lysine residues in GKG, all-**K**G, GD**K**G, D**K**-all, GS**K**G, S**K**-all, G**K**LG, all-KL, G**K**VG, and all-**K**V. All explored peptides were in their cationic state. The plots for the peptides were constructed with a Gaussian model and the parameters listed in ref.(12) for GKG and in Table S2 for the tetra-peptides. The plots for the coil library segments were built with files obtained from the website of the Dunbrack group with the kind permission of Dr.
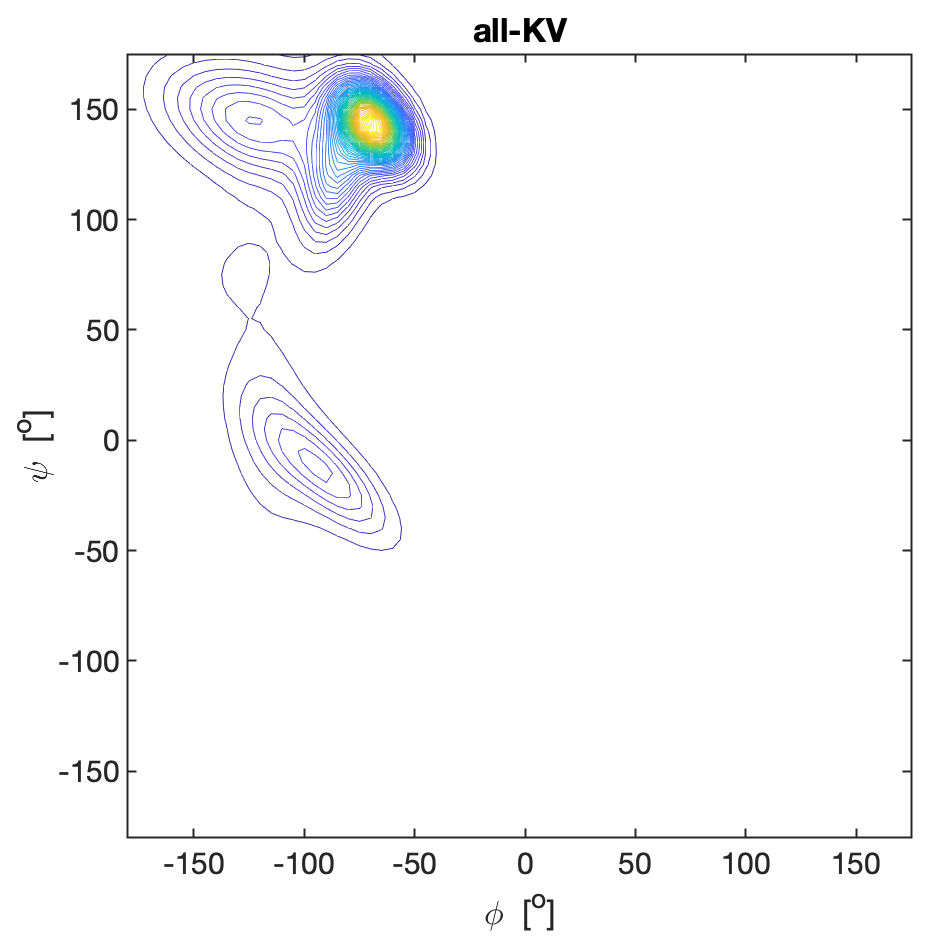

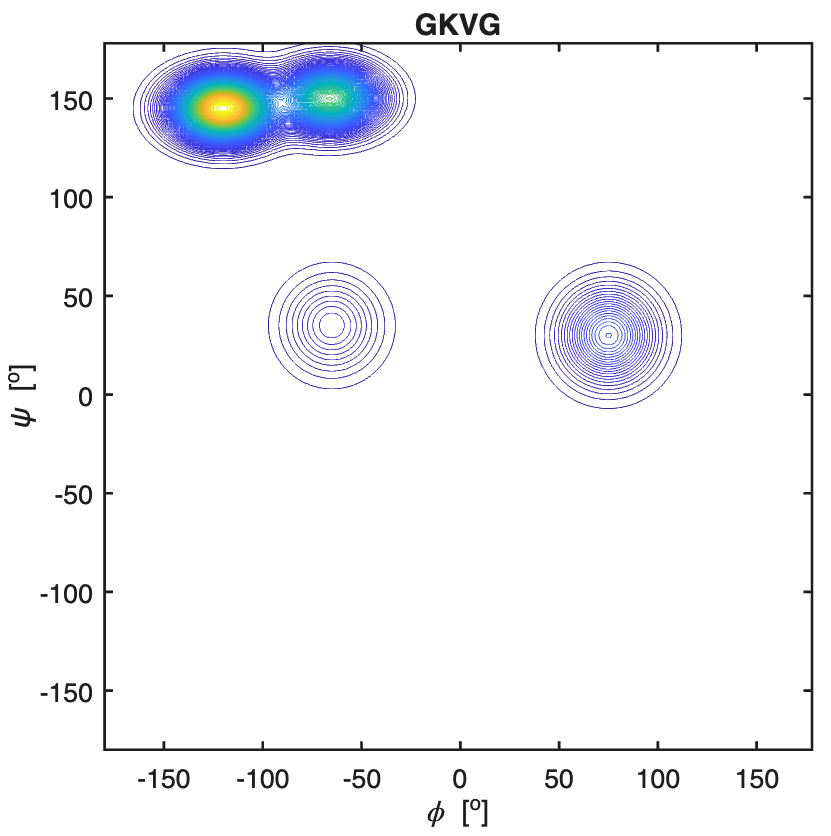

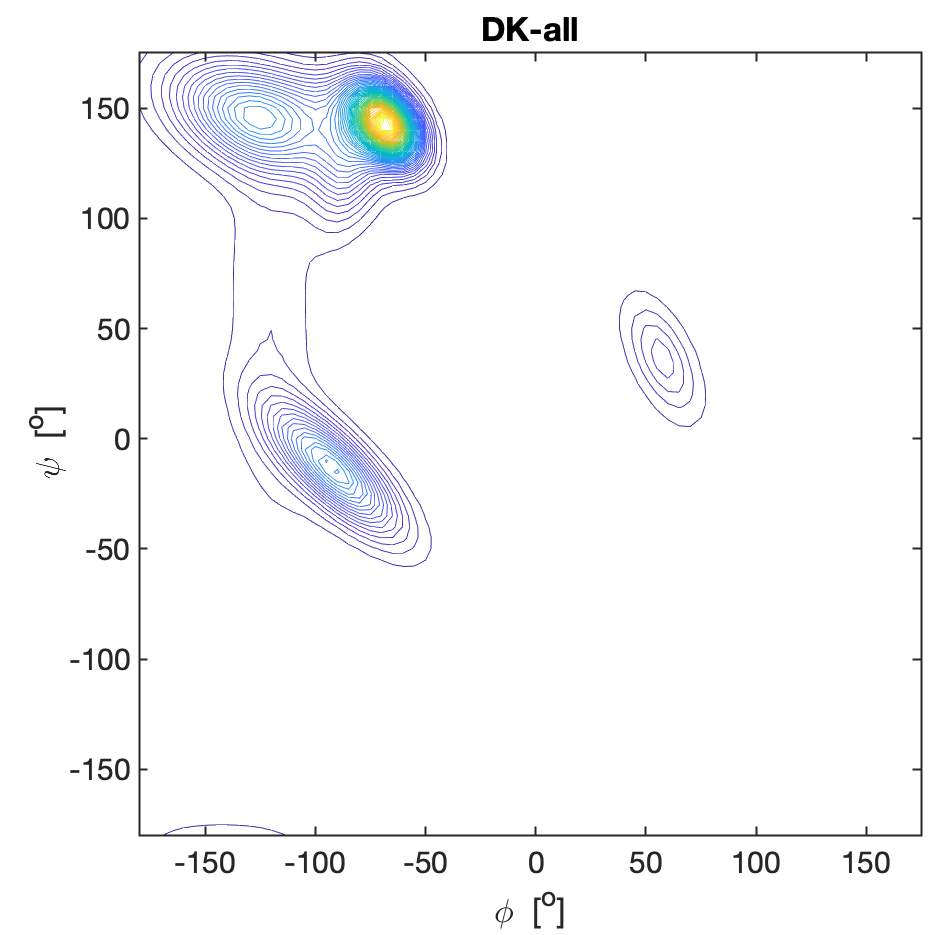

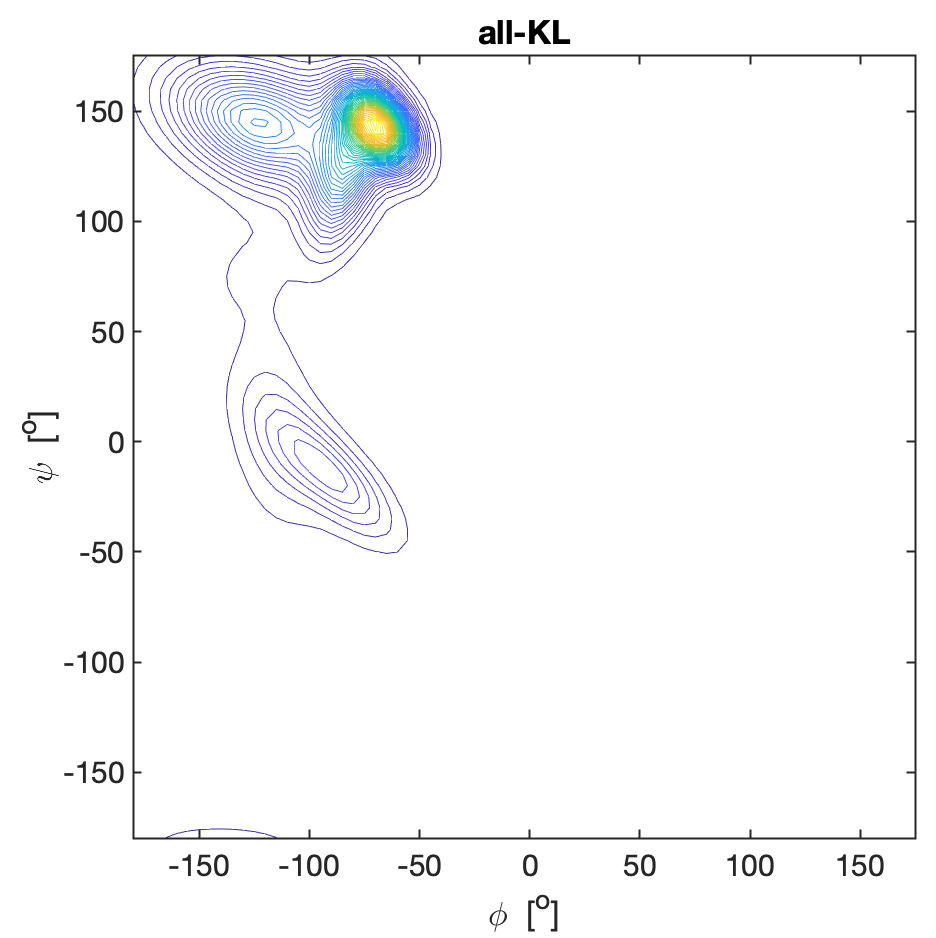

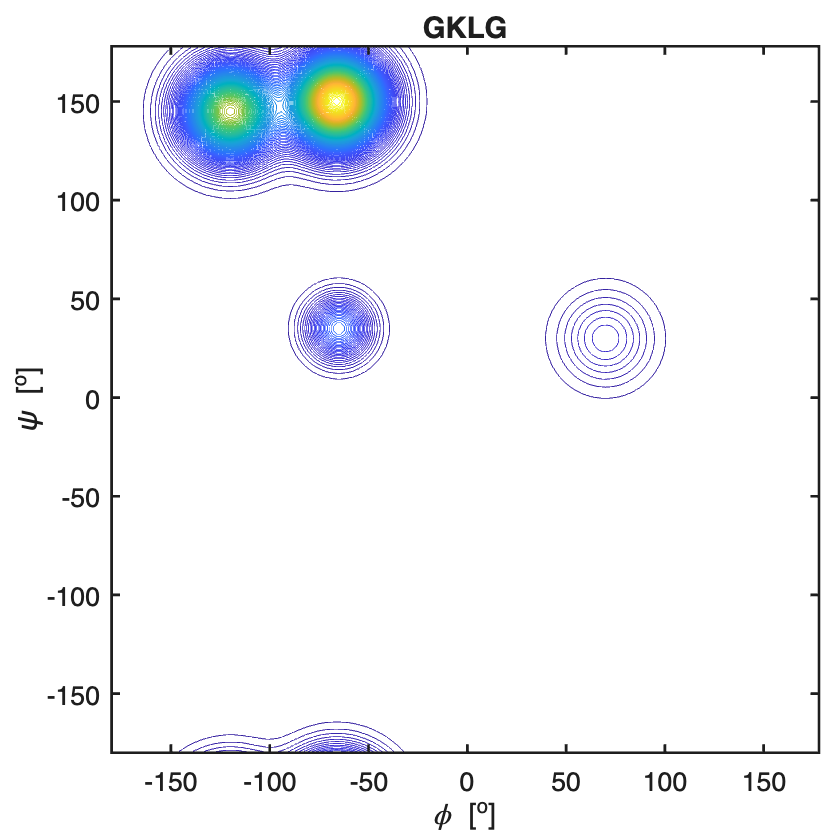

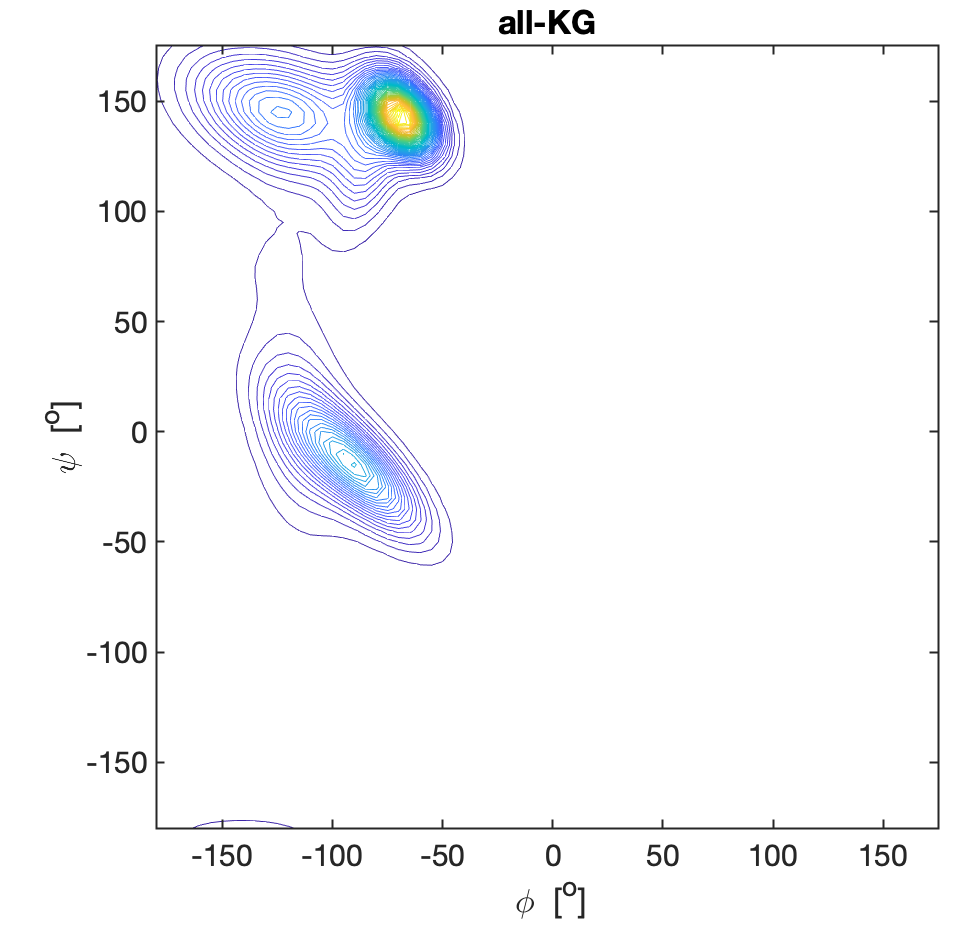

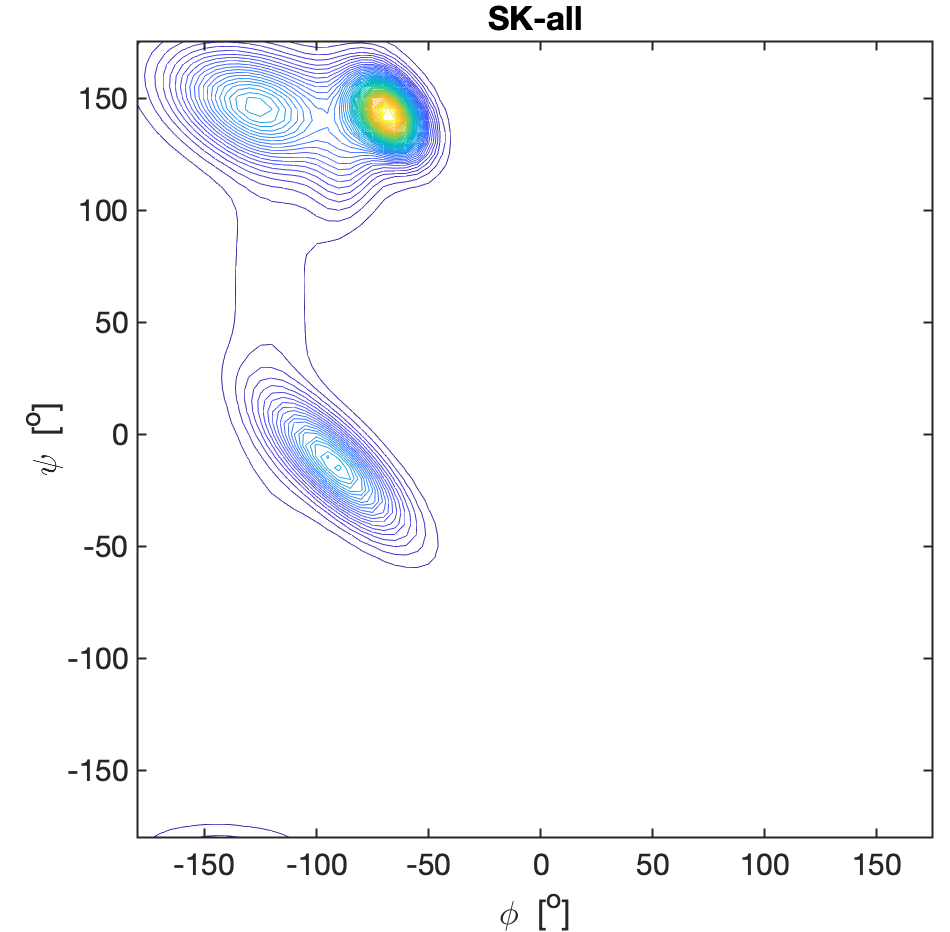

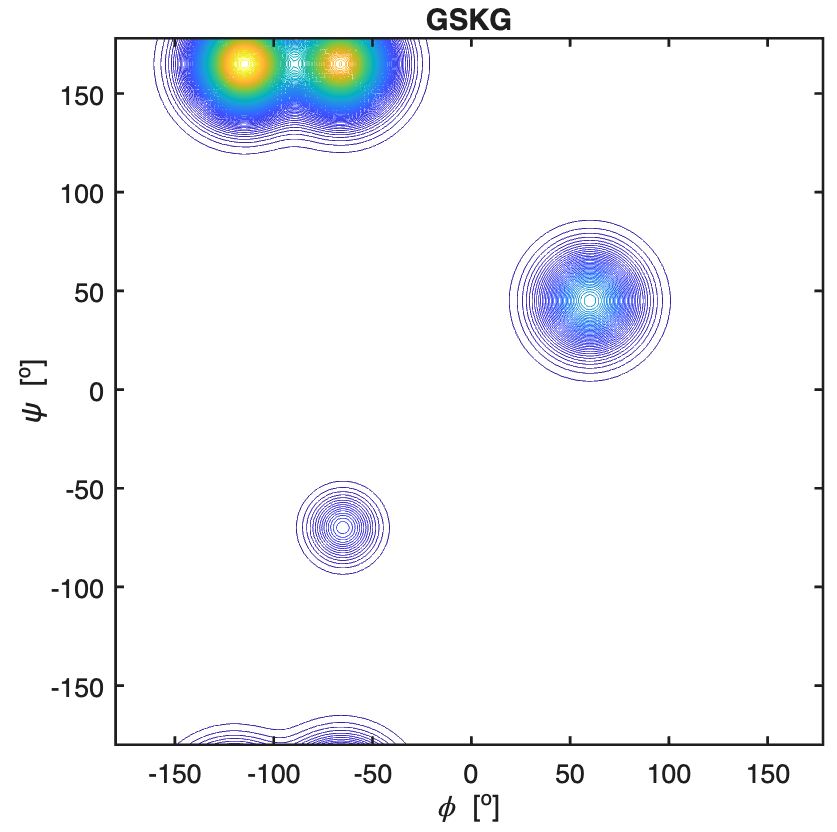

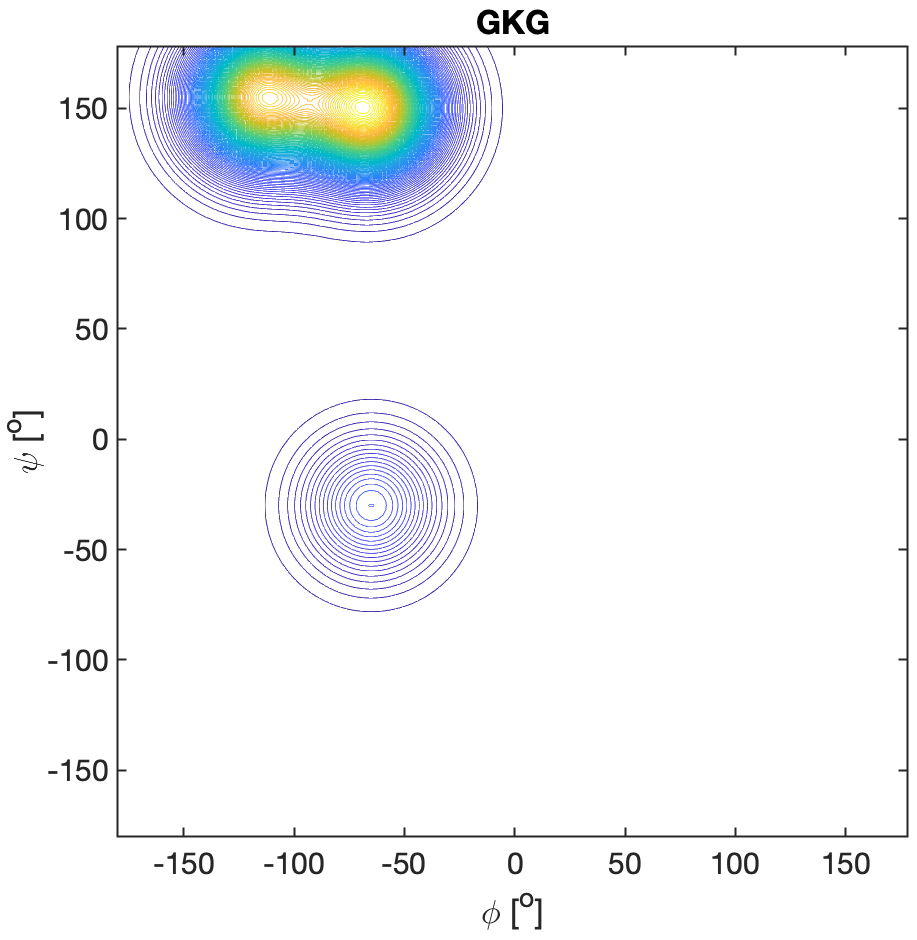
Dunbrack. (13)

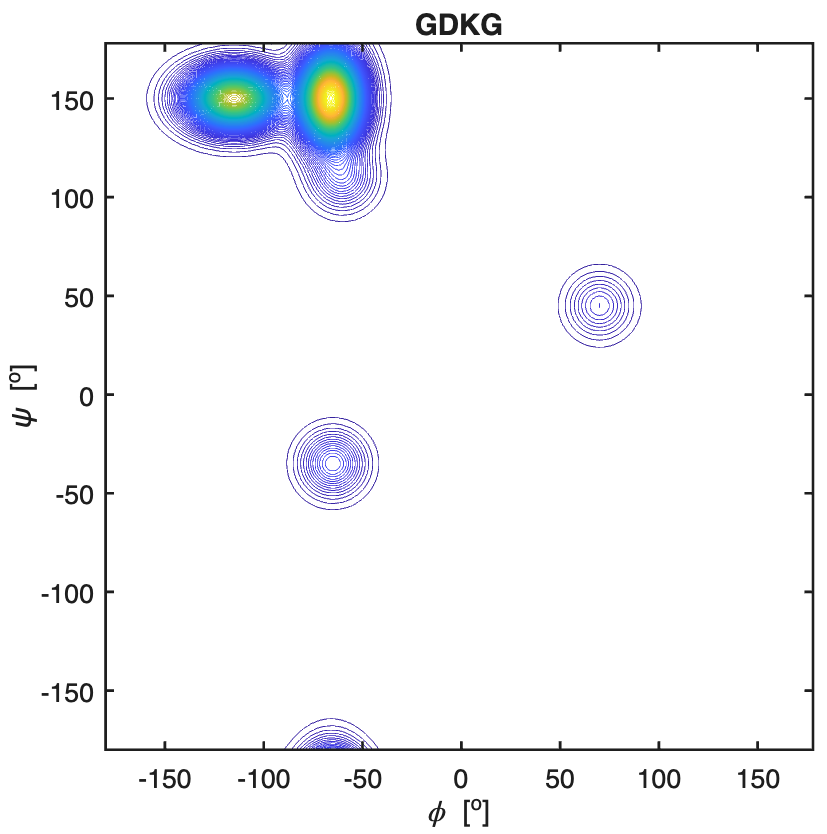

**Figure S14.** Ramachandran plots of valine residues in GVG, all-VG, GD**V**G, D**V**-all, GS**V**G, **SV**-all, G**V**LG, all-**V**L, GK**V**G, K**V**-all, GA**V**G and A**V**-all. All explored peptides were in their cationic state. The plots for the peptides were constructed with a Gaussian model and the parameters listed in ref.(14) for GVG and in Table S2 for the tetra-peptides. The plots for the coil library segments were built with files obtained from the website of the Dunbrack group with the kind permission of Dr. Dunbrack.(13)

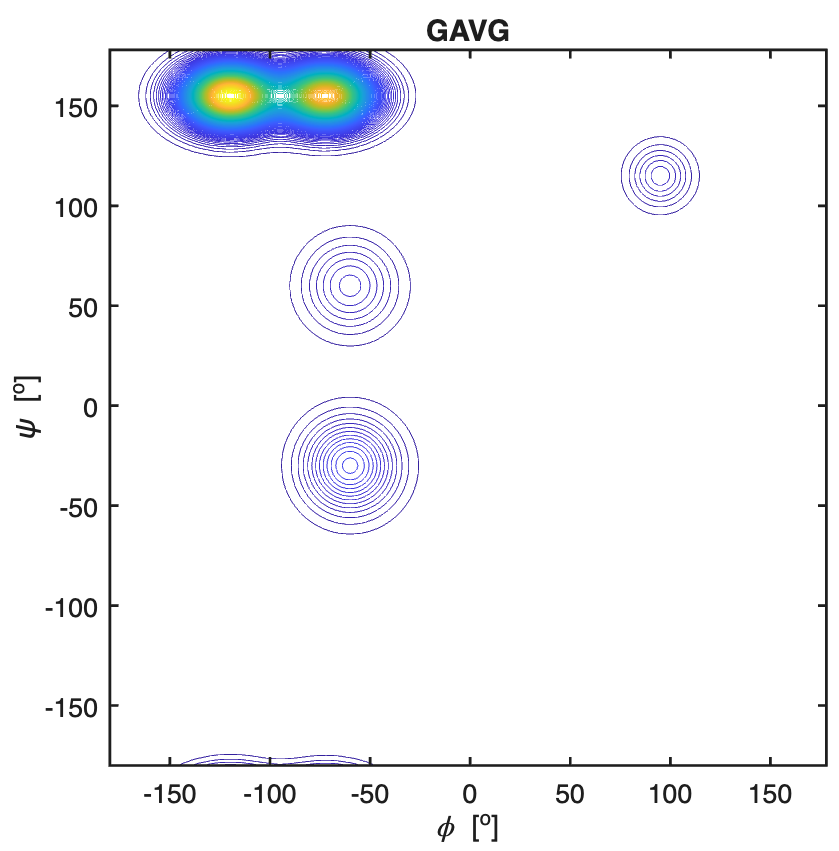

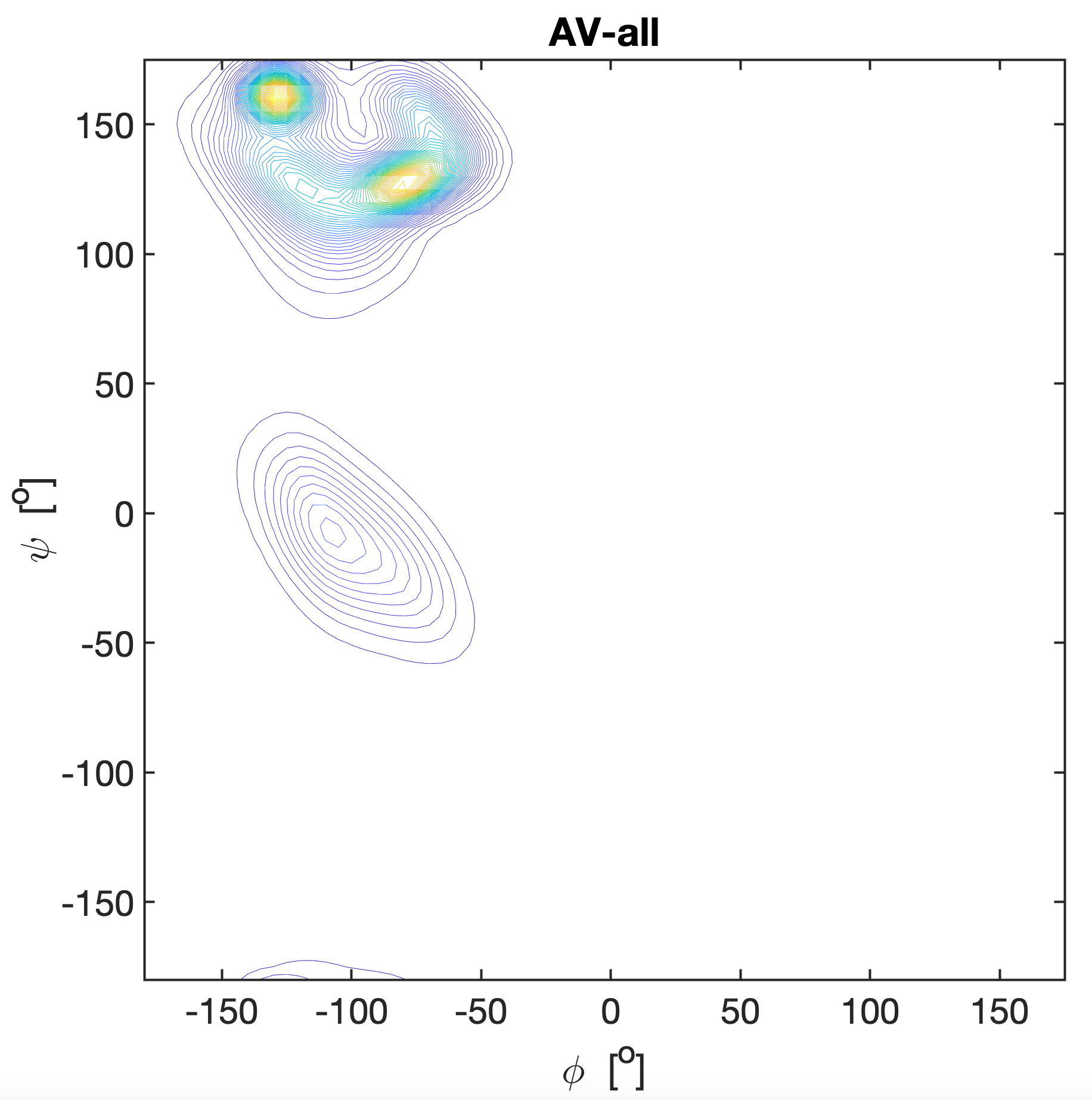

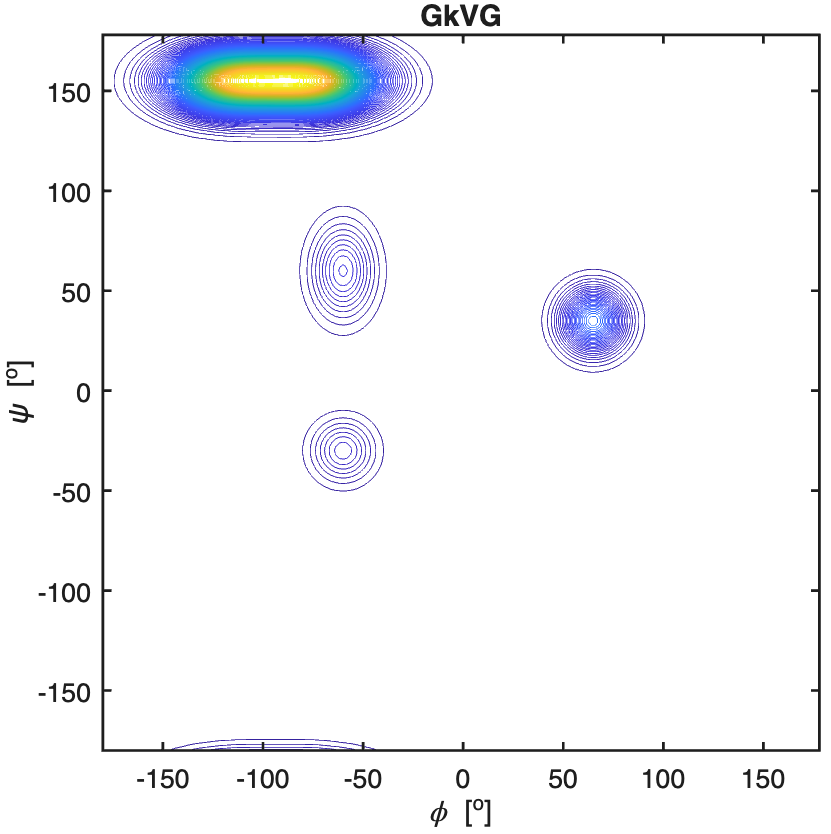

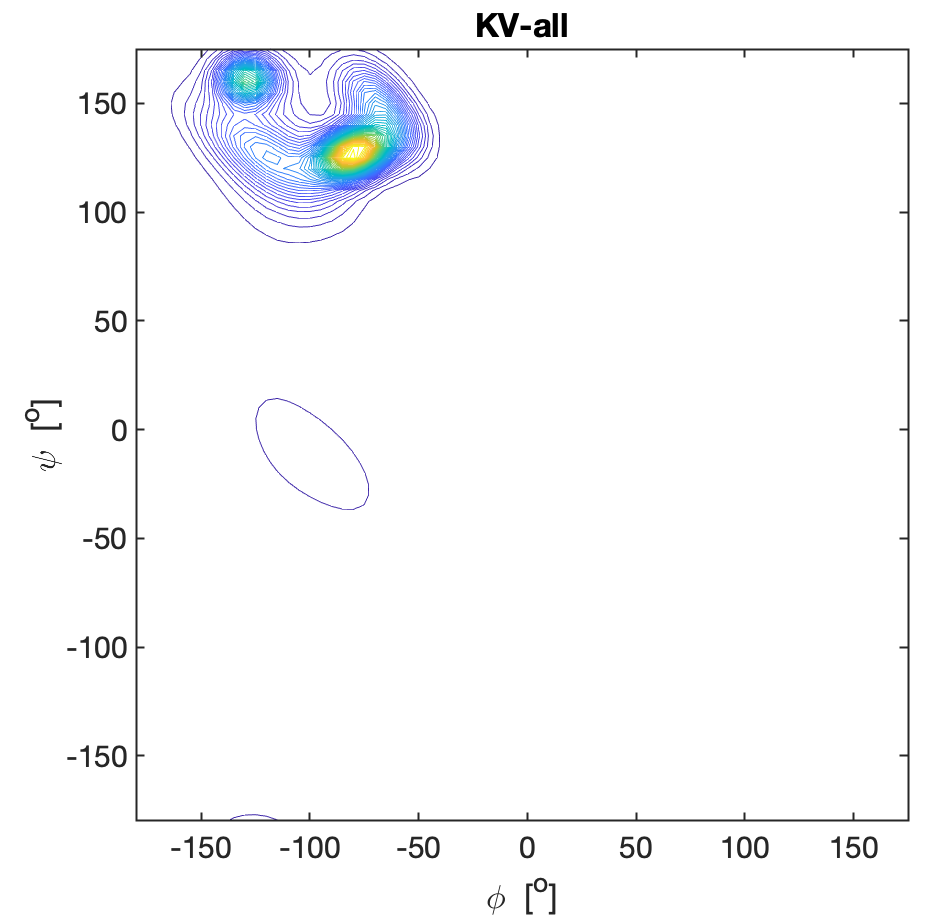

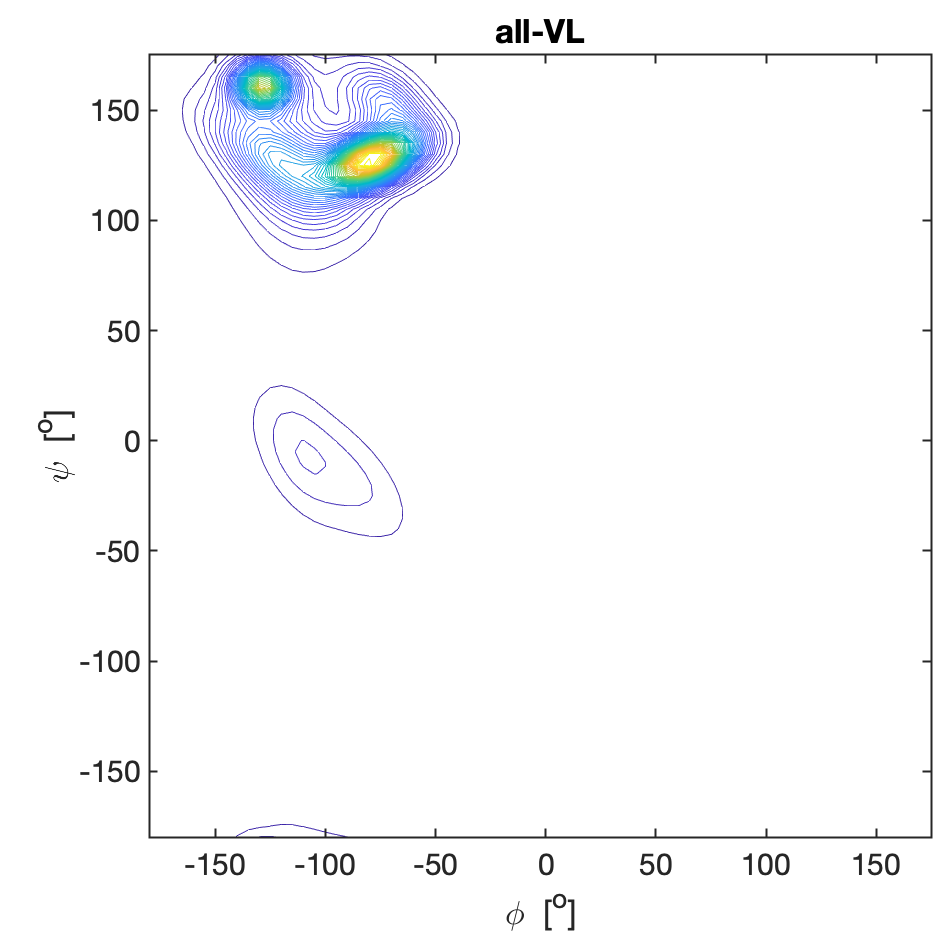

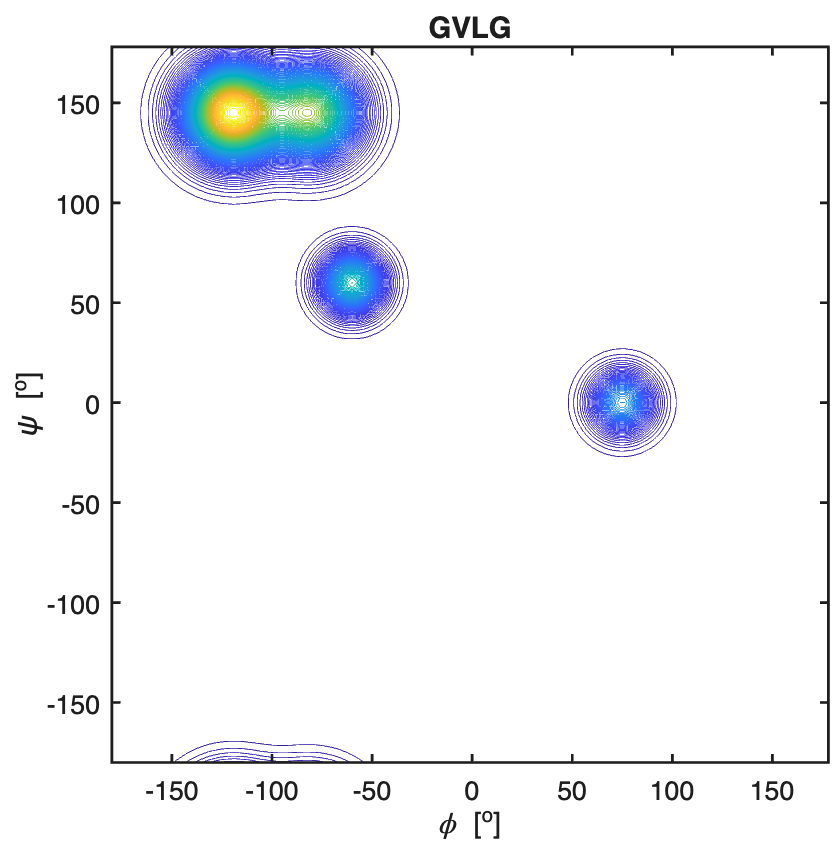

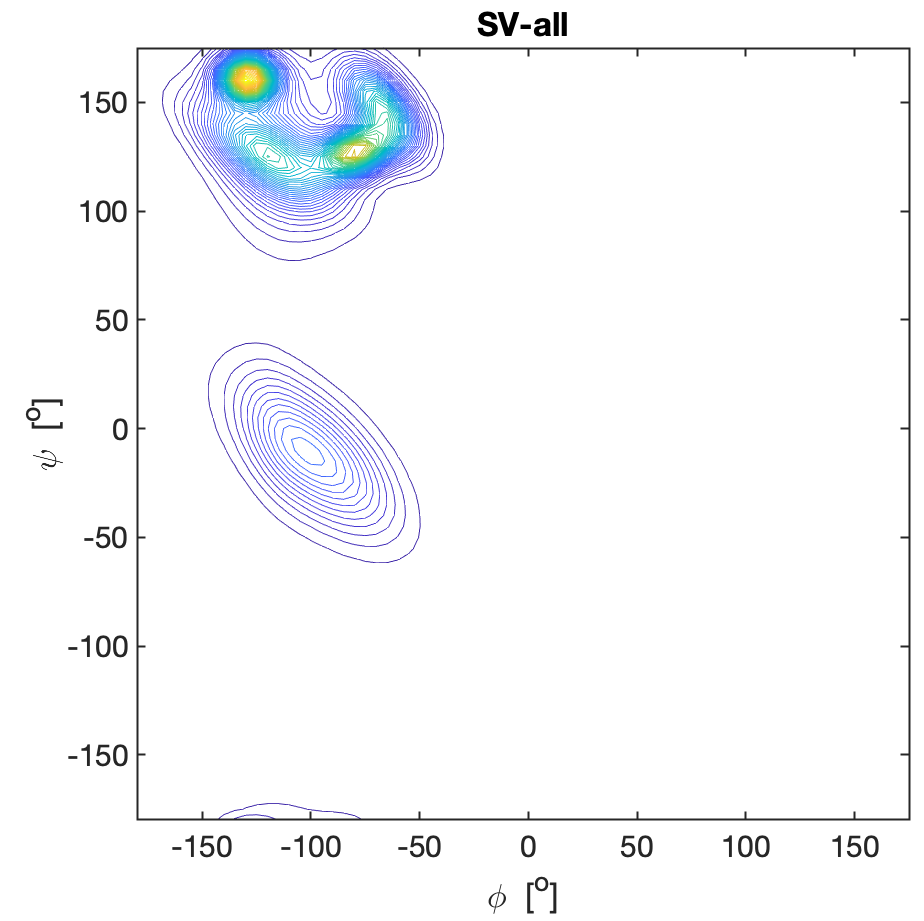

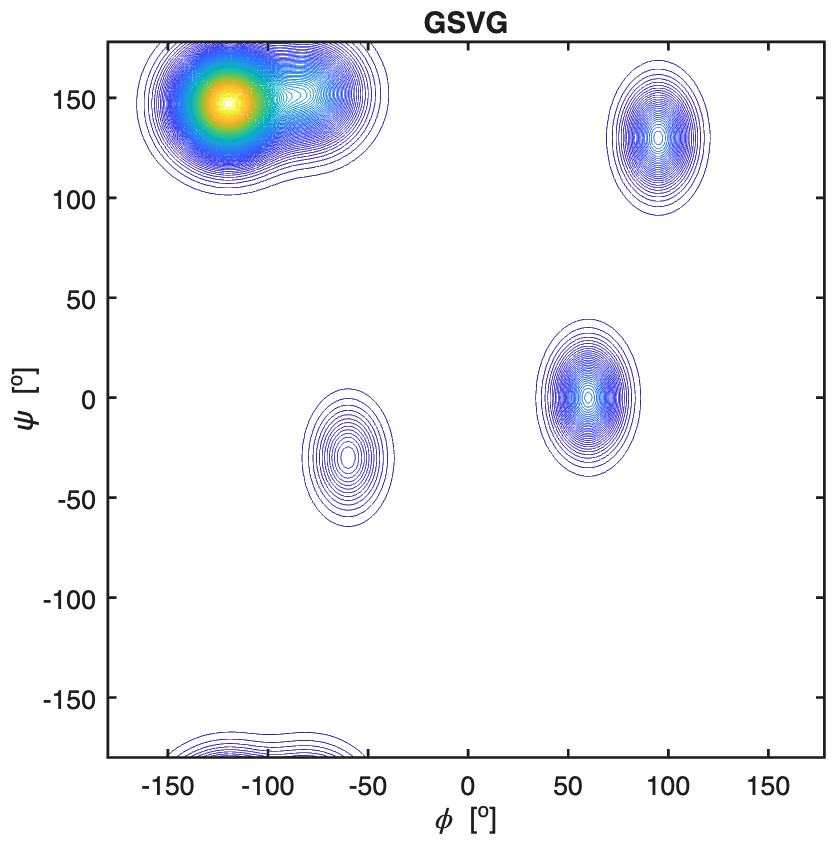

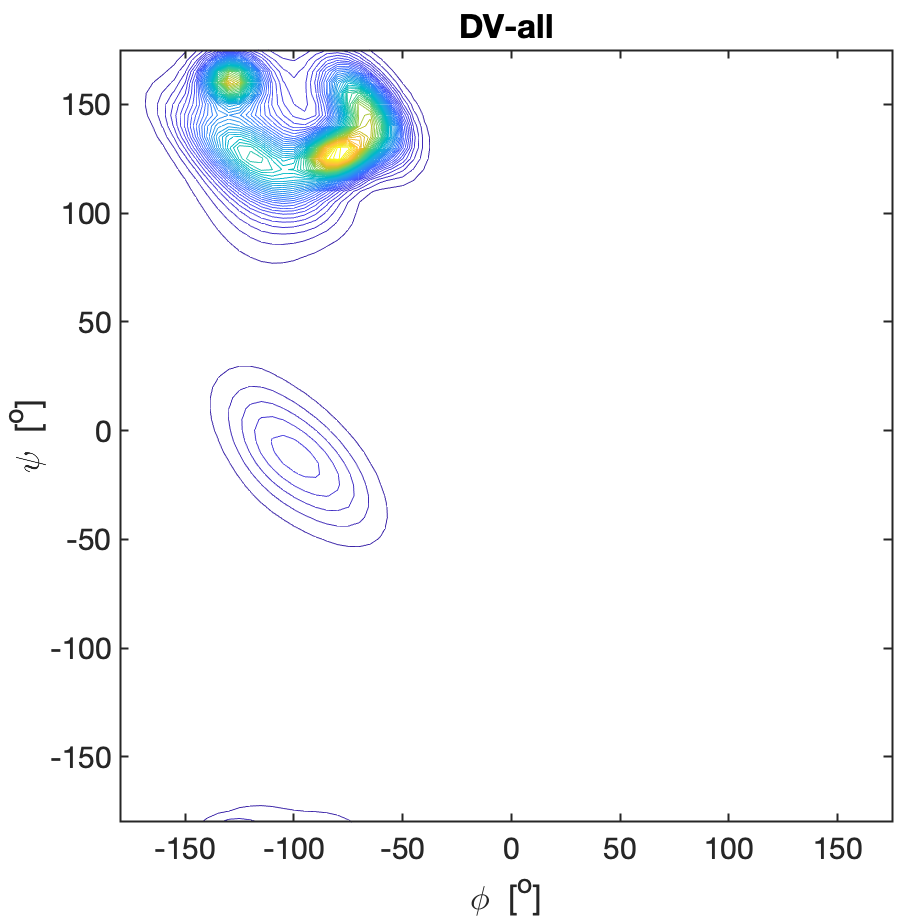

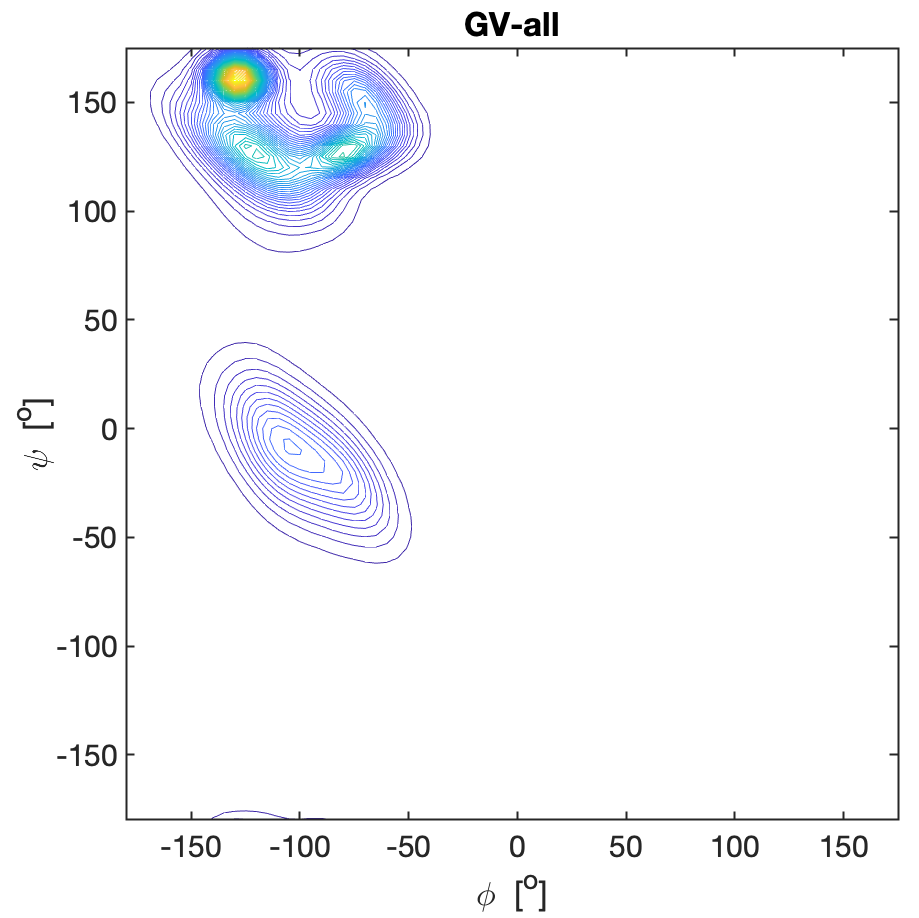

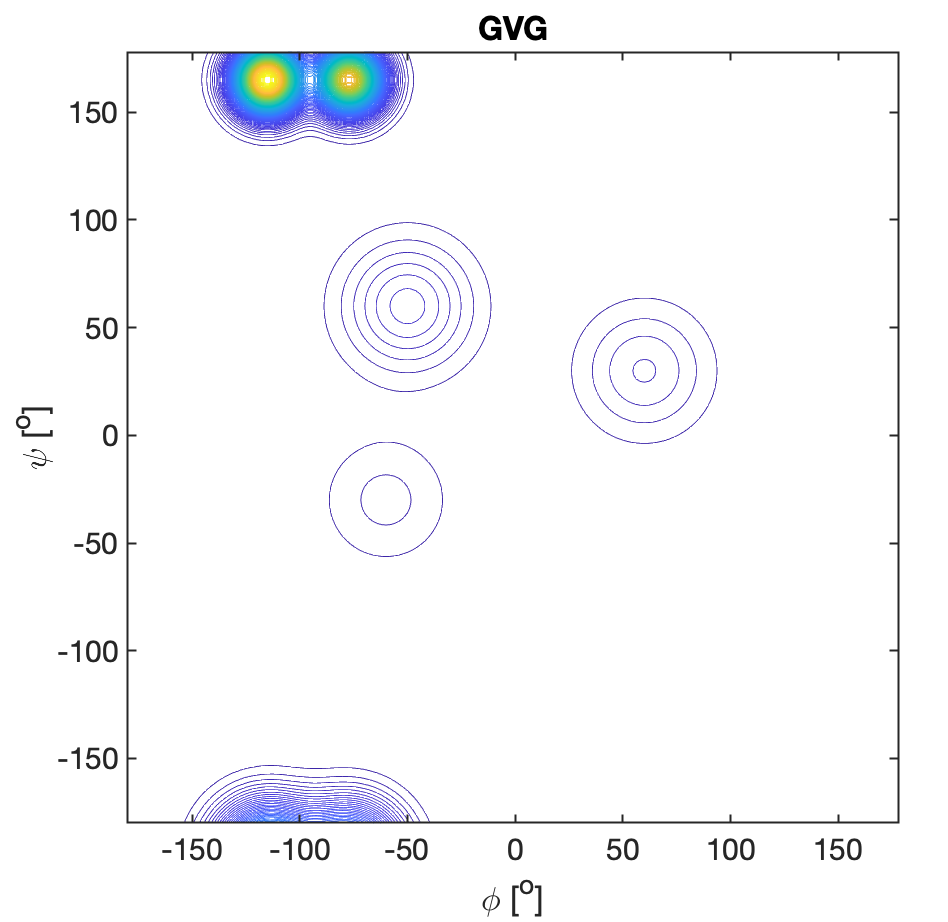

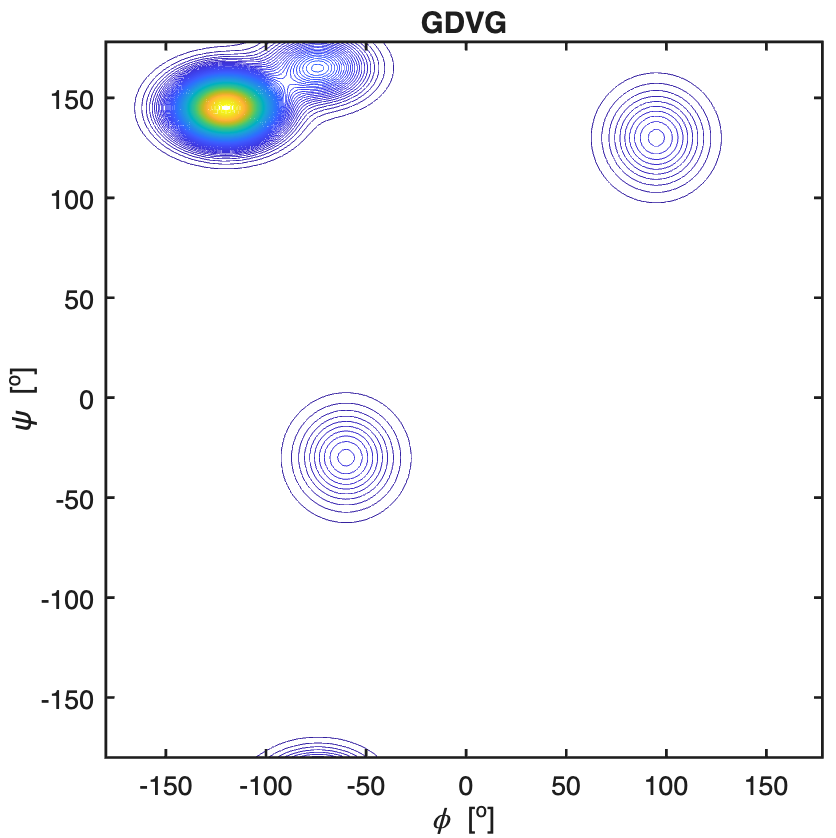

**Figure S15.** Ramachandran plots of phenylalanine residues in GFG, all-**F**G, G**F**AG, FA-all, G**F**DG, all-**F**D, GD**F**G, and D**F**-all. All explored peptides were in their cationic state. The plots for the peptides were constructed with a Gaussian model and the parameters listed in ref.(14) for GFG and in ref.(6) for the tetra-peptides. The plots for the coil library segments were built with files
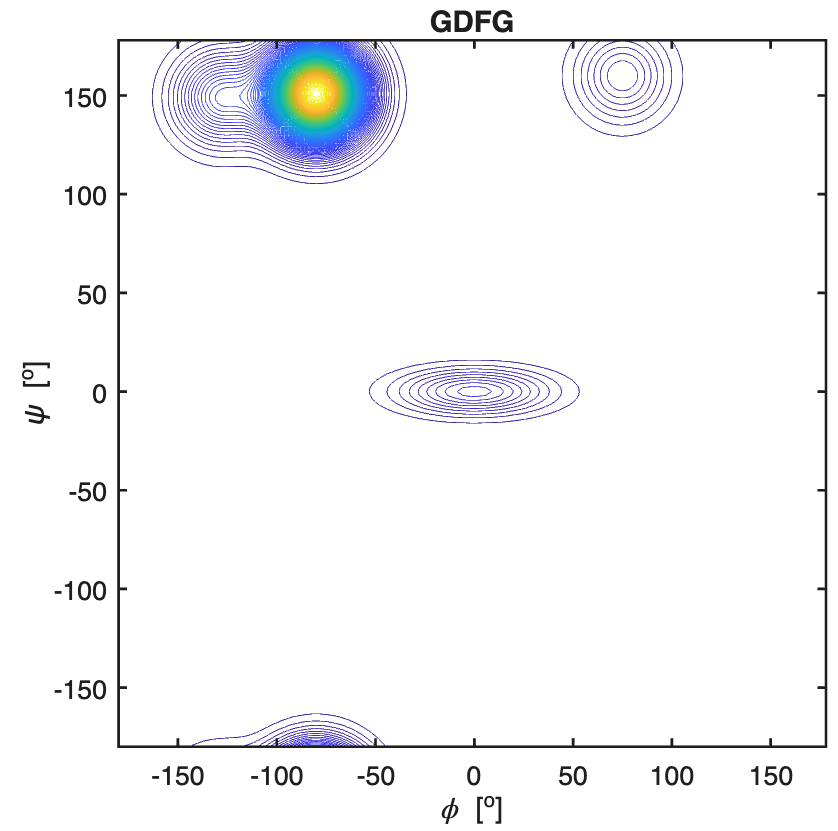

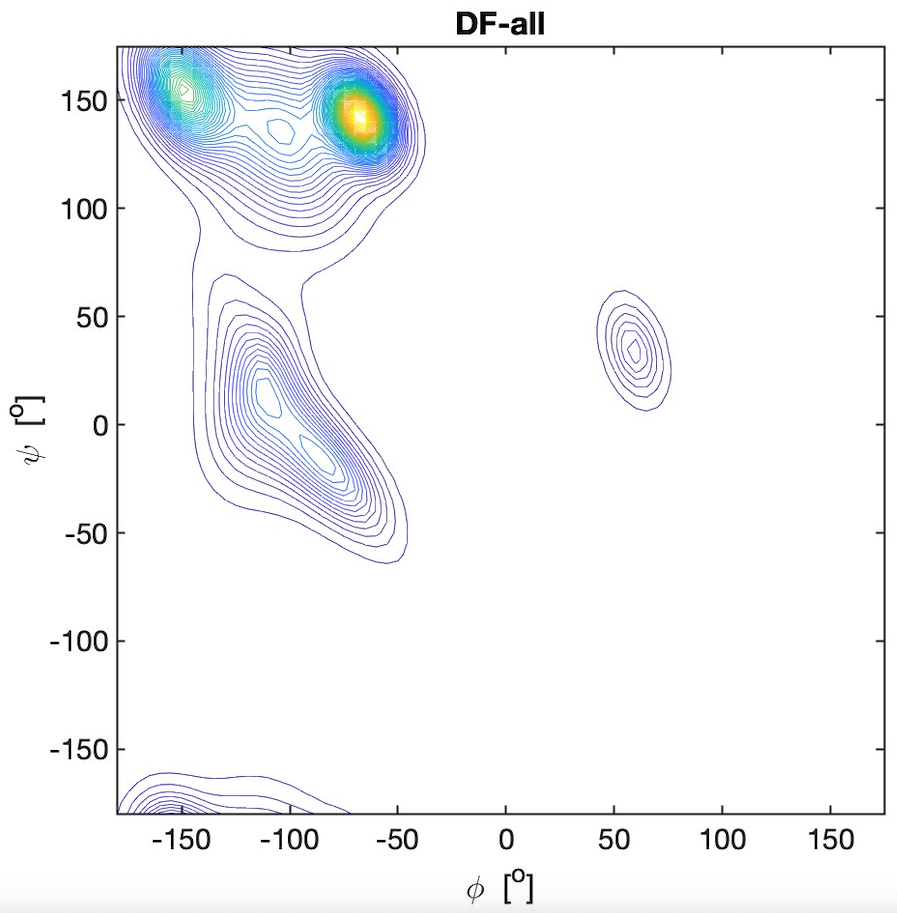
obtained
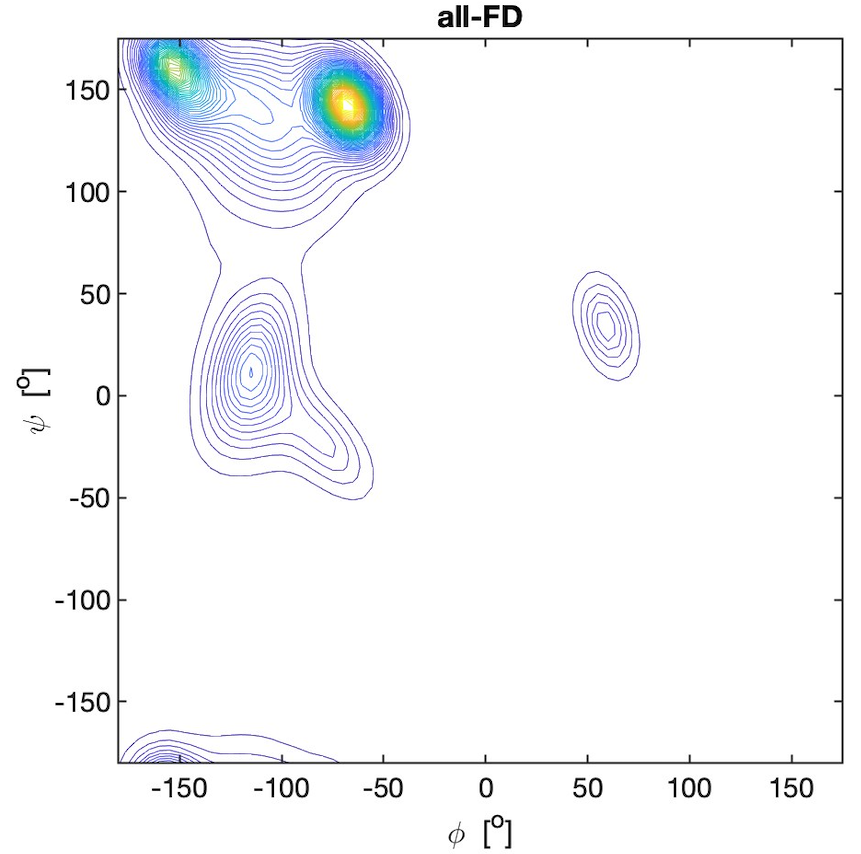

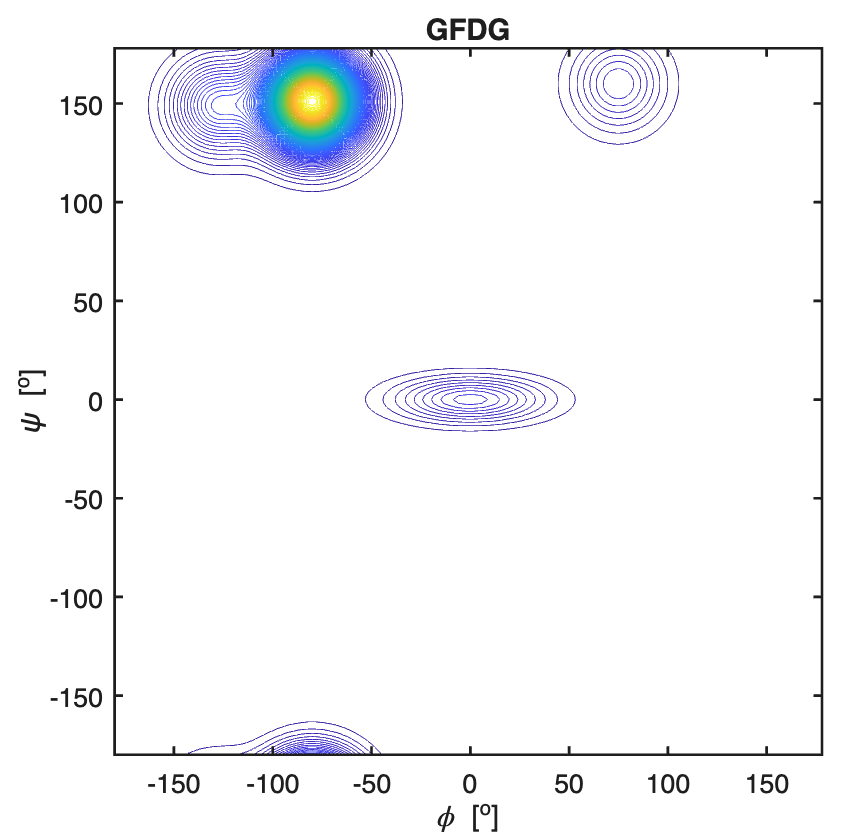

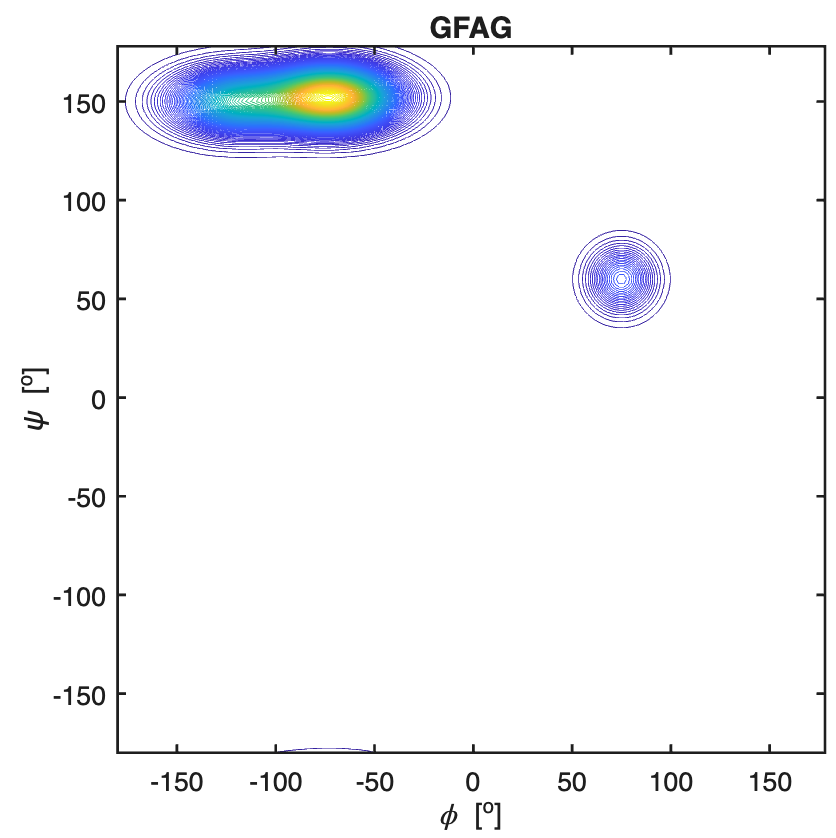

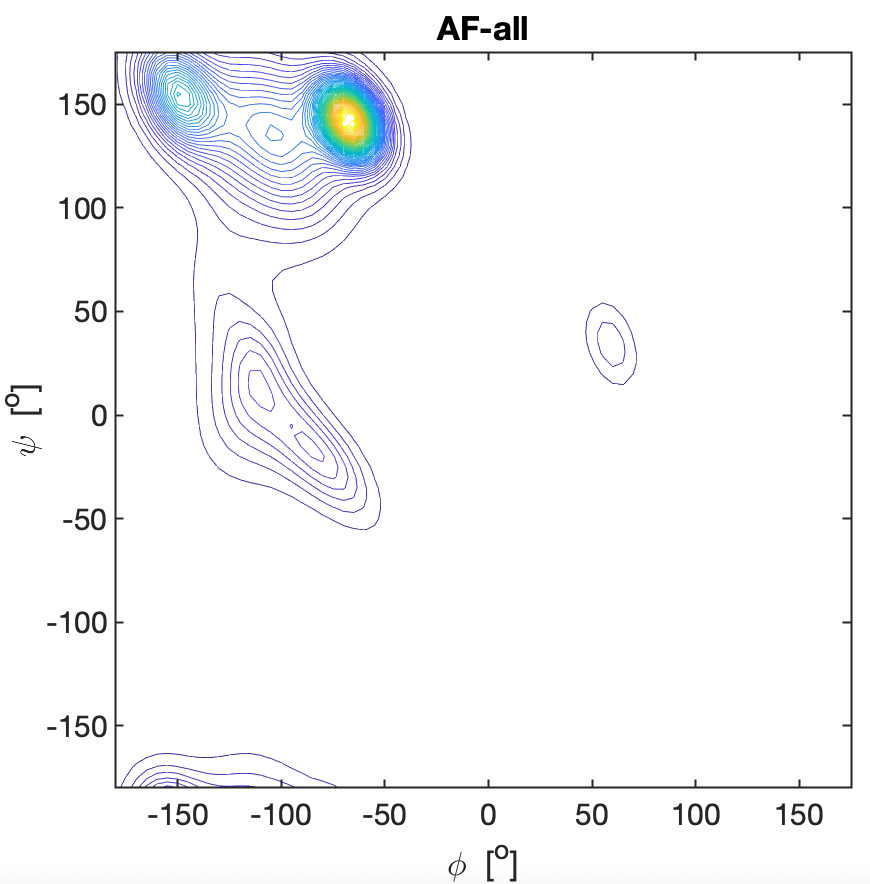

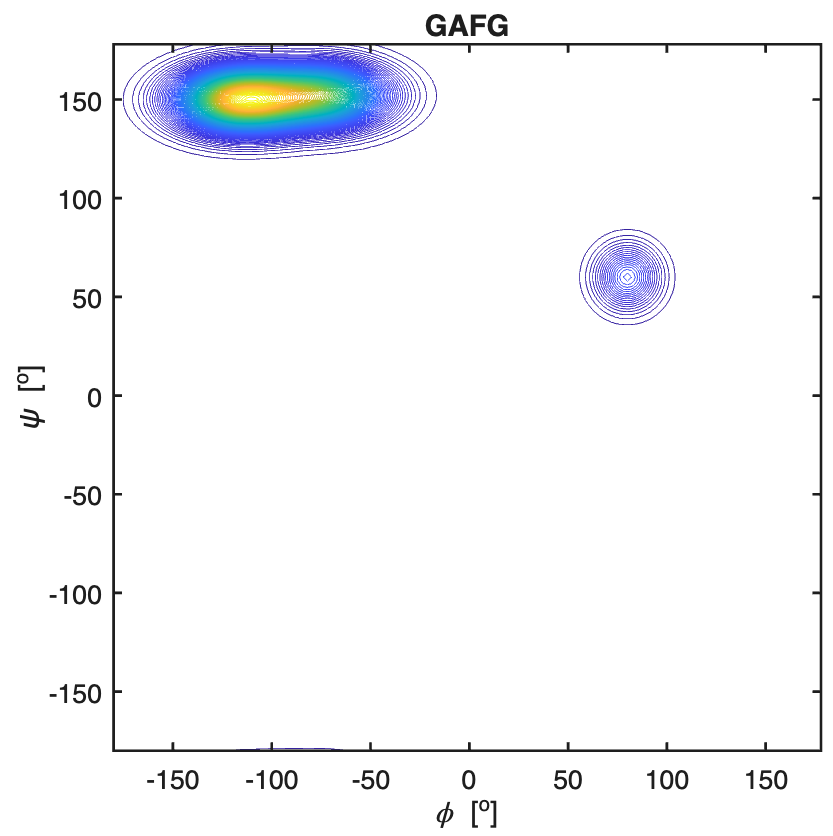

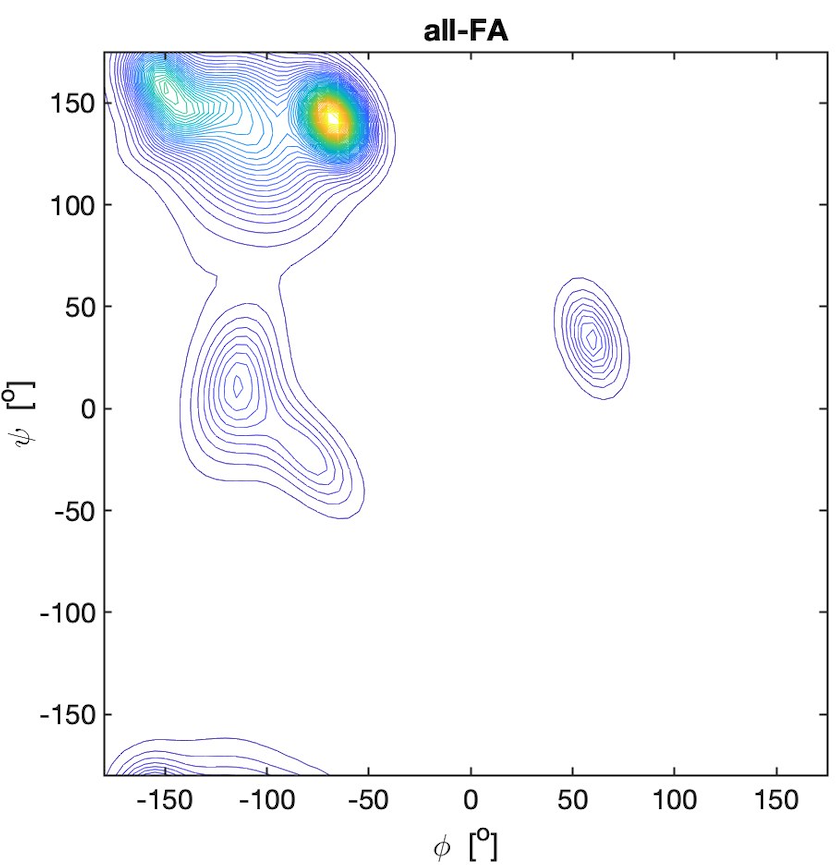

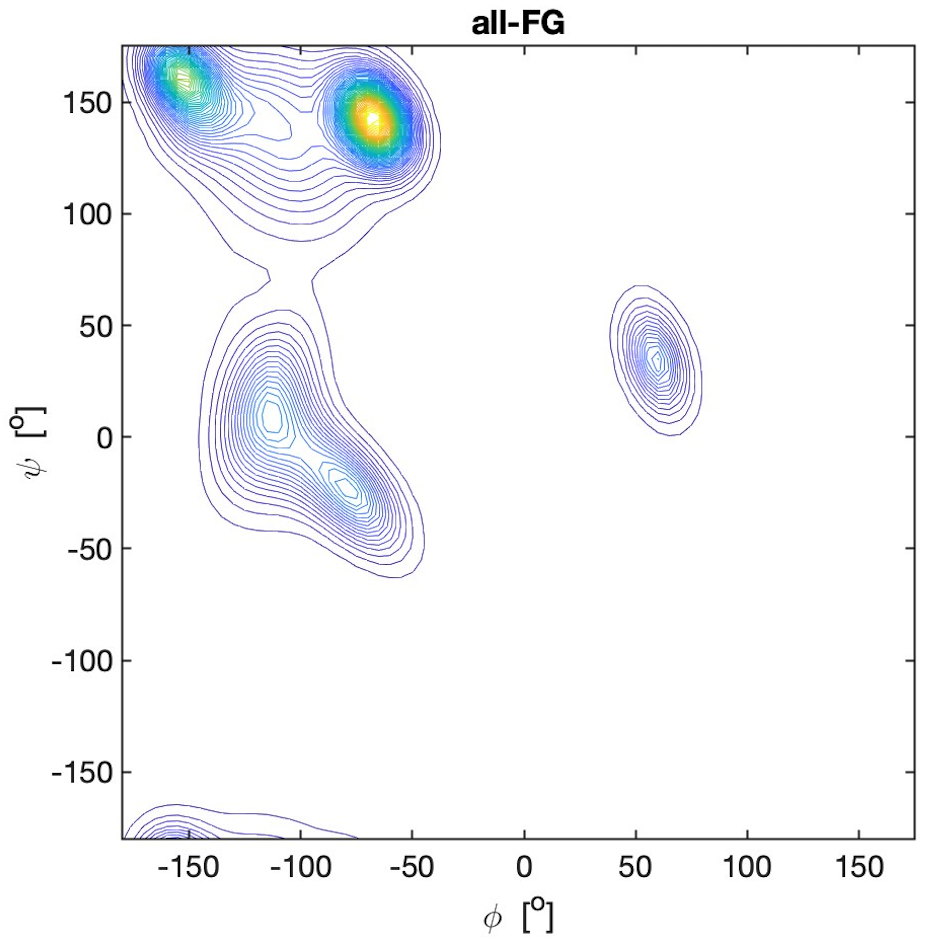
from the website of the Dunbrack group with the kind permission
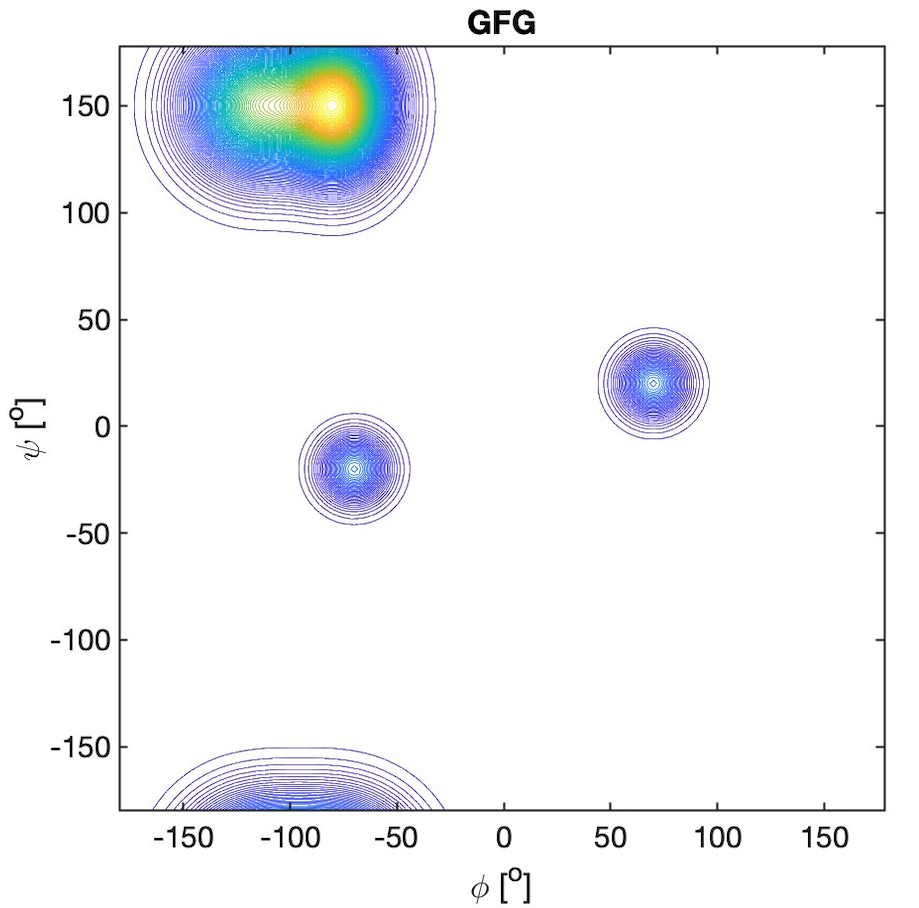
of Dr. Dunbrack.(13)

**Figure S16.** Ramachandran plots of phenylalanine residues in GRG, all-**R**G, G**R**RG, all-**R**R, GR**R**G, and R**R**-all. All explored peptides were in their fully protonated state. The plots for the peptides were constructed with a Gaussian model and the parameters listed in ref.(14) for GRG and in ref.(6) for the tetra-peptides. The plots for the coil library segments were built with files obtained from the website of the Dunbrack group with the kind permission of Dr. Dunbrack.(13)

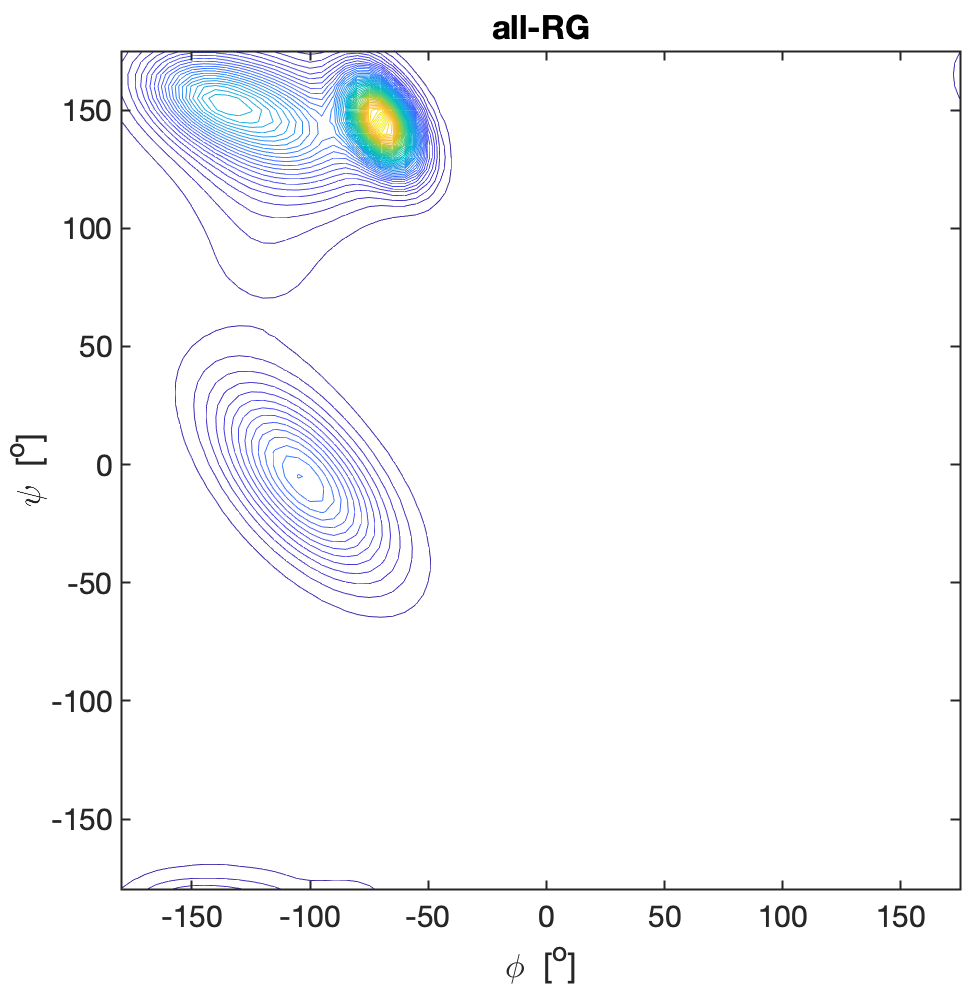

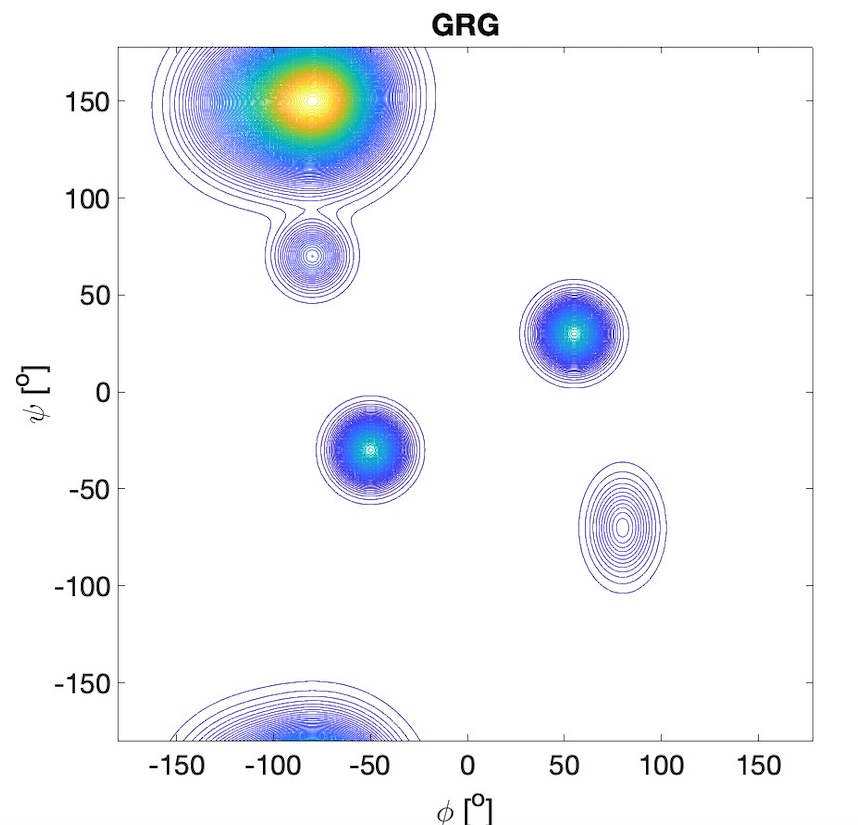

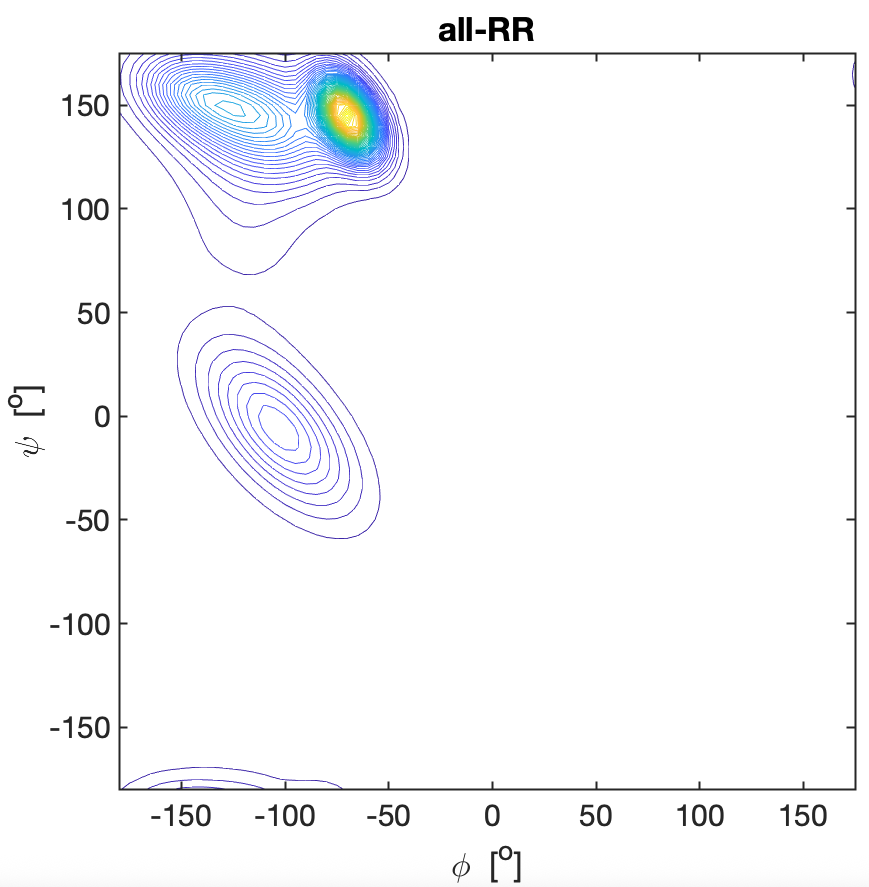

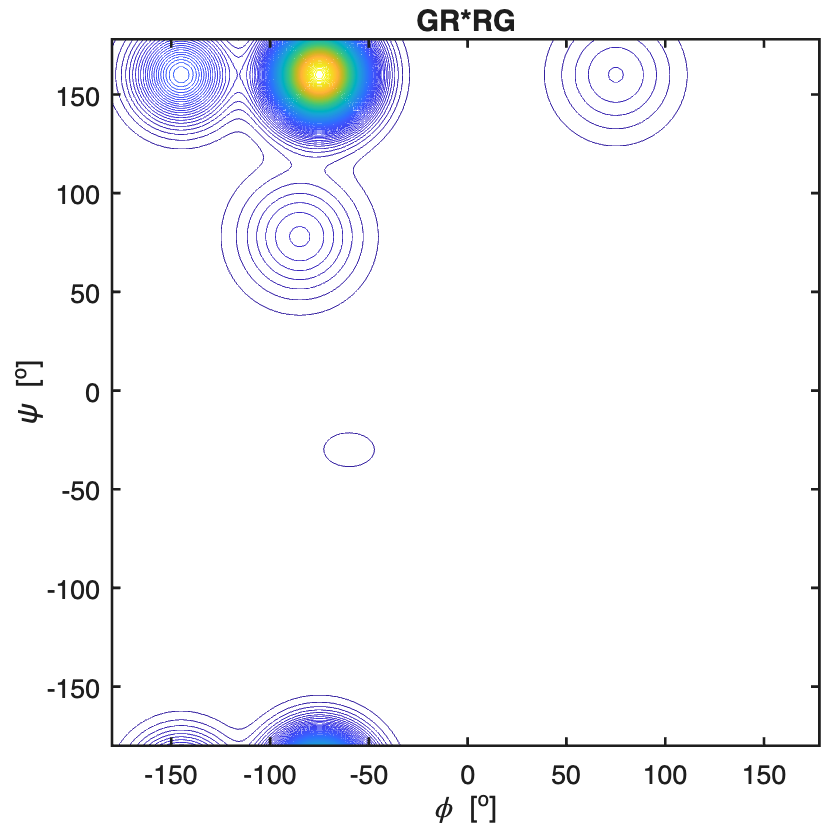

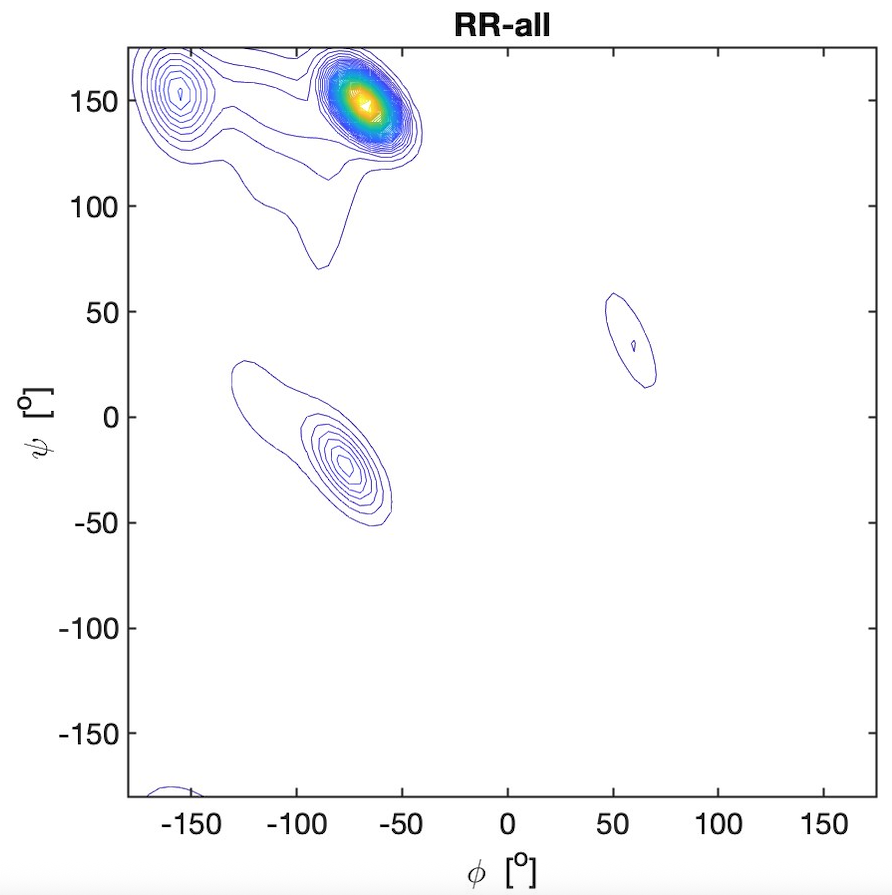

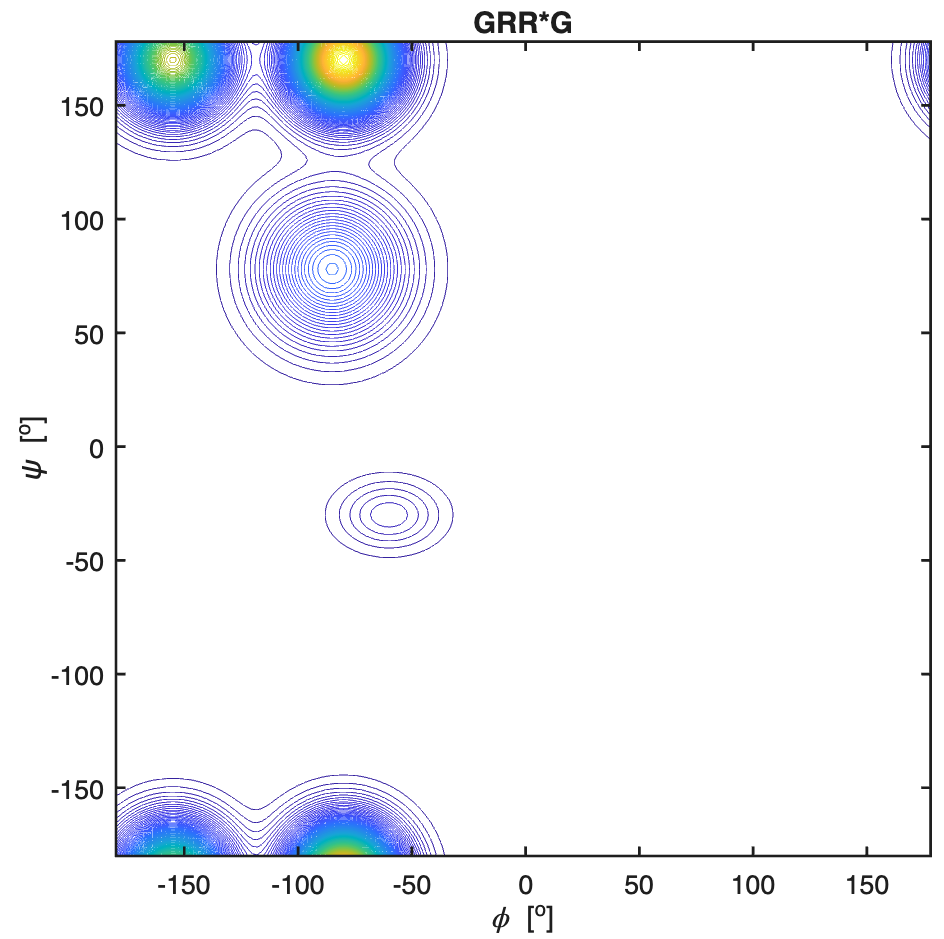

**Figure S17.** Mesostate population of the indicated amino acid residues in GXG, GXYG and in
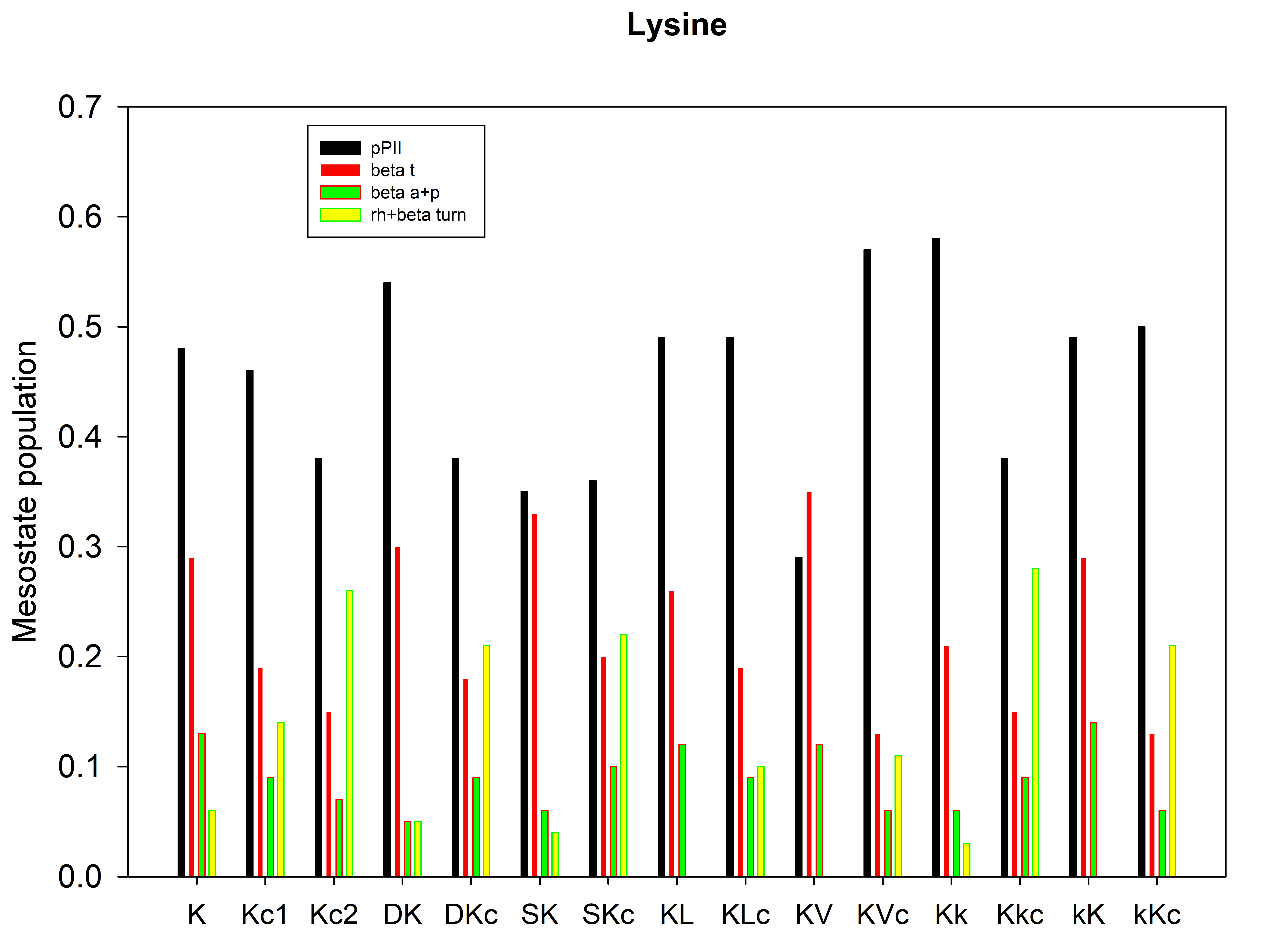

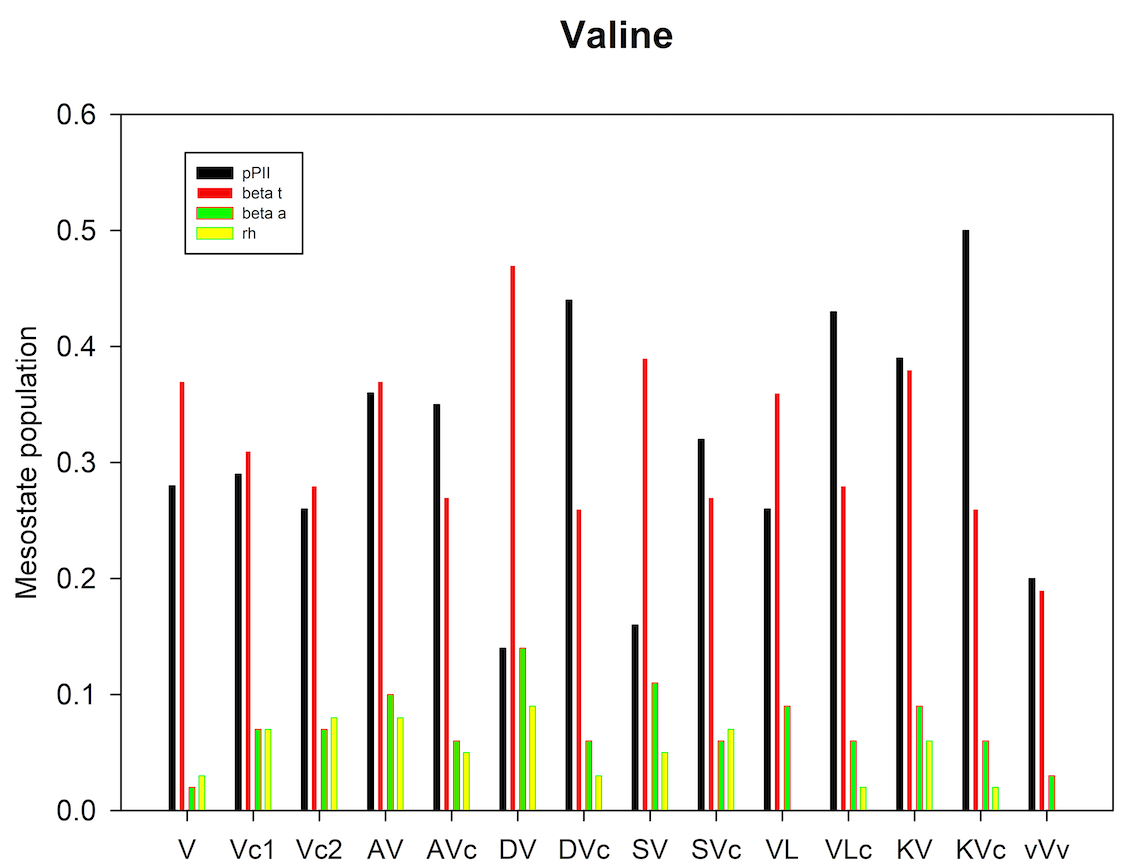

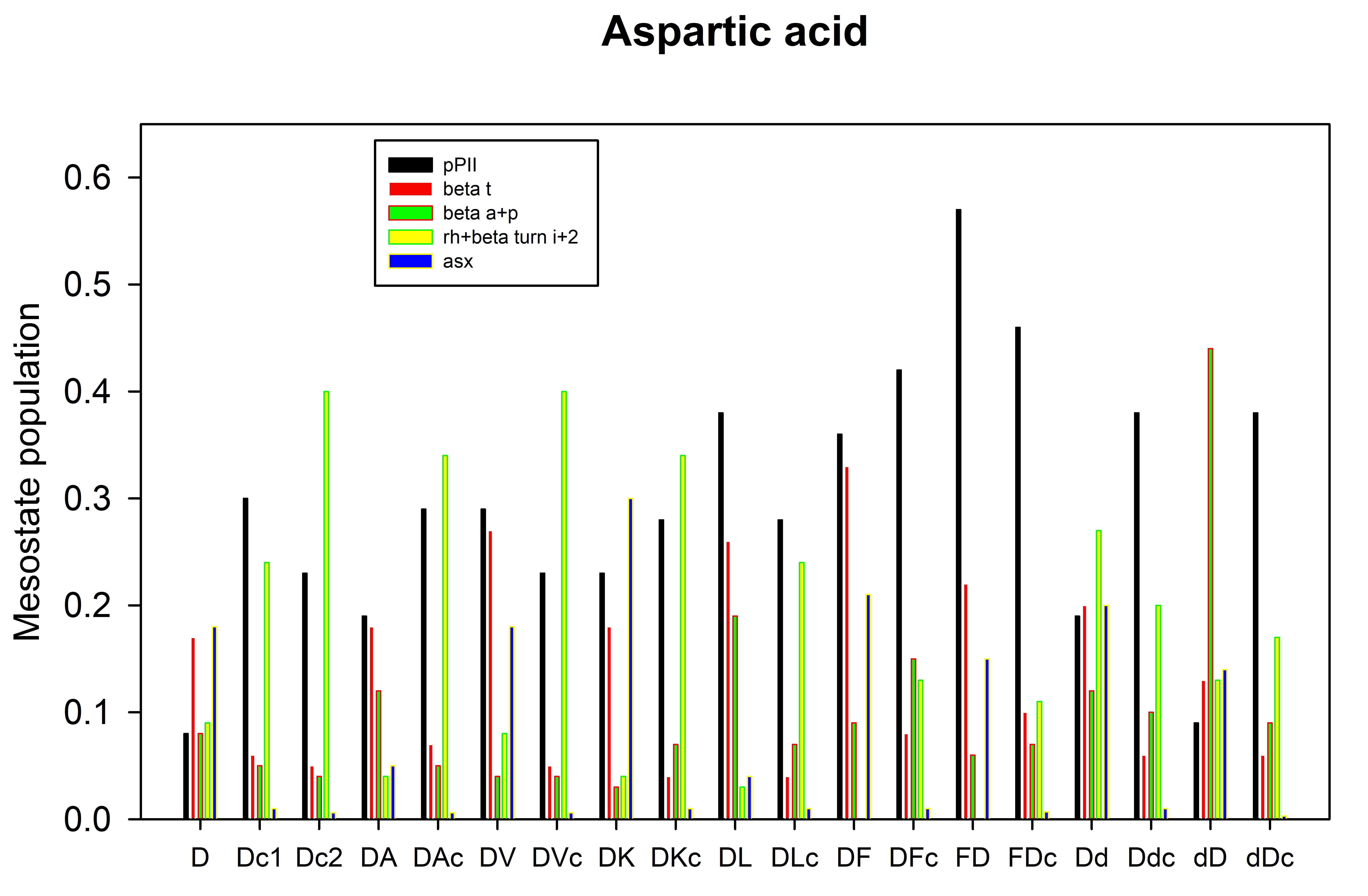
corresponding segments of the coil library of Ting et al.(13)
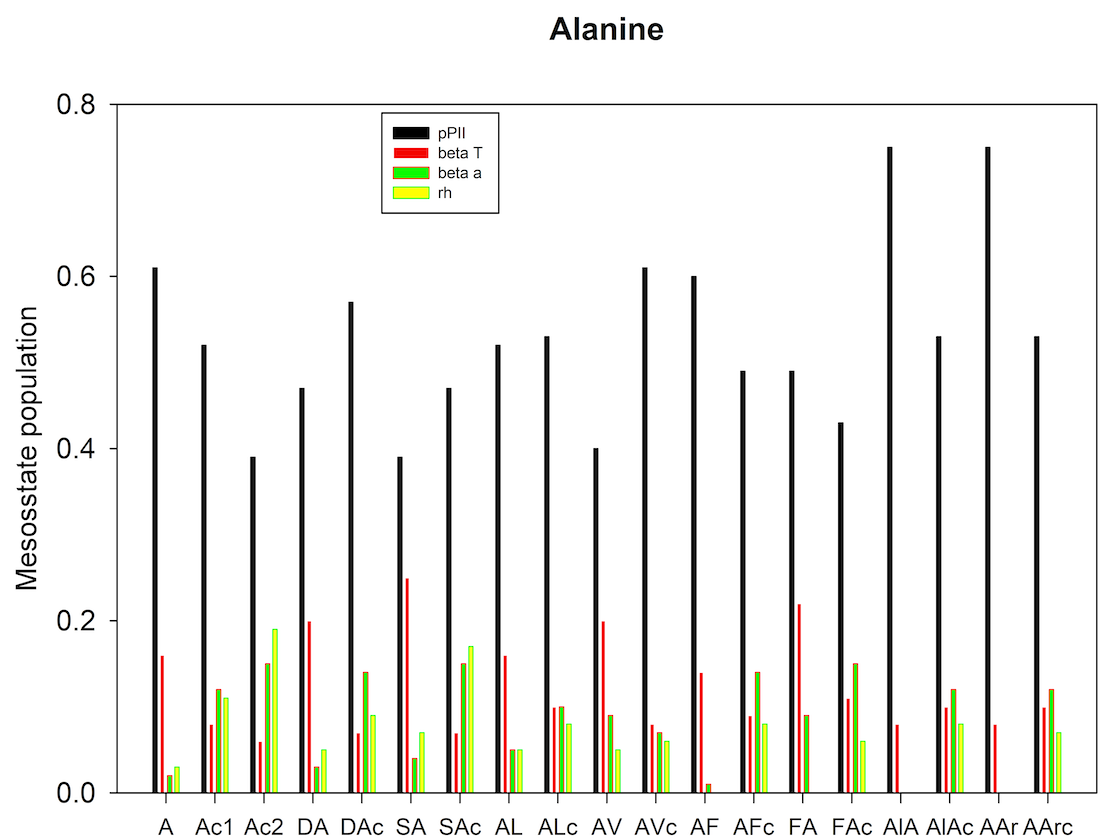

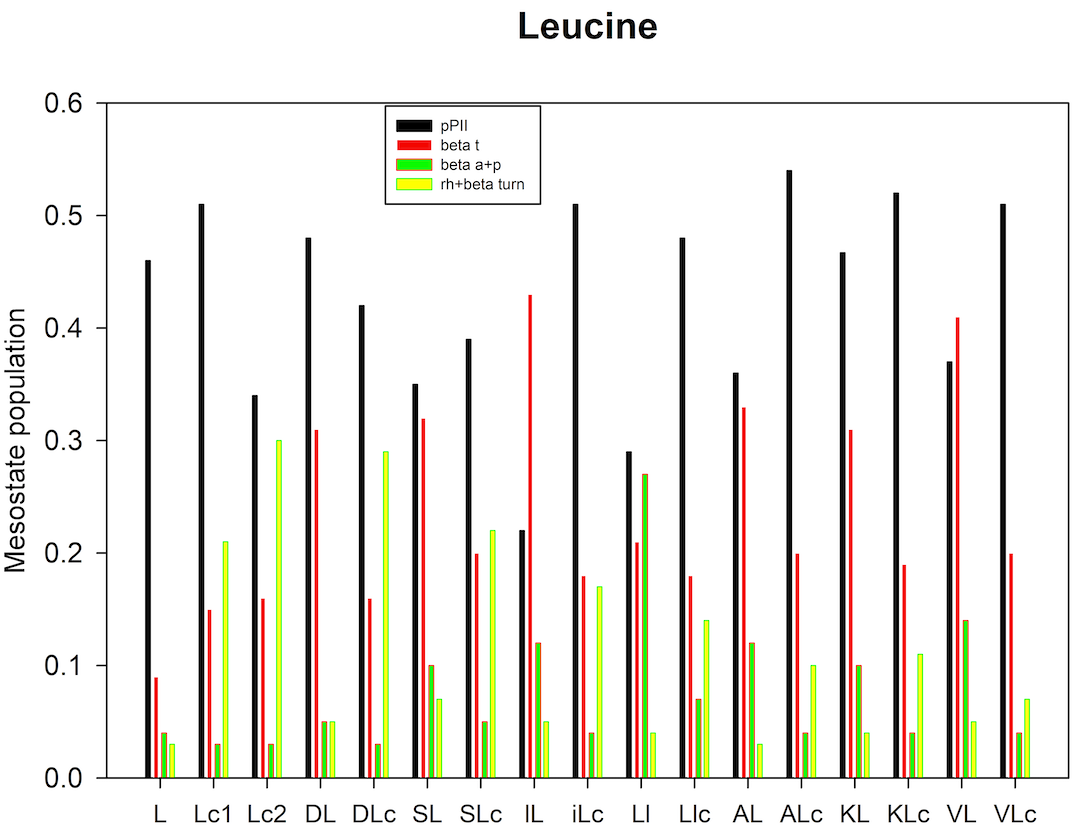

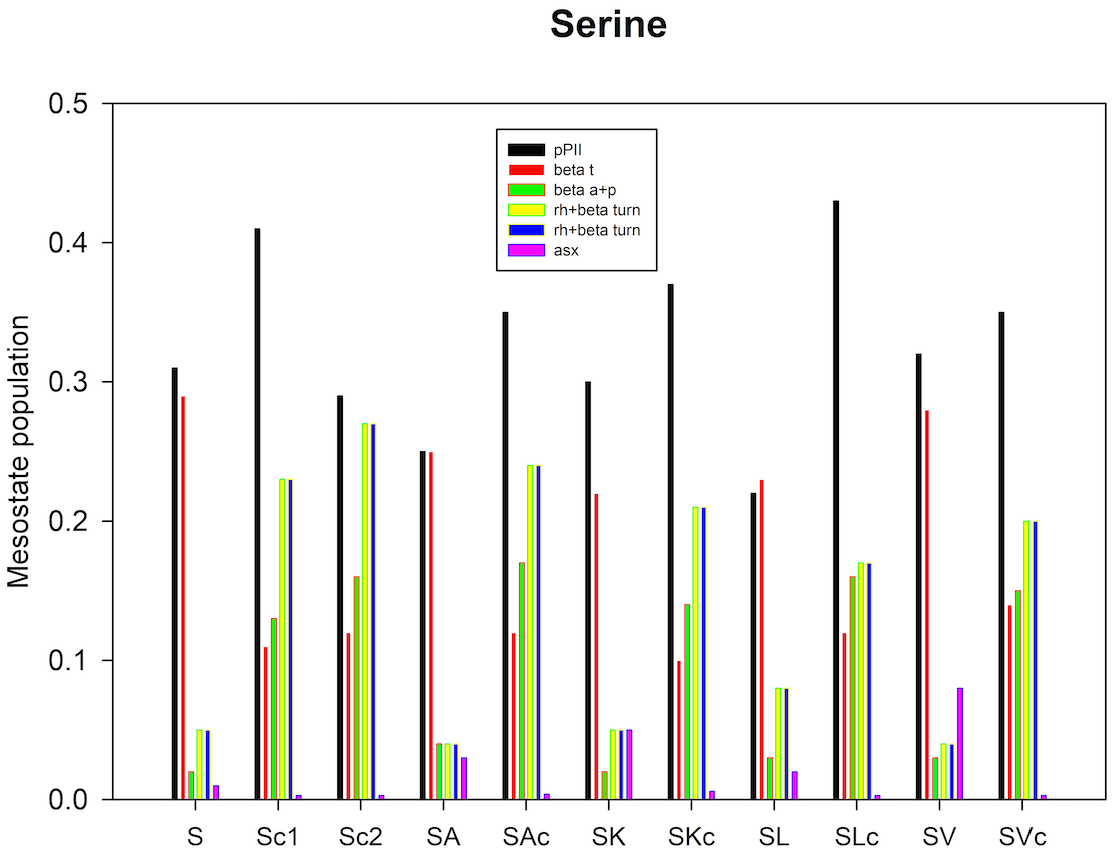

**Figure S18.** Comparison of ^3^J(H^N^H^Cα^) values of selected target residues (A,L and V) in GXYG (blue)(1) and blocked dipeptides with two amino acid residues (green).(17)

J-coupling [Hz]

|  | GXYG | XY-dipeptide |
| --- | --- | --- |
| Ag | 5.99 | 5.7 |
| Aa | 5.62 | 5.64 |
| Al | 6.58 | 5.92 |
| Av | 6.41 | 5.75 |
| Af | 6.08 | 5.89 |
| dA | 6.08 | 6.56 |
| sA | 6.48 | 6.34 |
| aA | 6.66 | 6.32 |
| fA | 5.89 | 6.14 |
| Lg | 6.69 | 7.3 |
| dL | 6.84 | 7.44 |
| sL | 7.22 | 7.32 |
| lL | 8.17 | 7.86 |
| aL | 7.47 | 7.42 |
| kL | 7.47 | 7.67 |
| vL | 7.27 | 7.22 |
| lL | 7.92 | 7.25 |
| Vg | 7.78 | 6.96 |
| dV | 8.03 | 7.78 |
| sV | 8 | 7.82 |
| aV | 7.21 | 7.75 |
| kV | 7.33 | 7.78 |
| Vl | 7.7 | 7.95 |

**Figure S19.** Probability density profile of serine in **all**-SL with respect to the φ-coordinate at ψ=150^o^. The plot was produced from files obtained from the website of the Dunbrack group with the kind permission of Dr. Dunbrack.(13)
